# Cryo-EM reveals patient- *versus* organ-specific structural diversity and bound ligands in λ6 light chain amyloids

**DOI:** 10.64898/2026.08.24.746819

**Authors:** Noorul Huda, Brian Spencer, Chad W. Hicks, Shobini Jayaraman, George A. Pantelopulos, Sherry Wong, Hui Chen, Robert B. Best, Vaishali Sanchorawala, Francesca Lavatelli, Tatiana Prokaeva, Olga Gursky

## Abstract

Immunoglobulin light chain (LC) amyloidosis is a debilitating multiorgan disease with limited treatment options. Sequence and structural variability make LC amyloids particularly challenging for therapeutic targeting. We report four cryo-EM structures of λ6-LC amyloid fibrils from four organs of two patients. Fibrils from different patients show different N-terminal conformations expanding known repertoire of λ6-LC amyloid folds. These folds contain a planar β-arch with a flexible linker containing the complementarity-determining region 2, flanked by N- and C-terminal segments in variable patient-specific conformations. The surface location of the structurally frustrated charged segment may contribute to the overrepresentation of the λ6-LC family in amyloidosis. These and other λ6-LC amyloid structures from different patients show different side chain packing. Conversely, cardiac, renal and splenic amyloids from the same patient exhibit similar structures with small peripheral organ-specific variations. Moreover, they show similar “orphan” densities, suggesting collagen-like triple helices bound to a tyrosine ladder along the fibril spine. Mass spectrometry detects collagen type-VI in tissue-extracted amyloids. Molecular dynamics simulations suggest amyloid binds collagen-VI triple helices via mixed interactions facilitated by the geometric complementarity between the layered amyloid structure and the triple helix. Similar interactions may drive formation of other amyloid-collagen complexes, influencing biological properties of amyloids.

## INTRODUCTION

Systemic amyloidoses are debilitating diseases wherein extracellular deposition of circulating proteins as amyloid fibrils in peripheral organs leads to organ damage and failure^1,2^. Since amyloid oligomers and mature fibrils contribute to the pathology in these diseases^3–5^, elucidating fibril structures and their interacting partners can inform the search for amyloid-targeting treatments. Cryogenic electron microscopy (cryo-EM) has revolutionized the field by determining near-atomic-resolution structures of amyloid fibrils from patients’ tissues^6,7^. Though distinct amyloid conformations, or strains, have been associated with subtypes of neurodegenerative disorders including Alzheimer’s, Parkinson’s and prion diseases^8–11^, no such association has emerged for systemic amyloidoses including their major form, immunoglobulin (Ig) light chain amyloidosis (AL). This life-threatening disease involves deposition of excess monoclonal Ig light chain (LC) in peripheral organs, particularly heart, kidney, liver and spleen^2,4,12,13^. Due to gene recombinations and somatic mutations^14^, each patient features a unique fibril-forming LC^15^ in a unique amyloid conformation^5,16–26^. This sequence and conformational variability is challenging for the structure-based therapeutic targeting. Another challenge is understanding how microenvironmental factors influence amyloid deposition in specific organs and leveraging it for therapeutic development.

LCs are 25 kDa proteins containing an N-terminal variable (V_L_) and C-terminal constant (C_L_) domains adopting native Ig β-sandwich folds (Fig. 1A, B). The fibril-forming V_L_ domains belong to two major isotypes, λ (overrepresented in AL repertoire) and κ, subdivided into families (λ1, λ3, λ6, etc.); of those, λ6 family, which contains V_L_ encoded by the *IGLV6-57* germline gene, is overrepresented in AL amyloidosis^15^. Cryo-EM structures of amyloid fibrils extracted from patients have revealed different folds for different λ-LCs^5,16–26^ but suggested structural similarities for amyloids derived from the same germline^21,22^. Furthermore, λ6-LC fibrils from the heart and kidney of the same patient showed near-identical structures^19^. Similarly, λ1-LC fibrils from the heart and fat of the same patient showed nearly identical structures with minor organ-specific differences at the fibril surface; conversely, different patients exhibited different amyloid structures^25^. These findings suggest that the dominant amyloid architecture is patient-specific rather than organ-specific. However, fibrils derived from patients with systemic amyloidosis including AL often show structural polymorphism^11,26,27^ that can be organ-specific^28^. Furthermore, extracellular factors including “amyloid signature” proteins can interact with pre-fibrillar and fibrillar LCs and influence fibril morphology^29^. Additionally, a cryo-EM study of cardiac and renal λ1-LC amyloids extracted from two different patients proposed that a particular surface charge distribution on the fibril end favors renal vs. cardiac deposition^24^. Therefore, organ-specific structural features in LC amyloids are far from clear.

**Figure 1.**
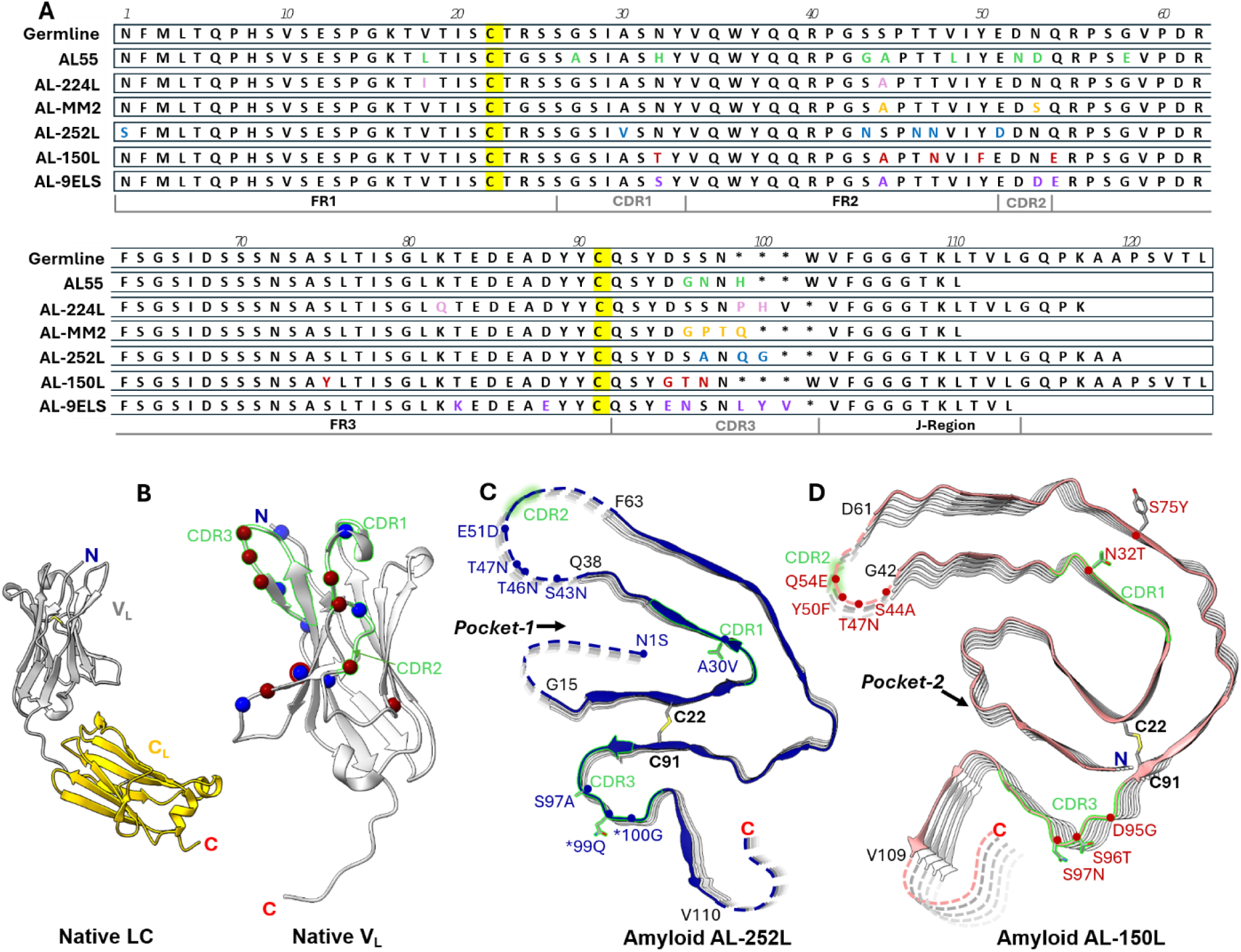
Structures of λ6-LC amyloid proteins. **A.** Amino acid sequences of six unique fibril-forming λ6-LCs whose amyloid structures have been determined by cryo-EM, and the corresponding germline sequence encoded by the *IGLV6-57* gene fragment. Proteins AL55^16^, AL-224L^22^, AL-252L and AL-150L (current study) are listed according to their AL-Base identifiers. AL-9ELS stands for the amyloid protein structure PDB ID: 9ELS^23^. AL-MM2 designates case 2 featuring both AL amyloidosis and multiple myeloma^25^. Colored letters indicate amino acids differences between each protein and its germline counterpart. C22 and C91 that form a conserved internal disulfide are highlighted. Framework (FR) and complementarity determining regions (CDR1, CDR2 and CDR3) in V_L_, along with the joined (J_L_) region linking V_L_ to C_L_, are indicated. Asterisks indicate sites of amino acid insertions. **B.** Native 3D structures of a full-length λ6-LC protein and its V_L_ domain (PDB ID: 6MG4). Amino acid mutations in AL-252L (blue dots) and AL-150L (red dots) differing from the germline sequence are mapped. **C, D**. Ribbon diagrams viewed down the fibril z-axis show one molecular layer of the cryo-EM structures of hepatic AL-252L amyloid (PDB: 35UN) and cardiac AL-150L amyloid (PDB ID: 13CN). Well-ordered amyloid core segments (solid lines), poorly ordered segments (dashed lines), their flanking residues (black letters) and the conserved internal disulfide, C22-C91, are shown. Blue (C) or red (D) letters indicate amino acid substitutions and insertion shown in panels A, B. In panels B-D green highlight marks CDR1, CDR2 and CDR3 segments. Pockets 1 and 2, which can accommodate N-terminal segments in different λ6-LC amyloids, are indicated.

To expand our understanding of LC amyloid folds, their patient-vs. organ-specific properties, and their natural ligands, here we report four new cryo-EM structures of λ6-LC amyloids from affected organs of two patients: liver (case AL-252, protein AL-252L) and heart, spleen and kidney (case AL-150, protein AL-150L). Though amyloids frequently deposit in the liver and spleen^30,31^, hepatic and splenic AL fibril structures have not been previously reported. Amyloids from different organs of AL-150 patient show near-identical structures with small variation around the periphery. Conversely, amyloids from different patients show major structural differences, including unexpected N-terminal conformations. Our results help elucidate key factors shaping the amyloid architecture and offer insights into amyloid interactions with collagen triple helices, with implications for other amyloids.

## RESULTS

### λ6-LCs form amyloid fibrils in AL-252 and AL-150 deposits

Amyloid fibrils were extracted from the unfixed post-mortem tissues of two patients. In AL-252 case^32^, a 55-year-old female with renal, hepatic and neurologic involvement passed away of spontaneous liver rupture following treatment with high-dose melphalan and autologous stem cell transplantation (Supplemental Table 1). Post-mortem analysis of all tissues explored (hepatic, renal, splenic and adrenal) showed unusually weak Congo red staining/birefringence by light/polarized microscopy (Supplemental Figure 1A, B). Immuno-electron microscopy showed λ-LC antibody reactivity (Supplemental Fig. 1C,D). The bone marrow-derived *IGL* gene sequence showed λ6-LC protein encoded by the germline gene fragments *IGLV6-57*01*, *IGLJ3*02*, and *IGLC3*04* (GenBank #MH996890). Liquid chromatography–tandem mass spectrometry (LC-MS/MS) of hepatic fibril extracts verified the identity of the bone-marrow-derived and amyloid-forming LC sequences (Supplemental Fig. 3C). 2D-PAGE of fibril extracts showed the most abundant species at 10-12 kDa (Supplemental Fig. 3A,B). LC-MS/MS of this species detected AL-252L fragments suggesting the fibrils contained V_L_ domain (Supplemental Fig. 3C).

In AL-150 case^33^, a 63-year-old male with soft tissue, cardiac and renal involvement passed away of heart failure (Supplemental Table 1). Post-mortem evaluation showed parenchymal and vascular amyloid deposits in multiple organs including heart, spleen and kidney, with strong Congo red staining/birefringence (Supplemental Fig. 2A-F). The bone-marrow-derived gene sequence showed λ6-LC protein encoded by the germline gene fragments *IGLV6-57*01*, *IGLJ3*02*, and *IGLC3*04* (GenBank #EF589390). LC-MS/MS of cardiac, splenic and renal fibril extracts established the sequence identity of the bone marrow-derived and amyloid-forming LCs (Supplemental Figs. 4C, 5C, 6C). 2D-PAGE of cardiac, splenic and renal fibril extracts showed the most abundant species at 10-12 kDa (Supplemental Figs. 4A,B, 5A,B, 6A,B). LC-MS/MS of this species detected AL-150L segments ranging from residues 1-118 to 1-153 (Supplemental Figs. 4-6, panels C).

Like in prior reports^22,33,34^, 2D Western blot of AL-252 and AL-150 amyloid tissue extracts detected full-length and fragmented LCs (Supplemental Figs. 3-6, panels A-B); LC-MS/MS identified additional amyloid signature and tissue-specific proteins (Supplemental Tables 2-5). Collagen type-VI (ColVI) was observed with similar abundance as in prior studies which reported ColVI-amyloid complexes^21,22^.

Cryo-EM structures of amyloid fibrils isolated from patients’ liver (AL-252L), heart, spleen, and kidney (AL-150L) were determined as described in Methods. Few other amyloid structures have been determined using data collected on a Glacios 2 microscope and refined using CryoSPARC software^22,35^. Supplemental Figures 7-11 and Supplemental Table 6 show the experimental workflow, the details of the data collection and analysis, and the model statistics. 2D classification detected no significant fibril polymorphism (Supplemental Figure 8) facilitating structural determination to 2.66-3.44 Å resolution (Table 1).

**Table 1.** Summary of cryo-EM structures of λ6-LCs amyloids. Amyloid proteins and organs from which they were extracted are listed. Proteins AL55^16^, AL-224L^22^, AL-252L and AL-150L (current study) are listed according to their AL-Base identifiers available via https://www.app.bumc.bu.edu/BEDAC_ALBase/Search. AL-9ELS stands for the amyloid protein structure with PDB ID: 9ELS^23^; AL-MM2 designates case-2 featuring both AL amyloidosis and multiple myeloma^26^. For each cryo-EM structure, parameters listed include the PDB ID, model resolution, N-terminal (NT) and C-terminal (CT) packing (described in the text), and stability estimates using PDBePisa^37^, https://www.ebi.ac.uk/pdbe/pisa/, and Atlas^38^, https://doi.org/10.5281/zenodo.15218932.

| Protein | Organ | PDB ID | Resolution (Å) | NT packing | Stability $\Delta G$ (kcal/mol) | |
| --- | --- | --- | --- | --- | --- | --- |
|  |  |  |  |  | PISA | ATLAS |
| AL55 | heart | 6HUD | 4.0 | Pocket-1 | -20.3 | -27.6 |
| AL55 | kidney | 8CPE | 4.0 | Pocket-1 | -23.3 | -30.2 |
| AL-224L | heart | 9OKA | 2.92 | Pocket-1 | -18.0 | -32.6 |
| AL-MM2 | fat | 9LJC | 2.70 | Pocket-1 | -22.1 | -32.0 |
| AL-252L | liver | 35UN | 3.44 | Flexible / Pocket-1 | -22.2 | -20.8 |
| AL-150L | heart | 13CN | 2.66 | Pocket-2 | -29.6 | -40.0 |
| AL-150L | spleen | 13FY | 2.79 | Pocket-2 | -31.1 | -34.1 |
| AL-150L | kidney | 35US | 3.40 | Pocket-2 | -29.4 | -36.2 |
| AL-9ELS | heart | 9ELS | 3.02 | Over CT surface | -19.7 | -28.0 |

### AL-252L hepatic amyloid shows a dynamic N-terminal conformation

AL-252L hepatic amyloid (3.44 Å resolution) contains a well-ordered core of 72 residues in two antiparallel segments, G15-Q38 (inner) and F63-V110 (outer) (Fig. 1C, Fig. 2). (Unless otherwise stated, we use continuous residue numbering for individual proteins; the numbering scheme of Fig. 1A is used only for comparing different LCs; in residues 1-98 these schemes coincide). Segment G15-Q38 forms a V-shaped pocket, termed pocket-1, encircled by segment F63-D95. The C-terminal tail (residues S96-V110) folds back upon the outer segment and makes a sharp turn at the conserved FGGG motif (residues 102-105). The inner and outer segments form a planar β-arch linked at opposite ends by the C22-C91 disulfide and by the poorly ordered central linker (residues Q39-R62) containing complementarity-determining region 2 (CDR2, residues 52-54) (Fig. 1C). The amyloid core is flanked by the N-terminal (residues S1-P14) and C-terminal segments (from K111 to approximately P118) that are poorly ordered and appear in the cryo-EM map at low contouring levels, forming a “fuzzy coat” (Fig. 2D, E, dashed lines).

**Figure 2.**
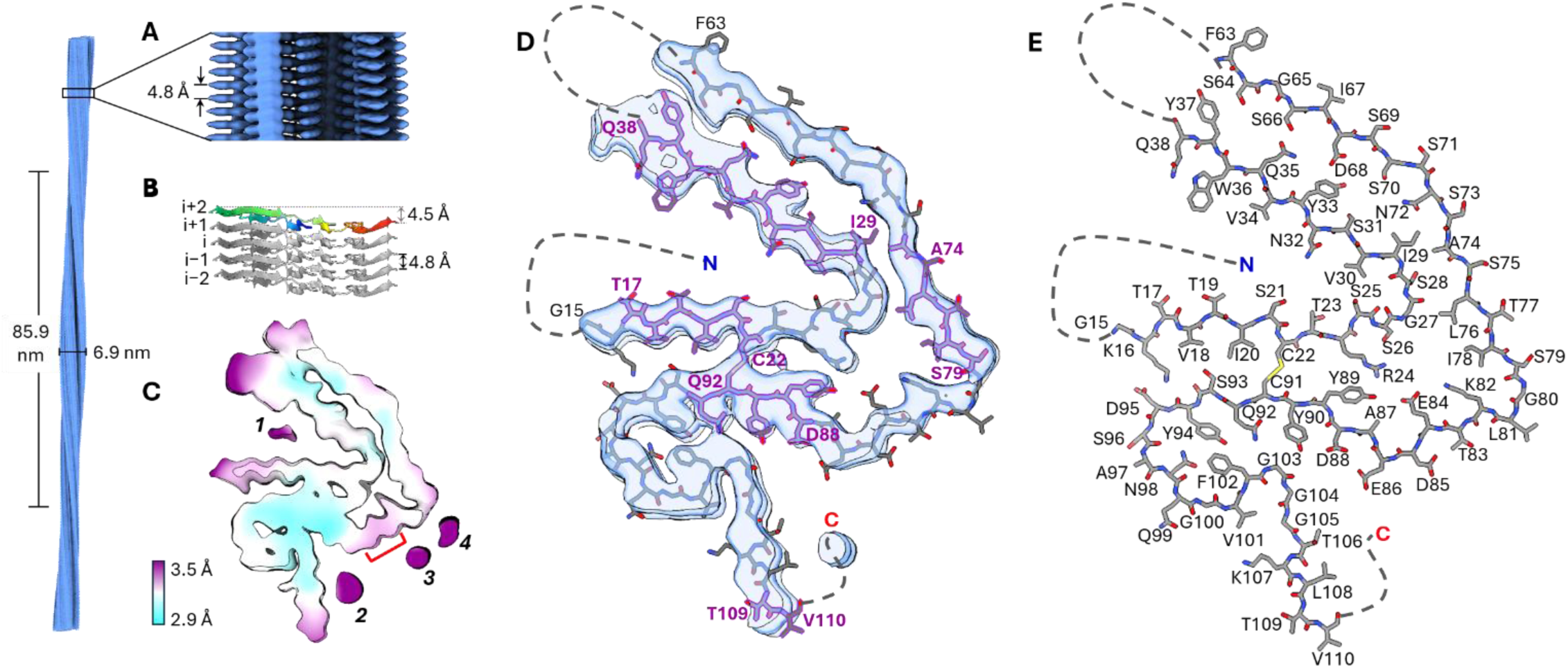
CryoEM reconstruction and atomic model of hepatic AL-252L amyloid fibril. **A.** Cryo-EM map of a fibril segment shows the crossover distance and the fibril width; a zoomed-in side view shows characteristic cross-β stacking with an inter-layer spacing. **B.** Five-layer atomic model of the fibril core (angled view). Top layer is rainbow-coloured from N-to C-terminus (blue to red) to illustrate planar molecular geometry. **C.** Cross-sectional view down the fibril z-axis of the reconstructed density coloured by local resolution. Extra densities are numbered 1 to 4. Red bracket marks the structurally frustrated acidic-rich segment, E84-D88. **D.** EM density and the atomic model of the amyloid core. Predicted amyloid-promoting regions and their flanking residues are in purple. Dashed lines indicate poorly ordered N-terminal, CDR2-containing linker, and C-terminal regions. **E.** Atomic model of one amyloid layer with residue assignments; dashed lines mark poorly ordered regions.

Several additional densities appear in the EM map at relatively low local resolution (Fig. 2C, purple). Density-1 in pocket-1 (Fig. 2C) merges with the β-arch inner segment at lower contouring levels, suggesting partially ordered N-terminal segment S1-L4 anchored in pocket-1. Similarly, density-2 merges with the well-ordered segment K107-V110 at lower contouring levels, suggesting partially ordered C-terminal segment anchored in a crevice on the fibril surface (Fig. 2D). External “orphan” density-3 and density-4 remain unassigned.

The major distinction between this and other available λ6-LC amyloid structures is that only AL-252L has a poorly ordered segment 1-14; also, only in AL-252L the entire amyloid core is nearly planar (Fig. 2B). In other structures, segment 1-14 is well-ordered and packs either inside pocket-1^16,22,26^ (Fig. 1D) or around the ordered C-terminal segment^23^ (Table 1). In AL-252L LC, N-terminal packing in pocket-1 may be affected by the N1S substitution. Additionally, A30V substitution in CDR1 in pocket-1, along with substitutions at other sites (nine in AL-252L compared to germline, Fig. 1A-C) potentially contribute to high flexibility of segment 1-14 in AL-252L. This flexibility complements this segments’ structural variability observed in other λ6-LC amyloids and shows that a well-ordered N-terminal conformation is not prerequisite for λ6-LC amyloid stability (Table 1).

### AL-150L cardiac amyloid shows N-terminus packed against the canonical disulfide

AL-150L cardiac amyloid (2.66 Å resolution) has a well-ordered 91-residue core containing antiparallel segments N1-G42 (inner) and D61-V109 (outer) (Fig. 1D, Fig. 3). The central linker in residues S43-P60 and the C-terminal segment T110-P118 are poorly ordered. Like in other λ6-LC amyloids, segment C22-C91 forms a nearly planar β-arch linked at opposite ends by the C22-C91 disulfide and the CDR2-containing linker spanning residues G42-P60. Well-ordered N- and C-terminal segments, N1-S21 and Q92-V109, protrude from the plane of the β-arch to form stabilizing cross-layer interactions, *i* to *i+1* and *i* to *i+2* (Fig. 3B).

**Figure 3.**
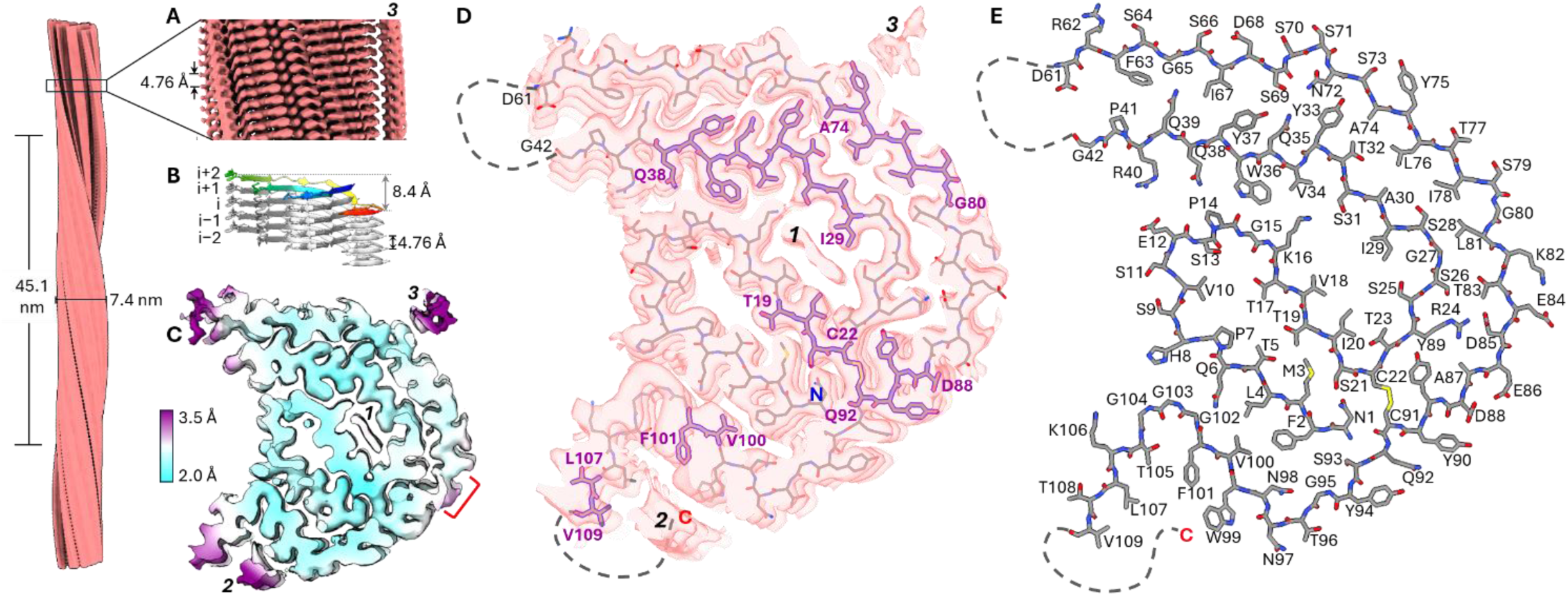
CryoEM reconstruction and atomic model of cardiac AL-150L amyloid fibril. **A.** Cryo-EM density of a fibril segment shows the crossover distance and the fibril width. A zoomed-in side view showing characteristic cross-β stacking with an inter-layer spacing. External density-3 is marked. **B.** Five-layer atomic model of the fibril core. Top layer is rainbow-coloured N-to C-terminus (blue to red) to emphasize the non-planar molecular geometry and the cross-layer interactions, i to i+2. **C.** Cross-sectional view down the fibril z-axis of the reconstructed density coloured by local resolution. Red bracket marks the structurally frustrated acidic-rich segment E84-D88. Numbers *1-3* indicate extra densities. **D.** EM density and the atomic model. Predicted amyloid-promoting regions and their flanking residues are in purple. Dashed lines indicate poorly ordered CDR2-containing linker and C-terminal regions. **E.** Atomic model of one fibril layer with residue assignments; dashed lines mark poorly ordered regions.

A major difference between this and other structures of λ6-LC amyloids is the N-terminal conformation. AL-150L shows residues N1-P14 packed against C91-G103 in a new inner pocket, termed pocket-2 (Figs. 1D, 4D), as opposed to pocket-1 (Figs. 1C, 4A) seen in most other λ6-LC amyloids^16,22,26^ (Table 1). Compared to these amyloids, the N-terminal segment in AL-150L undergoes a ∼180° backbone rotation around the P14-G15 junction, resembling that seen in AL-9ELS amyloid^23^ (see Discussion). Unlike AL-9ELS, in AL-150L amyloid segment 1-14 makes a turn near P7-H8 to pack in pocket-2, with N1 packed against the C22-C91 disulfide (Figs. 1D, 4D).

**Figure 4.**
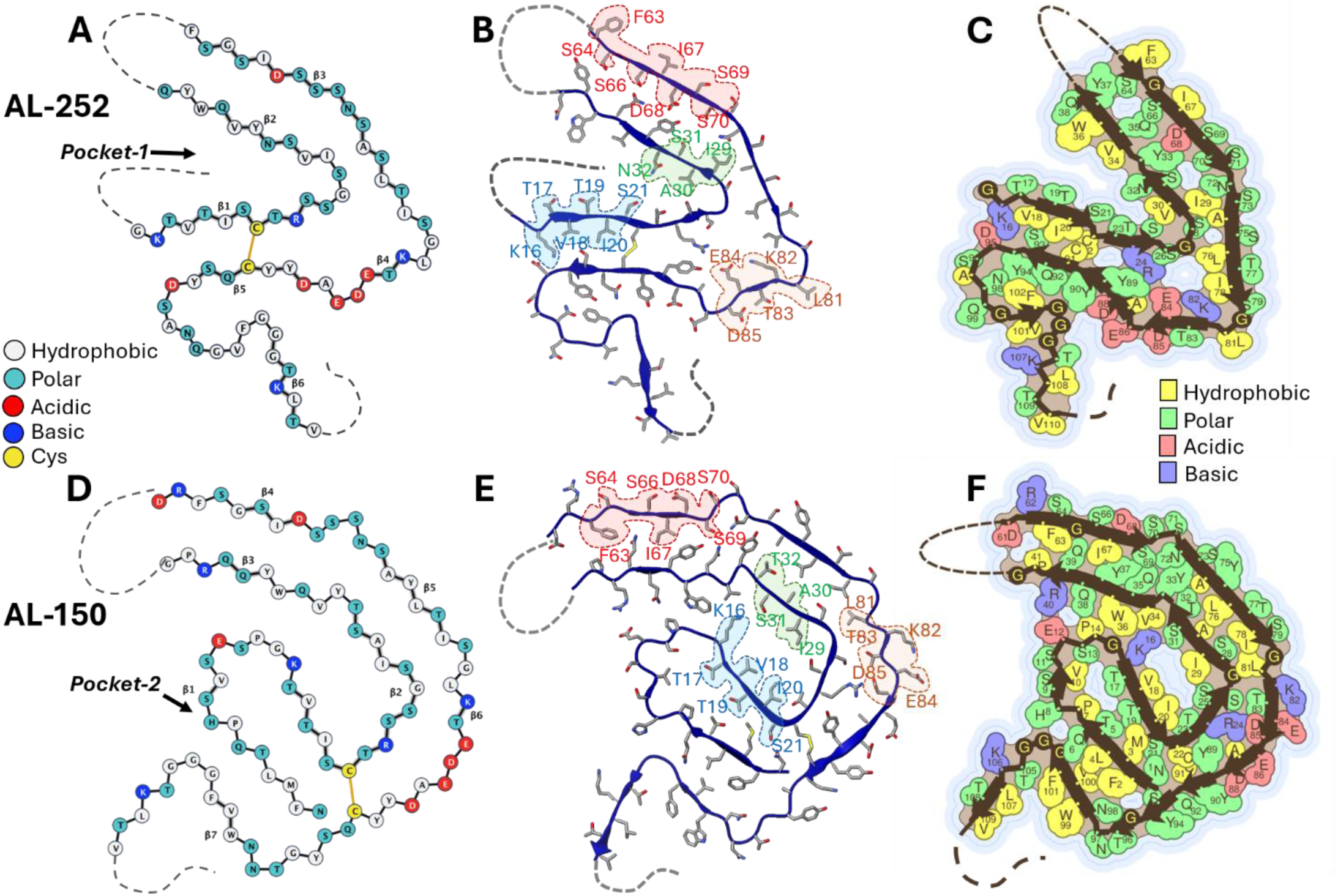
Backbone and side chain packing in AL-252L and AL-150L amyloid cores. Views down the z-axis show bead diagrams (**A, D**), segments with flipped side chain orientation in the two structures (**B, E**), and amino acid packing diagram (**C, F**) generated using Amyloid Illustrator^38^.

This unexpected conformation probably stems from some of the somatic mutations in AL-150L V_L_, five in the well-ordered core (N32T, S75Y, D95G, S96T, S97N) and four in the poorly ordered linker (S44A, T47N, Y50F, Q54E) (Fig. 1A,D); in this notation, the first and second residues represent AL and germline, respectively. While in AL-252L amyloid N32 side chain points inside pocket-1, in AL-150L amyloid T32 points outwards (Fig. 4B,E). Flipped orientation of this and other side chains (Fig. 4B,E) alters the lining of pocket-1, which is constricted to form an inner pore in AL-150L amyloid (Fig. 4E,F). Therefore, N32T substitution potentially contributes to the N-terminal displacement from pocket-1 in AL-150L amyloid. Additionally, the mutation-rich segment 95-97 from CDR3 extends away from C91 in AL-150L amyloid, creating a wide pocket-2 that accommodates the N-terminal segment (Fig. 1D, Fig. 4D-F). Substitutions such as D95G, which truncates the side chain, help form such a wide pocket-2. Additionally, D95G substitution eliminates the K16-D95 salt bridge that stabilizes the N-terminal packing in other λ6-LC amyloids^16,22,23^. Therefore, the new N-terminal conformation in AL-150L amyloid likely reflects combined effects of several mutations including N32T and D95G.

Prior studies of native recombinant AL-150L V_L_ protein have demonstrated the pro-amyloidogenic effect of N32T substitution^36^. Hydrogen-deuterium exchange mass spectrometry and stability studies showed N32T substitution destabilizes the native structure in V_L_, disrupts the local hydrogen-bonded network, increases the amyloid-forming sequence propensity, and accelerates amyloid formation *in vitro*^36^. Current results provide complementary information on how AL-150L mutations including N32T influence patient’s amyloid structure.

### AL-150L cardiac, splenic and renal amyloids show similar structures

AL-150L amyloids from the heart, spleen and kidney have very similar structures (Fig. 5A-C). Notably, PDBePISA and ATLAS algorithms^37,38^ predict these amyloids to be more stable than any other known λ6-LC amyloid structures (Table 1). Pairwise comparison of cardiac vs. splenic or renal amyloids using Chimera^39^ yields root-mean-square deviation for 91 common C_α_ atoms of only RMSD=0.6 Å and the Amyloid Packing Difference (APD), a metric to compare side chain packing in similar amyloid proteins^40^, of 1-3% (Supplemental Fig. 12), indicating nearly identical structures.

**Figure 5.**
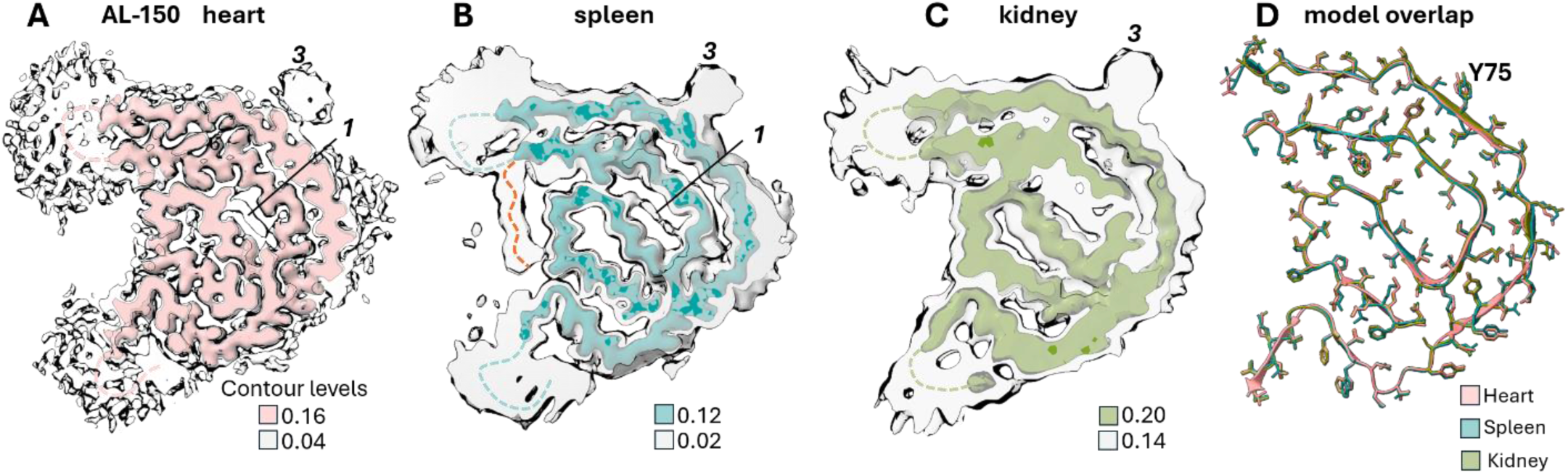
CryoEM reconstruction of cardiac, splenic and renal AL-150L amyloid fibrils. **A-C.** Cross-sectional maps at different contouring levels are viewed down the fibril z-axis. Dashed lines mark protein backbone in poorly ordered / low-occupancy regions, which are seen at low contouring levels (gray). Dashed orange line marks the area observed only in the splenic amyloid map at low contouring levels. Extra densities 1 (internal) and 3 (external) are indicated. **D**. Atomic model of AL-150L amyloids from the heart, spleen and kidney overlapped (Cα RMSD=0.6 Å).

The only significant difference is in the CDR2-containing linker G42-P60. At lower map contouring levels, splenic amyloid shows an additional well-defined density that extends from S43 to approximately F50 and packs against the well-ordered residues H8-E12 (Fig. 5B). Considering high proteolytic activity in the spleen, this finding suggests that a subset of LC molecules is cleaved in the flexible central linker near F50, liberating segment 43-50 to pack against the amyloid core. Consistent with this idea, AL-150L residues N47-E54, NVIFEDNE, contain predicted cleavage sites near F50 for several proteases that are active in the spleen, including cathepsins B, D (lysosomal proteases abundant in macrophages) and G (a serine protease)^41–44^. These cathepsins are detected by LC-MS/MS at low abundance in splenic amyloid tissue extracts (Supplemental Table 4). Like in type-A transthyretin fibrils^45^, proteolytic cut in the middle of some fibril-forming V_L_ molecules in splenic amyloid probably occurs after amyloid formation.

### Accommodation of the structurally frustrated EDEAD motif in λ6-LC amyloids

IgG λ-LCs contain a conserved acidic-rich motif in residues E84-D88 located on the fibril surface of all known λ6-LC amyloid structures. Unbalanced charges in such motifs are strongly disfavored due to electrostatic repulsion within and between identical molecular layers^6,46,47^. These conflicting interactions cause structural frustration exemplified by the poorly ordered EDEAD segment in AL-224L amyloid^22^. Like AL-224L, cryo-EM maps of AL-252L and AL-150L amyloids show reduced local resolution indicating increased dynamics in segment E84-D88 (Figs. 2C, 3C). Additionally, structure-based analysis using Amyloid Illustrator^38^ predicts highly unfavorable free energy contributions to amyloid stability from these acidic side chains, particularly D88 (Supplemental Fig. 13). Furthermore, the Protein Frustratometer tool^48^ predicts high structural frustration in D85, E86 and D88 for all available λ6-LC amyloid structures including AL-252L and AL-150L (Supplemental Fig. 14).

In the cryo-EM maps of AL-252L and AL-150L amyloids, the acidic-rich segment is well-resolved, suggesting a partial charge balance, and the cryo-EM structures suggest local stabilizing interactions (Fig. 6). In AL-252L amyloid, the inward-pointing E84 and K82 side chains form intra-layer or cross-layer ion pairs (Fig. 6B). E84 can also form intralayer or cross-layer ion pairs with R24, while R24 forms cross-layer hydrogen bonds with Tyr89 OH (Fig. 6B). Charges on the outward-facing D85, E86 and D88 side chains are screened by solvent (Fig. 6A). Moreover, 3.2 Å spacing between E86 and D88 side chains suggests carboxyl pairing (Fig. 6C), i.e. carboxyl protonation at an anomalously high pK_a_ facilitating proton sharing between adjacent carboxyls to eliminate their electrostatic repulsion and stabilize the structure^46,49^. This conjecture is supported by anomalously high pK_a_ predicted in AL-252L amyloid for these side chains, particularly D88 (Supplemental Table 7), using PROPKA-3 server^50^.

**Figure 6.**
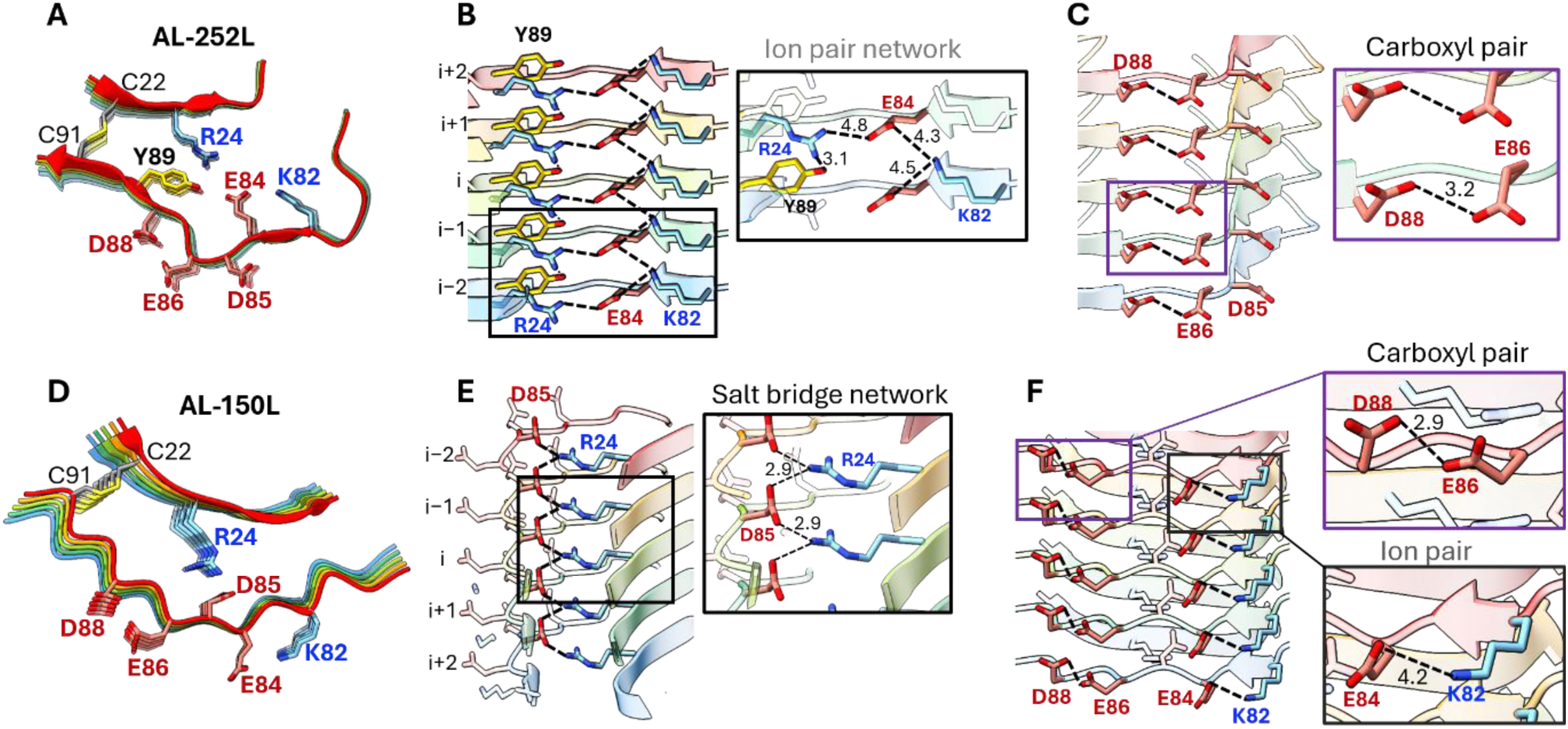
Stabilizing interactions in the structures of AL-252L and AL-150L amyloids involving the acidic-rich segment E84-D88. Top (**A, D**) and side views (**B, C** and **E, F**) of hepatic AL-252L (**A-C**) and cardiac AL-150L (**D-E**) amyloids show key side chains and their interactions (dashed lines). Interatomic spacings in Å are indicated for ion pairs (≤5 Å), salt bridges (≤4 Å), hydrogen bonds and carboxyl pairs (2.8-3.2 Å). Individual protein molecules are colored.

Stabilizing interactions in the EDEAD segment are also evident in AL-150L amyloid (Fig. 6D-F). Outward-pointing E84 and K82 side chains form an ion pair (Fig. 6F). Each inward-pointing D85 carboxylate forms salt bridges with two R24 guanidino groups from two adjacent layers, while each R24 forms salt bridges with two D85 carboxylates from adjacent layers, forming a stabilizing cross-layer network of 2.9 Å salt bridges along the fibril z-axis (Fig. 6E). Such a buried salt bridge network probably contributes to increased stability predicted for AL-150L compared to other λ6-LC amyloids (Table 1). Furthermore, solvent-facing E86 and D88 side chains spaced at 2.9 Å likely form a carboxyl pair (Fig. 6F) predicted by PROPKA-3 (Supplemental Table 7). In summary, salt bridge / ion pair networks, carboxyl pairing, and solvent screening of the acidic side chains on the fibril surface help partially balance their negative charges which, together with increased structural flexibility suggested by the decreased local resolution in the cryo-EM maps (Figs. 2C, 3C), helps accommodate the EDEAD motif in λ6-LC amyloids.

### Differential binding of Congo red to AL-252L and AL-150L amyloids

Birefringence upon tissue staining with Congo red is a histological gold standard in amyloidosis diagnostics^51^. Unusual cases such as AL-252 are clinically intriguing because weak tissue staining by Congo red despite amyloid deposition (Supplemental Fig. 1A,B) could have diagnostic implications^52^. To recapitulate this effect in solution, we monitored Congo red binding to tissue-extracted amyloids by tracking red shifts in the dye’s absorption spectra. As in tissues, AL-252L amyloid showed weak dye binding while AL-150L amyloid showed strong binding in solution (Supplemental Fig. 15). To unveil structural underpinnings, we leveraged the cryo-EM structure of transthyretin amyloid with bound Congo red, which preferentially binds in an amphipathic groove formed by well-ordered Arg ladders along the fibril surface^51^. A well-ordered R62 is present on the amyloid surface of AL150L but is absent from AL-252L (Fig. 4). Though AL-252L features surface R55 in the disordered linker, this site is not expected to strongly bind and immobilize the dye in a rigid conformation that is necessary for strong birefringence. The absence of well-ordered Arg on the AL-252L amyloid surface provides a structural hypothesis for the observed staining phenotype. Nevertheless, Congo red can potentially bind at other surface sites^51^ including Lys arrays^53^. Both amyloid structures contain such arrays (K107 in AL-252L, K88 and K106 in AL-150L, Fig. 4) potentially contributing to weak dye binding observed in AL-252L amyloid.

### Cardiac, splenic and renal AL-150L amyloids show similar extra densities

Like most other tissue-derived amyloids^22,23,25,46^, the cryo-EM maps of LC amyloids contain strong “orphan” densities that cannot be attributed to the fibril-forming protein (Figs. 2C, 3C). Cardiac and splenic AL-150L amyloids show not only similar structures but also similar extra densities: an elongated density-1 in the internal pore and a large external density-3 near Y75 (Fig. 5A, B). Despite lower resolution of renal amyloid, its cryo-EM map also shows a large external density-3 near Y75 (Fig. 5C). These independent observations exclude the possibility of noise and suggest binding of similar ligands to AL-150L amyloids in different organs. To help identify these ligands, we used the highest-resolution map and atomic model of cardiac amyloid.

Density-1 in the inner pore is elongated in the x-y plane (Fig. 5A, B) and discontinuous in the z-direction. This pore, which forms upon constriction of pocket-1 in AL-150L amyloid, is lined by K16, V18, I20, T23, S25, I29, and S31 side chains (Fig. 4D-F). We conjecture that an unidentified small amphipathic inclusion co-assembles with AL-150L protein, stabilizes the amyloid pore, and helps alleviate the destabilizing effect and high frustration predicted for K16 (Supplemental Figs. 13B, 14B). Small hydrophobic or amphipathic inclusions have also been observed in internal pores of other amyloids^6,54^ including LCs^5,22^. These inclusions are probably incorporated during amyloid formation potentially acting as amyloid cofactors^6,54^.

Notably, the external density-3 near the outward-facing Y75 array is hollow in the x-y plane, continuous along the fibril z-axis, and is centered ∼15 Å away from the nearest LC atoms (Fig. 3A, C; Fig. 5), a geometry consistent with a collagen triple helix^55–57^. This density is not observed near S75 at the surface of AL-252L (Fig. 2) or any other λ6-LC amyloids and is probably related to the S75Y substitution in AL-150L protein. Several other LC structures show external densities near exposed aromatics, which are consistent with collagenous triple helices^22,25^.

LC-MS/MS analyses of cardiac, splenic and renal AL-150L amyloid tissue fractions detected ColVI chains α1, α2 and α3 (Supplemental Tables 3-5). ColVI chains were found with similar abundance in prior studies of cardiac AL59 λ3-LC and AL-224L λ6-LC fibrils, which were extracted using similar protocols and contained ColVI bound to LC fibrils^21^ or were strongly implicated to contain it^22^. Therefore, the orphan density near Y75 potentially represents ColVI triple helix. Triple-helical domains of other proteins cannot be excluded, e.g. collagen type-I (which was most abundant in renal amyloid) and complement C1q (which was detected by LC-MS/MS in all AL-150L samples at lower abundance). Since ColVI was among the most abundant proteins in cardiac, splenic and renal AL-150 amyloid extracts, as well as in other studies of cardiac AL amyloids^21,22,25^ and transthyretin amyloids^58^, ColVI was selected for further analysis.

### Molecular dynamics simulations of amyloid in complex with ColVI triple helix

Compared to fibril-forming collagens such as type-I, ColVI triple helices (335/336 residues per chain) are enriched in charged and aromatic residues^59^. In natural collagens with low imino acid content, triple helices are thought to exhibit a 10/3 structure featuring ten GXY repeat units per helical turn (where X is often a proline and Y is an hydroxyproline), a helical twist of 36° per unit, and a pitch of ∼28.6 Å^56,60^. This pitch matches the ∼28.8 Å thickness of 6 amyloid layers with ∼4.8 Å interlayer spacing. This geometric complementarity helps explain excellent agreement between the extra density seen in the cryo-EM map of cardiac AL-150L amyloid, which was reconstructed using amyloid helical symmetry, and the collagen triple helix (Fig. 7A,B).

**Figure 7.**
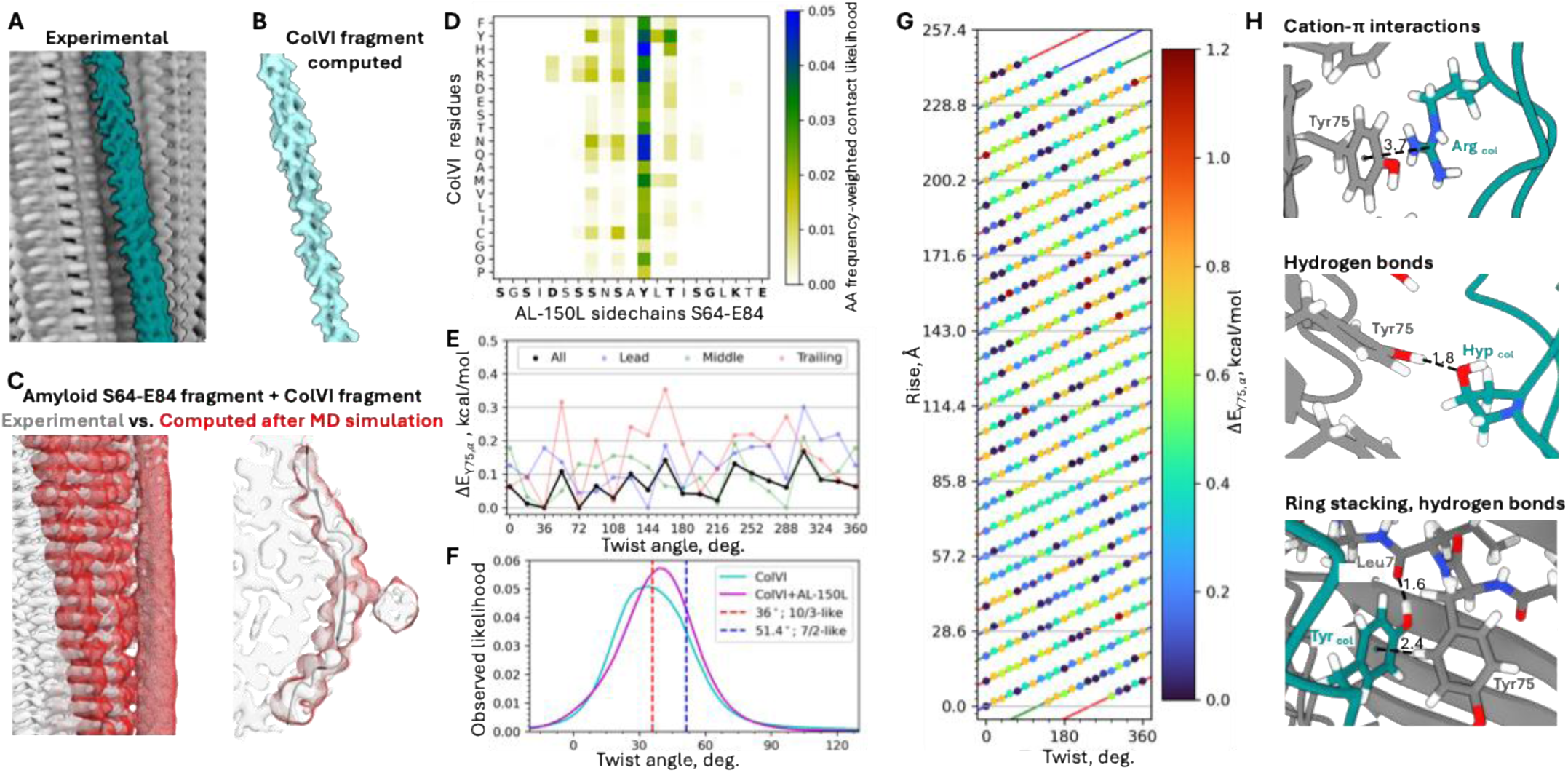
Molecular modeling of amyloid interactions with ColVI triple helix. **A.** Cryo-EM map of cardiac AL-150L amyloid (grey) shows extra density near Y75 (teal). **B.** Computed map of a triple-helical ColVI fragment #1 (configuration 1 after MD simulations described in supplemental Methods). **C.** Experimental density determined by cryo-EM (grey) overlaps the computed density (red mesh) after MD simulations of a complex containing AL-150L amyloid residues S64-E84 and eight ColVI triple-helical fragments. **D.** Contact probability between AL-150L amyloid residues S64-E84 (x-axis) and ColVI residues plotted by residue type (y-axis), weighted for ColVI residue frequency. Contacts between non-hydrogen atoms of ColVI and amyloid within a 3.5 Å cutoff, *p_i,a_*, were determined in MD simulations. **E.** Distribution of triple-helical twist angles in absence (cyan) and presence of AL-150L amyloid (magenta) determined in MD simulations. 0°, 36°, and 51.4° lines show values for the untwisted, 10/3, and 7/2 triple-helix models. **F.** Relative interaction free energy, DE, for each face defined by 20 Cγ angular positions of *X*- and *Y*-amino acids in a 10/3 triple helix. In panels F and G, positive values represent less favorable energy as compared to ColVI Tyr, which interacts most favorably with amyloid (DE=0). **G.** Helical net of the full ColVI triple-helical domain in a 10/3 model^60^ depicts relative free energy of amino acid side chain interaction with Y75 ladder on amyloid, ΔE*_Y75,α_*. Dots represent side chains colored according to interaction energy, from 0 (most favorable, navy) to red (least favorable, dark red). Twist and rise values correspond to Cγ positions for the lead (blue), middle (green), and trailing (red) alpha chains (see Supplement for details). **H.** Representative residue contacts between AL-150L amyloid and ColVI triple helix observed in MD trajectories; distances are in Å.

To evaluate amyloid-collagen interactions in atomic details, we performed molecular dynamics (MD) simulations using eight 30-to 42-residue fragments of the ColVI triple-helical domain as described in Supplemental Methods. As starting models, we used four alternative orientations of each fragment relative to the Y75 amyloid ladder (Supplemental Fig. 16). During 1 μs simulations, ColVI triple helices freely diffused about the Y75 ladder (Supplemental Fig. 17), sampling electron density that closely superimposed the extra density near Y75 in the cryo-EM map (Fig. 7C). Charged, aromatic, and polar residues of ColVI, weighted for amino acid frequency, exhibited the highest propensity to interact with Y75 ladder (Fig. 7D). Unweighted contact frequencies suggested that charged residues, Gly and Hyp are principally responsible for ColVI-amyloid interactions (Supplemental Fig. 18). All residues interacted with the Y75 ladder via π-rings, hydrogen bonding, or charge interactions; they also interacted weakly with neighboring residues (Fig. 7D,E,F). Of all ColVI side chains, Tyr had the greatest propensity to interact with amyloid via π-stacking with Y75 and hydrogen bonding to L76 backbone (Fig. 7D,H). ColVI charged and polar side chains formed multiple contacts with up to 87 ns-long average lifetimes on amyloid (Supplemental Fig. 19). The average ColVI triple-helical twist was 36.4° in the absence and 38.0° in the presence of amyloid, suggesting slightly tighter triple-helical winding to complement amyloid structure (Fig. 7F; Supplemental Fig. 20A). Scoring the interaction of Y75 with the *X*- and *Y*-angular positions of the ColVI triple-helical domain using a mean field contact potential indicates that the 20 unique helical faces show no strong preference for the Tyr ladder, suggesting no dominant rotational orientation of the triple helix relative to amyloid (Fig. 7E,G Supplemental Figs. 20B, 21).

In summary, local complementarity between the triple helix and the periodic amyloid structure helps maximize their mixed lateral interactions to form a stable complex. We conjecture that similar interactions may stabilize amyloid-ColVI complexes described in other studies at lower resolution^21,22,25,58^.

## DISCUSSION

This study reports four λ6-LC amyloid structures from four affected organs of two clinical cases, AL-252L and AL-150L. These structures, taken together with cryo-EM fibril structures of four previously reported cases^16,22,23,26^ (Table 1, Fig. 8), expand the conformational repertoire in this amyloid protein family and reveal shared, patient-specific and organ-specific features. These structures suggest strongly that the patient-specific LC sequence determines the fibril architecture, while organ-specific factors induce modest peripheral changes.

**Figure 8.**
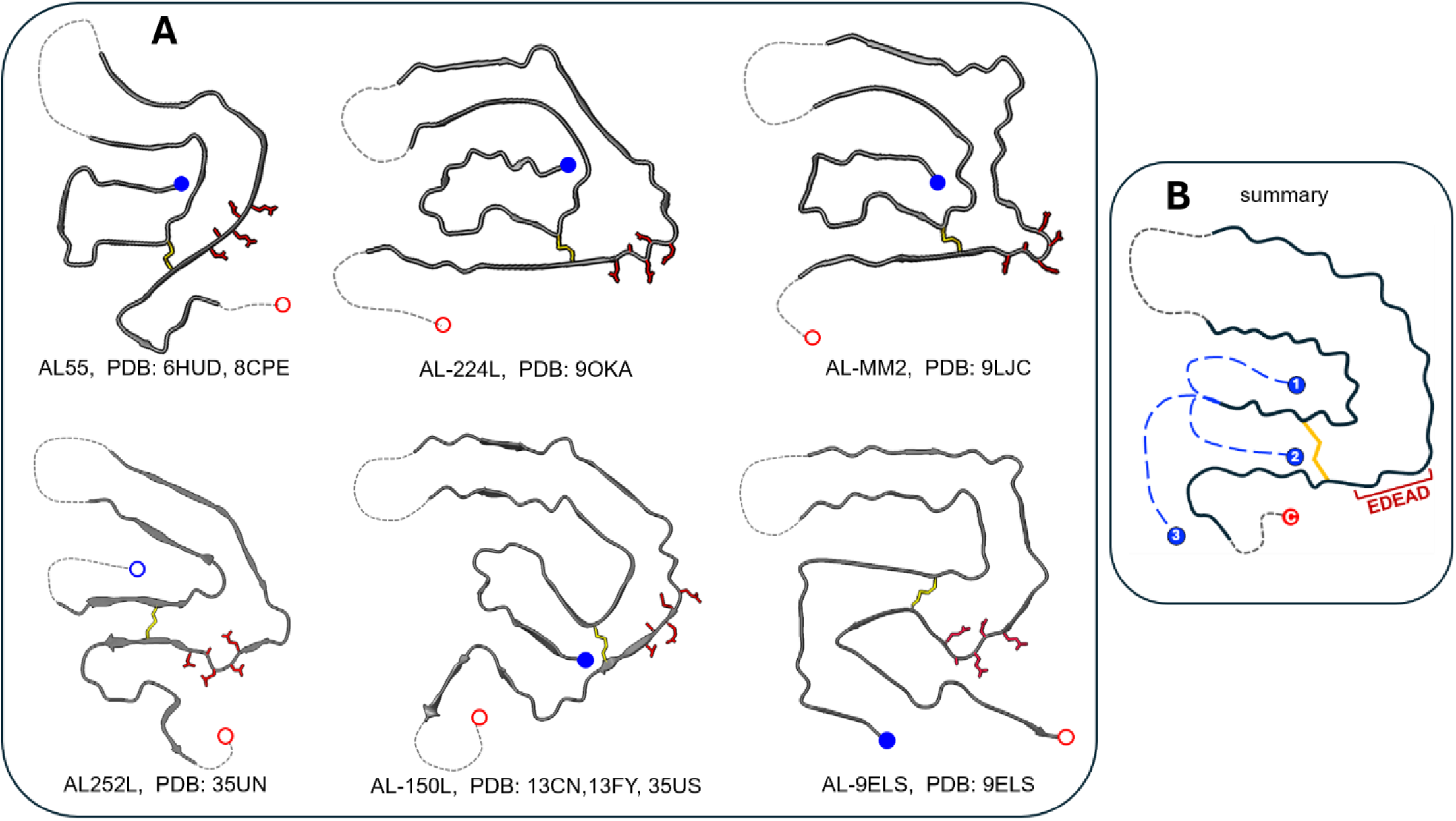
Conformational repertoire in human tissue-derived λ6-LC amyloids. **A.** Ribbon diagrams show amyloid folds for the nine known cryo-EM fibril structures of λ6-LCs from six different patients. Amyloid cores are in solid lines; poorly ordered segments are in dotted lines. N- and C-termini are marked by blue and red dots that are either solid (well-ordered) or open (poorly ordered). Red stick models show side chains in the acidic-rich EDEAD segment. **B.** Scheme showing alternative conformations of the N-terminal segment in λ6-LC amyloids.

All known λ6-LC structures contain a nearly planar ∼70-residue β-arch disulfide-linked at C22-C91, which folds back at the poorly ordered central linker containing hypervariable CDR2 (residues 53-55). In these structures the 1^st^ β-arch segment spans from C22 to approximately F50, which is shorter than the 2^nd^ segment spanning from approximately E51 to C91. To accommodate this difference in length, the 1^st^ segment packs in the interior and is encircled by the 2^nd^ segment around the fibril surface. This surface location facilitates solvent exposure of the conserved acidic-rich motif in residues E84-D88 (Fig. 8). To accommodate this energetically unfavored motif in amyloid, its excess charges must be balanced, at least in part, via salt bridges/ion pairing, carboxyl pairing, solvent screening (Fig. 6) and increased local dynamics indicated by reduced local resolution (Figs. 2C, 3C). Surface location of the EDEAD motif in λ6-LC fibrils facilitates solvent screening and increased local dynamics. In contrast, this acidic motif is sequestered in most available structures of λ1-LC and λ3-LC amyloids (Supplemental Fig. 22). We hypothesize that surface location of this structurally frustrated motif diminishes its gate-keeping potential in amyloid formation^22^ and facilitates its accommodation in λ6-LC fibrils, which potentially contributes to the overrepresentation of the λ6-LC protein family in AL amyloidosis.

The β-arch in λ6-LC amyloids is flanked by the N- and C-terminal segments in variable conformations. This conformational variability stems, in part, from mutations in two hypervariable regions, CDR1 and CDR3 (Fig. 1B-D). In three out of six unique λ6-LC amyloid structures (Table 1, Fig. 8), segment 1-14 packs in pocket-1^16,22,26^. Alternatively, segment 1-14 can be poorly ordered (AL-252L), pack in pocket-2 (AL-150L), or encircle the C-terminal segment at the fibril surface (AL-9ELS) (Fig. 8). Since residues 1-14 are identical in these six λ6-LCs (except for N1S substitution in AL-252L, Fig. 1A), their conformational variability must reflect amino acid differences elsewhere in the protein. A case in point is AL-150L substitutions N32T in CDR1 and D95G in CDR3, which destabilize the N-terminal packing in pocket-1 and help accommodate it in pocket-2 (Fig. 1D). This exemplifies how concerted effects of several mutations from different molecular regions contribute to AL amyloidosis by stabilizing a biologically fit amyloid conformation.

Though the β-arch has emerged as a shared structural motif in λ6-LC amyloids, it shows inverse side chain orientation (flipping) and registry shifts in amyloids from different patients. Comparison of AL-252L and AL-150L amyloids shows flipped side chain orientation in four segments from the β-arch and one C-terminal segment (Fig. 4B, E). Additionally, β-arch segments show registry shifts; e.g. in AL-252L amyloid, Y33 side chain packs against D68 and S70 side chains, while in AL-150L amyloid Y33 packs between N72 and A74, showing a 4-residue shift in registry (Fig. 4). As a result, AL-252L and AL-150L amyloids show major packing differences (Fig. 4A, B): the difference in side chain contacts for 71 common amyloid core residues is APD=87% (Supplemental Fig. 12A, B). Similarly, pairwise comparison of other λ6-LC amyloid structures from different patients shows APD=75-95% (Supplemental Fig. 12A). Side chain flipping and backbone registry shifts have been observed in other amyloids with similar but not identical primary^20,40^ or 3D structures^25^ and were proposed to provide a general mechanism for adaptation of amyloid fold to small side chain differences^22^. Current study supports this mechanism; it reflects packing versatility of the cross-β conformation in pathologic amyloids, as opposed to packing specificity in native protein structures.

In stark contrast to amyloids from different patients, amyloids from different organs of AL-150 case show near-identical structures (Fig. 5, Supplemental Fig. 12). Minor differences detected at the fibril surface reflect organ-specific microenvironments, such as the high proteolytic activity in the spleen (Fig. 5B). This finding, together with other structural studies of amyloids from different organs of the same patient^19,26^, suggest that amyloid architecture is shaped primarily by the patient-specific factors, while the organ-specific factors cause small if any structural changes at the fibril periphery.

Collectively, findings of similar LC amyloid structures in different organs of the same patient^19,26^, including heart, spleen, liver, kidney and fat (Fig. 5, Supplemental Fig. 12A), provide a challenge for a recent study attributing large structural differences between cardiac and renal LC amyloids from different patients to organ-specific microenvironment^24^. Notably, similar fibril structures in different organs are consistent with amyloid propagation in the body via seeding. Such a template-based mechanism was explored in AL LC fibrillation studies in vitro^61^ and implicated for other systemic amyloidoses in vivo^62–64^.

Taken together, known λ-LC fibril structures from different patients suggest that a major determinant for the amyloid architecture is the amino acid sequence of the fibril-forming protein encoded by the germline progenitor and modulated by somatic mutations. Environmental factors, either shared or organ-specific, can importantly influence amyloid deposition and properties. The nature of such factors is emerging from reports that ColVI, the major collagen in the pericellular space of the extracellular matrix, can form superhelical complexes with amyloids in tissues^21,58^. Like several recent studies, the current LC-MS/MS analysis of cardiac, renal and splenic AL-150L amyloids detects all three ColVI chains at relatively high abundance (Supplemental Tables 3-5). Cryo-EM maps of these and other amyloids contain orphan densities coordinated by aromatic ladders on the fibril surface, which are conjectured to represent collagen-like triple helices^22,25^. Current study offers detailed insights into amyloid–triple helix interactions. Approximate complementarity between the periodic structures of amyloid and the 10/3 triple helix enables us to visualize collagen in unprecedented detail in the 2.66 Å cryo-EM map reconstructed using amyloid helical symmetry (Fig. 7A). This complementarity facilitates extensive mixed interactions, including π-stacking and hydrogen bonding, involving mainly Y75 ladders on amyloid and various ColVI side chains (Fig. 7D,H). These interactions potentially help protect bound triple helices from the proteolytic degradation in vivo and from the processive cleavage by collagenase^65^ during fibril isolation in vitro. Reciprocally, bound collagen probably influences amyloid properties, including amyloid recognition and clearance by macrophages^66^, amyloid proliferation via fibril fragmentation or secondary nucleation^61^, or other amyloid interactions. Since periodic structure with ∼4.8 Å spacing is shared by different amyloids, its geometric complementarity to the triple helix is probably relevant to other amyloid-triple helix interactions.

## STUDY LIMITATIONS

Cryo-EM, which is the method of choice for structural studies of tissue-extracted amyloids, has limitations. Firstly, tissue-extracted amyloid fibrils including AL LCs are often polymorphic^11,22,27^, yet most cryo-EM studies of LCs resolve the structure of one dominant polymorph. Recently reported structures of AL-9ELS amyloid polymorphs show a conserved amyloid conformation and differ in the number of protofilaments^23,28^. Whether the rule “one AL patient – one amyloid fold” extends to other amyloid polymorphs will be determined in future cryo-EM studies.

Secondly, cryo-EM alone is insufficient to unambiguously identify bound ligands in tissue-derived amyloids. Unless the ligand occupancy is high and the symmetry matches that of amyloid, the ligand density is blurred or undetectable in the EM maps obtained by helical reconstruction using amyloid symmetry^53^. This warrants the use of complementary approaches, each with its own limitations. LC-MS/MS is a semi-quantitative method that detects proteins co-isolated with amyloids but does not prove their direct binding to amyloid. Immunocapture-based methods to probe the biochemical composition of the amyloid-ligand complexes^22^ are limited by epitope accessibility. Computational approaches can test if a ligand potentially forms a stable complex with amyloid, yet do not prove it. Additional insights can be obtained from comparative studies of recombinant fibrils formed with or without a ligand, although direct comparison with their tissue-derived counterparts may be hindered if different fibril structures form in vitro and in vivo.

Lastly, even a mild water-based procedure of amyloid isolation from tissues for cryo-EM analysis removes physiologically relevant components including collagens, thereby limiting our understanding of amyloid-ligand interactions in situ. Cryogenic electron tomography overcomes this limitation but has low resolution^21,58^. Therefore, identification of bound ligands in amyloids remains challenging.

## Abbreviations

AL: amyloid light chain;
APD: amyloid packing difference;
C_L_: light chain constant domain;
cryo-EM: cryogenic electron microscopy;
Ig: immunoglobulin;
LC: light chain;
LC-MS/MS: liquid chromatography–tandem mass spectrometry;
MD: molecular dynamics;
RMSD: root mean square deviation;
V_L_: light chain variable domain.

## Ethical statement

This study was conducted in accordance with the Declaration of Helsinki. Informed consent for the sample and data collection was obtained from the patients with the approval of the Institutional Review Board at the Boston Medical Center.

## Data availability statement

The LC-MS/MS proteomics data have been deposited to the ProteomeXchange Consortium via the PRIDE partner repository with the dataset identifier PXD083069. The atomic models and corresponding cryo-EM maps were deposited in the Protein Data Bank and Electron Microscopy Data Bank under accession codes PDB 35UN and EMD-77193 for AL-252L liver, PDB 13CN and EMD-76964 for AL-150L heart, PDB 13FY and EMD-77052 for AL-150L spleen, and PDB 35US and EMD-77197 for AL-150L kidney. MD data and analyses will be available via a Zenodo archive.

## Acknowledgements

We are grateful to Dr. Esther Bullitt, the Director of the Boston University cryo-EM core facility where the data were collected, and to Dr. Christopher W. Akey for sharing the ATLAS workstation used for local cryo-EM data processing and modelling. We thank Dr. Elena Klimtchuk for generating recombinant V_L_ protein used as a control in this work, and Mei Chen and Steven Kolakowski at the Harvard Center for Mass Spectrometry for help with the proteomics analyses. We gratefully acknowledge the patients’ families for providing tissue samples for research.

## Funding

This research was supported by the National Institutes of Health grants GM135158 and S10OD032253, Robert Champion Amyloidosis Research Fund, Wildflower Foundation, and by the Italian Ministry of University and Research grant FIS00001548. G.A.P. and R.B.B. were supported by the Intramural Research Program of the National Institute of Diabetes and Digestive and Kidney Diseases within the National Institutes of Health (NIH). The contributions of the NIH authors are considered works of the United States government. The findings and conclusions presented in this paper are those of the authors and do not necessarily reflect the views of the NIH or the US Department of Health and Human Services.

## Author contributions

Conceptualization: NH, TP, SJ, FL, OG. Investigation: NH, BS, SW, HC, SJ, GAP, SW, HC, FL. Analysis: NH, SJ, GP, RB, FL, TP, OG. Funding acquisition and resources: VS, FL, HC, CWH, RBB, OG. Data visualization: NH, TP, SJ, GAP. Writing and editing: OG, NH, SJ, GAP, TP. Review: CWH, FL, VS, RBB, SJ, OG.

## Competing interests

The authors declare no competing interests.

## Additional information

Supplementary information including Supplemental Methods, Supplemental references, Supplemental Tables 1-7, and Supplemental Figures 1-22, is available.

## METHODS

### Sample collection and clinical characterization of patients AL-252 and AL-150

Bone marrow aspirate, clinical information, laboratory data, and post-mortem tissues were obtained from the sample biorepository and patient database maintained by the Boston University Amyloidosis Center. Supplemental Methods provide details of patients’ evaluation, and Supplemental Table 1 lists detailed clinical, laboratory and histological characteristics of patients. The clinical case report on patient AL-252^32^ and in vitro studies of recombinant AL-150L protein^33,36^ have been reported previously. Details of tissue histological analysis and gene sequencing are provided in Supplemental Methods.

### Amyloid fibril extraction and proteomic analysis

Fibrils were extracted from the frozen autopsied tissues following published protocols^16,22^. Briefly, overnight tissue digestion with *C. histolyticum* collagenase (5 mg/ml in a buffer containing 20 mM Tris, 140 mM NaCl, 2 mM CaCl_2_, pH 8.0) was followed by 15 cycles of homogenization in Tris EDTA buffer (20 mM Tris, 140 mM NaCl, 10 mM EDTA, pH 8.0) to remove soluble proteins, and by repeated homogenization of the remaining pellet in ultrapure water. The fibril-containing supernatants from the water homogenization cycles were retained. To evaluate the protein yield, 5 ml of each fraction were quantified using a Pierce BCA Protein Assay Kit (Thermo Fisher Scientific). The cryo-EM and proteomic analyses were performed using the visibly cloudy supernatant fractions from the water-extraction cycles. Proteomic analysis of fibril extracts using LC-MS/MS is described in Supplemental Methods and the results are shown in Supplemental Tables 2-5.

### Cryo-EM data collection, image processing and helical reconstruction

UltrAuFoil R1.2/1.3 copper 300-mesh grids (Quantifoil) were glow-discharged for 45 s at 15 mA using a PELCO easiGlow system. A 3-µL aliquot of fibril suspension was applied to a grid and immediately blotted for 3 s at a blot force of 3. The grids were plunge-frozen in liquid ethane using a Vitrobot Mark IV maintained at 4 °C and 100% relative humidity.

Cryo-EM data were collected at the Boston University Cryo-EM Core Facility using a Thermo Fisher Scientific Glacios 2 transmission electron microscope operated at 200 kV and equipped with a Falcon 4i direct electron detector. Data were acquired at a nominal magnification of 130,000×, corresponding to a calibrated pixel size of 0.89 Å pixel⁻¹. An energy-filter slit width of 10 eV was used. Each movie was recorded over 4.10 s and fractionated into 35 fractions.

The total electron exposures were 49.02 e⁻ Å⁻² for AL-252L liver, 49.50 e⁻ Å⁻² for AL-150L heart, 49.14 e⁻ Å⁻² for AL-150L spleen, and 48.70 e⁻ Å⁻² for AL-150L kidney datasets. The corresponding dose rates were 9.89, 9.96, 9.91, and 9.84 e⁻ pixel⁻¹ s⁻¹. The nominal defocus range was −0.8 to −1.9 µm for the AL-252L dataset and −0.8 to −2.5 µm for all AL-150L datasets. A total of 9,890, 13,522, 8,994, and 7,176 exposures were collected for AL-252L liver and AL-150L heart, spleen, and kidney, respectively. Data collection parameters are summarized in Supplemental Table 6; the data processing workflow is shown in Supplemental Fig. 7.

All datasets were processed using CryoSPARC v4.5.3. Raw exposures were imported into CryoSPARC Live and subjected to Patch Motion Correction followed by Patch CTF Estimation. Micrographs were assessed based on image appearance, estimated defocus, CTF fit, total motion, and relative ice thickness. Following initial processing and quality filtering, 6,685, 8,823, 7,251, and 5,833 micrographs were retained for processing of the AL-252L liver, AL-150L heart, AL-150L spleen, and AL-150L kidney datasets, respectively.

Amyloid fibrils were identified using Filament Tracer. Filament coordinates were inspected to remove incorrectly traced fibrils, fibril intersections, fibril ends, and contaminating features. Overlapping fibril segments were extracted using an intersegment separation of 45 Å. Segments used in the final processing branches were extracted or re-extracted in 400-pixel boxes. Supplemental Fig. 8 illustrates fraction of micrographs used for image processing and the 2D /3D particle classification.

Particle stacks were cleaned through iterative rounds of 2D classification and class selection. Classes containing non-amyloid contaminants, overlapping fibrils, poorly aligned segments, fibril ends, or poorly defined fibrillar features were excluded. Initial 3D volumes were generated using *ab-initio* reconstruction with multiple requested classes. The best-defined fibrillar volumes were selected for helical refinement in C1 point-group symmetry.

Helical rise and twist values were evaluated based on the overall map resolution, continuity of the fibril backbone, separation of successive cross-β layers, and definition of side chain densities. Additional rounds of segment re-extraction, 2D and 3D classification, and local and global CTF refinement were performed as required for each dataset. Final maps were generated using Helix Refine. Overall resolutions were estimated using the gold-standard Fourier shell correlation (FSC) criterion of 0.143 from independently refined half-maps (Supplemental Fig. 9).

For hepatic AL-252L fibrils, the initial uncleaned segment stack contained 465,812 segments. Iterative 2D classification was used to remove contaminants and poorly aligned segments. Initial fibrillar volumes were generated using ab-initio reconstruction, and the best-defined volume was selected for helical refinement. Closely related combinations of helical rise and twist were evaluated, followed by further segment cleaning and local and global CTF refinement. The final AL-252L liver reconstruction contained 196,850 segments and was calculated in C1 symmetry using a helical rise of 4.80 Å and a twist of −1.00°. The final map reached an overall resolution of 3.44 Å.

For AL-150L fibrils from heart, spleen, and kidney, the datasets were processed using a similar workflow of filament tracing, segment extraction, iterative 2D classification and class selection, ab-initio reconstruction, helical refinement, 3D classification, and local and global CTF refinement (Supplemental Fig. 7). The initial uncleaned segment stacks contained 413,361, 386,656, and 400,131 segments for the heart, spleen, and kidney datasets, respectively (Supplemental Fig. 8). For each dataset, the best-defined fibrillar volume from ab-initio reconstruction was selected for helical refinement, and additional classification and CTF-refinement steps were used to remove heterogeneous or poorly resolved segment populations.

The final reconstruction of cardiac AL-150L amyloid contained 211,260 segments and was calculated in C1 symmetry using a helical rise of 4.76 Å and a twist of −1.90°, reaching an overall resolution of 2.66 Å. The final AL-150L spleen reconstruction contained 225,004 segments and was calculated in C1 symmetry using a helical rise of 4.76 Å and a twist of −1.85°, reaching an overall resolution of 2.79 Å. The final AL-150L kidney reconstruction contained 99,553 segments and was calculated in C1 symmetry using a helical rise of 4.77 Å and a twist of −1.85°, reaching an overall resolution of 3.40 Å.

### Map post-processing, local resolution estimation, model building and refinement, and confidence maps

Final maps were postprocessed using CryoSPARC. Map sharpening was performed using the CryoSPARC Sharpen job, and local-resolution distributions were estimated using the Local Resolution job. Sharpened and unsharpened maps were inspected in UCSF ChimeraX to evaluate backbone continuity, separation of successive cross-β layers, side-chain definition, and peripheral or unassigned densities. Image-processing and reconstruction statistics are summarized in the Supplemental Table 6.

Atomic models were built and manually adjusted using Coot v0.9.8.96. The AL-252L liver and AL-150L heart atomic models were built de novo into their corresponding cryo-EM maps. For each structure, a single molecular layer was initially traced manually in Coot. Sequence registration was guided by the backbone path, the position of the conserved intramolecular disulfide bond, and densities corresponding to bulky aromatic and branched side chains. Regions lacking interpretable density were not included in the atomic models.

Each single-layer model was expanded to five molecular layers in UCSF ChimeraX using the helical rise and twist determined from the cryo-EM reconstruction. Additional layers were positioned above and below the central layer while preserving the experimentally determined helical relationship between adjacent subunits. The resulting five-layer models were used as starting models for real-space refinement.

Because the AL-150L fibrils isolated from heart, spleen, and kidney showed the same overall fibril fold, the de novo AL-150L heart model was used as the starting model for the spleen and kidney structures. The cardiac amyloid model was fitted independently into the cryo-EM maps for the spleen and kidney amyloids and was refined against each map to obtain the final organ-specific structures.

All five-layer models were refined using Real-space refinement in PHENIX v1.21.2. β-sheet secondary-structure restraints, restraints for the intramolecular disulfide bond, standard stereochemical restraints, and non-crystallographic symmetry restraints between equivalent molecular layers were applied during refinement. Model geometry and agreement with the experimental density were improved through iterative cycles of real-space refinement in PHENIX and manual adjustment in Coot.

Final model quality was assessed using the cryo-EM validation tools implemented in PHENIX. Validation included map–model correlation, MolProbity analysis, clashscore, Ramachandran and rotamer statistics, Cβ deviations, and deviations from ideal bond lengths and bond angles. Numerical model-building and validation statistics are provided in Supplemental Table 6. Molecular graphics and structural analyses were prepared using UCSF ChimeraX.

Helical handedness was evaluated by comparing original and hand-flipped cryo-EM maps with corresponding five-layer atomic models. Candidate models were real-space refined against their respective maps using equivalent settings. Highly sharpened maps were inspected for backbone corrugation and carbonyl orientation. For both AL-252L hepatic and AL-150L cardiac amyloids, the left-handed models showed better agreement with the density, higher map–model correlations, and more favourable Ramachandran and rotamer statistics than the right-handed models, supporting a left-handed fibril architecture (Supplemental Figure 10).

The statistical significance of the extra densities was evaluated using false discovery rate (FDR) thresholding. Confidence maps were generated from the unmasked, unsharpened maps of AL-252L hepatic and AL-150L cardiac amyloids using default parameters, without incorporating local resolution or atomic-model information. Two unidentified orphan densities in AL-252L and two extra densities in AL-150L (inclusion in pocket-1 and the external density near Y75) remained encompassed by the confidence maps at thresholds of 0.9999999 for AL-252L and 0.99999999 for AL-150L, corresponding to FDRs of 0.00001% and 0.000001%, respectively. (Supplemental Figure 11). These FDR values exceed the probability of the extra densities to represent noise.

### Computational studies

Analysis of amyloid structural stability using PDBePISA and ATLAS servers, side chain contacts using APD algorithm, frustration indices using the Protein Frustratometer server, and pKa values using PROPKA-3 server was performed as described in Supplemental Methods. Molecular dynamics simulations were performed as described in Supplemental Methods using CHARMM36mGP force field optimized for collagen structure and dynamics^57^ and utilizing the computational resources of the National Institutes of Health HPC Biowulf cluster, https://hpc.nih.gov.

## Notes

**Conflict of Interest Statement**: The authors declare no competing interests.

http://www.ebi.ac.uk/pride

https://www.rcsb.org/

### Competing Interest Statement

The authors have declared no competing interest.

### Summary of Updates

Few typos missed in the original submission have been corrected. This includes Figure 7 legend.

