## Supplemental Methods, Tables, Figures for "Cryo-EM reveals patient- *versus* organ-specific structural diversity and bound ligands in λ6 light chain amyloids"

- **SUPPLEMENTAL METHODS**
- **SUPPLEMENTAL REFERENCES** (in square brackets; main references are in superscripts)
- **SUPPLEMENTAL TABLES 1 – 7**
- **SUPPLEMENTAL FIGURES 1 – 22**

### SUPPLEMENTAL METHODS

#### Clinical characterization of patients AL-252 and AL-150

***Case AL-252***: A 55-year-old female with treatment-naïve AL amyloidosis and a history of subcapsular liver hemorrhage presented with renal, gastrointestinal, hepatic and autonomic nervous system involvements (Supplemental Table 1). Her evaluation demonstrated λ LC restriction, clonal IgAλ on serum and urine immunofixation electrophoreses, and normal free LC κ/λ ratio. The bone marrow and kidney biopsies were positive for amyloid by Congo red staining; the staining intensity was not documented. The immunohistochemistry of renal tissue showed positive staining with antibody to λ LC. The patient passed away of spontaneous liver rupture following treatment with high-dose melphalan and autologous stem cell transplantation (HDM/SCT) complicated by septic shock. Post-mortem evaluation demonstrated parenchymal amyloid deposition in multiple organs including the liver, and the lack of vascular amyloid deposits. The autopsy report noted very weak Congo red staining in all tissues studied by light macroscopy in polarized view.

***Case AL-150***: A 56-year-old male with treatment-naïve AL amyloidosis presented with soft tissue involvement as demonstrated by macroglossia, submandibular glands enlargement, periorbital ecchymoses, skin petechiae and nodules, and nail dystrophy (Supplemental Table 1). His evaluation showed λ LC restriction, clonal λ on serum and urine immunofixation electrophoreses, and abnormal free LC κ/λ ratio. The fat pad biopsy was positive for amyloid by Congo red staining; the intensity of staining was not documented. The amyloid fibril typing by either immunoelectron microscopy or mass spectrometry was not available at the time of patient's evaluation. He was treated with HDM/SCT followed by revlimid with no treatment response. On the fifth year of follow-up, he developed cardiac and equivocal renal involvements and passed away of heart failure the following year at age 63. Post-mortem evaluation demonstrated parenchymal and vascular amyloid deposition in multiple organs.

The unfixed post-mortem AL-252 and AL-150 tissues were frozen at -80 °C and used for cryo-EM and proteomic analyses as described below. Amyloid fibrils were extracted from the post-mortem liver (AL-252L), heart, spleen and kidney (AL-150L) tissues. Formalin-fixed paraffin-embedded tissue blocks were used for histological studies.

#### Histological and immunoelectron microscopy analyses of tissues

Post-mortem formalin-fixed, paraffin-embedded hepatic tissue from case AL-252 and cardiac, splenic, and renal tissues from case AL-150 were evaluated for amyloid deposition by Congo red staining imaged by light microscopy under both bright-field and polarized light. Amyloid typing of the AL-252 hepatic tissue was performed by immunogold electron microscopy as previously described [1]. Briefly, primary antibodies were polyclonal rabbit anti-human antibodies against λ and κ LCs (Agilent Technologies #A019102-2 and #A019302-2, respectively). Secondary goat anti-rabbit IgG antibodies conjugated to 15-nm gold particles (Ted Pella #15727) were used for immunogold labeling. Electron micrographs were acquired using a JEOL 1011 transmission electron microscope. Amyloid typing was not performed on tissues from case AL-150.

### Gene sequencing

The *IGL* genes were cloned and sequenced from unselected bone marrow plasma cells of patients AL-252 and AL-150 as previously described[22]. Once the *IGLV* gene has been identified, nucleotide sequence errors introduced by FR1 primers were corrected by additional PCR amplification with 5′ primer for the *IGLV6-57* leader region and a 3′ universal *IGLC* primer. The LC sequences, AL-252L and AL-150L, were deposited in GenBank (accession numbers MH996890 and EF589390, respectively).

### 2D-PAGE and Western blot analyses of amyloid fibril extracts

Amyloid fibrils were extracted using a water extraction procedure described in Methods. 2D SDS-PAGE of four tissue fibril extracts was performed under reducing conditions as previously described[16,22]. Briefly, proteins from the cloudy water fraction (200 µg) solubilized in the isoelectric focusing buffer (IEF) were first separated on 11 cm strips with immobilized non-linear pH 3–10 gradient (Bio-Rad) using a Bio-Rad Protean IEF cell, and then on 8–16% polyacrylamide gradient gels (Criterion TGX gels, Bio-Rad). Gels were stained with GelCode Blue Stain Reagent (Thermo Fisher Scientific) and imaged using an ImageQuant LAS 4000 (GE). Spots of interest were excised, and the proteins were identified by LC-MS/MS as described below. For Western blotting, proteins were transferred overnight onto a PVDF membrane (Millipore) using Criterion Blotter (Bio-Rad) and probed with a polyclonal rabbit anti-human λ LCs antibody (Fortis Life Sciences) at 0.125 µg/ml concentration.

### Proteomic analysis of fibril extracts by LC-MS/MS

For proteomic analysis of the extracted fibrils from selected water fractions, 25 µg of proteins were subjected to in-solution digestion with MS-grade trypsin (Thermo Fisher Scientific; enzyme:protein ratio 1:20) overnight at 37 °C. For excised protein spots from the 2D SDS-PAGE, in-gel digestion was performed as described[16,22]. Samples were cleaned up with Thermo SPE C18 tips and the LC-MS/MS analysis was conducted using Q Exactive HF-X High Resolution Orbitrap (Thermo Fisher Scientific) coupled with Ultimate 3000 nanoLC (Thermo Fisher Scientific) at Harvard Center for Mass Spectrometry. Peptides were trapped on a trapping cartridge (300 µm x 5 mm PepMap™ Neo C18 Trap Cartridge, Thermo Fisher Scientific) prior to separation on an analytical column (µPAC, C18 pillar surface, 50 cm bed, Thermo Fisher Scientific). Mobile phase system consisted of water with 1% formic acid (A) and acetonitrile with 1% formic acid (B), using a gradient elution for the total of 110 min (1-32% B over 80 min followed by 32-45% B over 20 min), followed by washout with up to 95% B at a flow rate of 300 nl/min. The mass spectrometer operated in positive mode and data dependent acquisition for all analyses. A full scan ranging from 350 to 1400 m/z was performed with a mass resolution of $12\times10^4$. The top three most intensive precursor ions from each scan were used for MS2 fragmentation with normalized collision energy of 30 at a mass resolution of $3.0x10^4$. Data were processed using the Proteome Discoverer software (Thermo Fisher Scientific) version 3.1, using a human protein database downloaded in November 2024 from UNIPROT and augmented with the AL-252L and AL-150L sequences. The following criteria were used for identification: cleavage enzyme trypsin (semi-specific); cysteine carbamidomethylation (as a static modification in AL-150L samples from spleen and kidney); deamidation of asparagine and glutamine; methionine oxidation; and conversion of N-terminal glutamine into pyroglutamate (as dynamic modifications). Intensity-based label-free calculation of protein abundance in each sample was performed using the precursor ions quantifier node in Proteome Discoverer software.

### Analysis of amyloid packing difference

Pairwise differences in the side chain packing for the nine known λ6-LC amyloid structures (listed in Table 1) were calculated using the amyloid packing differences (APD) software[40] available at https://github.com/3dem/APD. Four structures were determined in the current study (PDB ID: 13CN, EMD-76964; PDB ID: 13FY, EMD-77052; PDB ID: 35US, EMD-77197; PDB ID: 35UN, EMD-77193); one structure was solved in our prior work (PDB ID: 9OKA, EMD-70557); and four structures were retrieved from the Protein Data Bank (PDB ID: 6HUD, EMD-0274; PDB ID: 8CPE, EMD-16780; PDB ID: 9ELS, EMD-48160; PDB ID: 9LJC, EMD-63142). Residue numbering was consistent in all structures.

For each structure, script helix.py was used to extend the protofilament along the helical axis using the twist, rise, box size, and pixel size reported in the corresponding EMDB entry, to generate a nine-layer model. Script contacts.py identified residue-residue contacts, defined as non-hydrogen side chain atoms or $C_\alpha$ atoms for glycine within 6.5 Å distance. Script compare.py was used to perform 36 pairwise comparisons and calculate the APD as $(N_{different} + N_{extra}) / (N_{common} + N_{extra}) \times 100\%$. Here $N_{common}$ and $N_{extra}$ reflect residues ordered in both (common) or only one (extra) of the compared structures, and $N_{different}$ reflects residues with unique contacts. All reported values are XY-APD, which considers common contacts regardless of the layer offset in the two structures along the fibril z-axis. Pairwise XY-APD values were compiled into a symmetric distance matrix and visualized as a heatmap using GraphPad Prism, version 11.0.2 (Supplemental Figure 12A). Bead diagrams showing side-chain contacts comparing structure pairs (Supplemental Figure 12B) are generated using the script compare.py.

**Calculating pKa values, structural frustration indices and energy maps for amyloid core residues**

Fibril core structures of hepatic AL-252L and cardiac AL-150L amyloids (PDB ID: 35UN and 13CN), each containing five molecular layers, were used for the following analyses. First, $pK_a$ values of the ionizable residues in (E84, D85, E86, D88) and near (R24, K82) the acidic-rich segment E84-D88 were predicted using PROPKA3 algorithm[50] version 3.5.1 using default parameters, pH 7.0. The results are listed in Supplemental Table 7.

Second, polarity (Fig. 4C, F) and energy maps (Supplemental Fig. 13) for the fibril cores were generated using Amyloid Illustrator[38] version 3, available at https://zenodo.org/records/15218932 using default parameters, pH7.

Third, structural frustration was evaluated using the Frustratometer 2 server[48], available at http://frustratometer.qb.fcen.uba.ar/, which identifies highly frustrated (dynamic regions with conflicting interactions) and minimally frustrated (stable core) regions in a protein structure. The default Electrostatics_k value of 4.15 was used with the sequence separation of 3 residues. Frustration indices for residues from the central molecule in the 5-layer stack were plotted as a function of residue number (Supplemental Figure 14).

**Evaluating Congo red binding using absorption spectroscopy**

To evaluate dye binding to tissue-extracted amyloids, the Congo red stock solution (2 mg/ml, 2.87 mM) was prepared by dissolving 2 g of the dye in 800 ml of 100% ethanol and adding 30 g NaCl dissolved in 100 ml water, for a final volume of 1 liter. A working stock (100 µM) was prepared by diluting 34.8 µl of Congo red stock into 965.2 µl of assay buffer (10 mM sodium phosphate pH 7.4, 150 mM NaCl) on the day of the experiment. Assays were performed in 96-well flat-bottom polystyrene plates (Greiner, Cat. No. 655101) using a final volume of 300 µL per well. Tissue-extracted amyloid samples were added to the assay buffer to a final protein concentration of 0.05 mg/ml. Congo red working stock (30 µl) was added last to a final concentration of 10 µM. Protein alone in buffer, Congo red alone in buffer, and buffer-only wells served as controls. All samples were incubated for 30 min at room temperature before recording spectra. Recombinant $V_L$ protein, which was generated and purified as described[36] provided negative control. Pre-formed amyloid fibrils of human synthetic $A\beta_{1\text{-}42}$ peptide (StressMarq Biosciences, Cat. No. SPR-487C, Lot No. XA888343) provided positive control. Controls were used at a final protein concentration of 0.05 mg/ml with 10 µM Congo red.

Absorption spectra were recorded using a Tecan Infinite M1000 Pro plate reader controlled by i-control software (version 1.11.1.0), scanning from 400 to 800 nm in 2 nm steps with 25 flashes per read. Data were processed using GraphPad Prism (version 11.0.2). For background corrections, spectra of buffer-alone were subtracted from the buffer+Congo red spectra; spectra of protein alone were subtracted from the protein+Congo red spectra; and the spectrum of amyloid fibril was subtracted from the fibril+Congo red spectrum. Next, the wavelength $\lambda_{max}$ of maximal absorbance was determined for each sample. A red shift in $\lambda_{max}$ relative to Congo red in buffer indicates dye binding to amyloid[52]. All experiments were performed in biological and technical triplicates. Supplemental Fig. 15 shows the results.

**Molecular dynamics (MD) simulations of AL-150L amyloid in complex with ColVI triple helix**

***Model building of amyloid surface and ColVI triple helices for MD simulations.*** To model interactions of the ColVI triple helix with amyloid, we used the coordinates of cardiac AL-150L amyloid (PDB ID: 13CN) to create a small model of amyloid surface centered at Y75. The model contained 42 layers (200 Å long) of segment S64-E84 packed according to the fibril geometry, with a -1.90° twist and a 4.79 Å rise per layer. The N- and C-termini of this segment were not blocked. The Y75 ladder oriented along the amyloid fibril z-axis defined a principal coordinate along which initial configurations of the triple-helical fragments could be placed. Triple-helical fragments containing up to 42 residues per chain (up to 120 Å long) were modeled in four alternative initial configurations relative to the Y75 ladder (face-up, face-down, and rotated 180˚ around z-axis in both up and down configurations, Supplemental Fig. 16B). The *z*-coordinate defines axial sliding of the triple helix along the Y75 ladder, the *y*-coordinate defines the relative lateral position of the triple helix along the amyloid surface, and the *x*-coordinate defines the distance of the triple helix from the amyloid surface. To retain the amyloid structure during simulations, the positions of N, $C_\alpha$ and C atoms of the amyloid backbone were restrained with strong, 500 kJ/mol/nm$^2$ harmonic positional restraints. Additionally, for side chains G65, I67, S69, N72, A74, L76, I78, L81, and T83, which point into the fibril core, the non-hydrogen atomic positions were restrained with 100 kJ/mol/nm$^2$ harmonic positional restraints.

The triple-helical domains of human ColVI were divided into eight fragments; each fragment was no more than 42 residues long and excluded residues violating the XYG-repeat in triplets. Amino acid sequences of these eight fragments are listed below; residues excluded from the canonical sequence due to XYG-motif violations are crossed through.

**α1**

1: GARGPPGLRGDPGFEGERGKPGLPGEKGEAGDPGRPGDLGPVG
2: GYQGMKGEKGSRGEKGSRGPKGYKGEKGKRGIDGVDGVKGEMG
3: GYPGLPGCKGSPGFDGIQGPPGPKGDPGAFGLKGEKGEPGADG ~~EAG~~
4: GRPGSSGPSGDEGQPGEPGPPGEKGEAGDEGNPGPDGAPGERG
5: GGPGERGPRGTPGTRGPRGDPGEAGPQGDQGREGPVGVPGDPG
6: GEAGPIGPKGYRGDEGPPGSEGARGAPGPAGPPGDPGLMGERG ~~EDGPAGNGTEG~~
7: GFPGFPGYPGNRGAPGINGTKGYPGLKGDEGEAGDPG ~~DDNNDIA~~
8: GPRGVKGAKGYRGPEGPQGPPGHQGPPGPDG

**α2**

1: GIPGPSGPKGYRGQKGAKGNMGEPGEPGQKGRQGDPGIEGPIG
2: GFPGPKGVPGFKGEKGEFGADGRKGAPGLAGKNGTDGQKGKLG
3: GRIGPPGCKGDPGNRGPDGYPGEAGSPGERGDQGGKGDPGRPG ~~RRG~~
4: GPPGEIGAKGSKGYQGNSGAPGSPGVKGAKGGPGPRGPKGEPG
5: GRRGDPGTKGSPGSDGPKGEKGDPGPEGPRGLAGEVGNKGAKG
6: GDRGLPGPRGPQGALGEPGKQGSRGDPGDAGPRGDSGQPGPKG ~~DPGRPGFSYPG~~
7: GPRGAPGEKGEPGPRGPEGGRGDFGLKGEPGRKGEKG ~~EPADPG~~
8: GPPGEPGPRGPRGVPGPEGEPGPPGDPGLTG

**α3**

1: GCSGQRGDRGPIGSIGPKGIPGEDGYRGYPGDEGGPGERGPPG
2: GVNGTQGFQGCPGQRGVKGSRGFPGEKGEVGEIGLDGLDGEDG
3: GDKGLPGSSGEKGNPGRRGDKGPRGEKGERGDVGIRGDPGNPG ~~QDSQ~~
4: GERGPKGETGDLGPMGVPGRDGVPGGPGETGKNGGFGRRGPPG
5: GAKGNKGGPGQPGFEGEQGTRGAQGPAGPAGPPGLIGEQGISG

6: GPRGSGGAAGAPGERGRTGPLGRKGEPGEPGPKGGIGNRGPRG ~~ETGDDGRDGVG~~
7: GSEGRRGKKGERGFPGYPGPKGNPGEPGLNGTTGPKG ~~IRGRRG~~
8: GNSGPPGIVGQKGDPGYPGPAGPKGNRGDSG

All prolines in the Y-position were modeled as hydroxyproline (Hyp; O). Based on the cryo-EM structures of ColVI heterotrimers [2], the strands were constructed with α1, α2, and α3 in the leading, middle, and trailing positions in each ColVI fragment. All fragments were constructed according to the 10/3 triple helix model [3]. Remaining atom positions were constructed using IC tables in the CHARMM molecular dynamics simulation package [4].

First, each ColVI fragment was aligned with the center of the Y75 ladder in the *z*- and *y*-directions; the triple helix axis was placed 2 nm away from the Y75 hydroxy oxygen in the *x*-direction. Next, each fragment was placed in four alternative initial orientations, to present two opposite sides and two different orientations of the triple helix to amyloid (Supplemental Fig. 16). From each of these four initial orientations, a 1 μs MD simulation was performed. In total, 32 simulations (8 ColVI fragments, each in 4 orientations) in the presence of the amyloid surface were performed. Additionally, 1 μs trajectories of each fragment were simulated in the absence of amyloid (8 in total).

To prevent ColVI fragments from forming aberrant interactions with the backside or edges of the amyloid model, a 2 nm-wide flat-bottom harmonic positional restraint of 1000 kJ/mol/nm$^2$ was applied to all $C_\alpha$ atoms of ColVI; this enables a fragment to unbind from amyloid surface or slide along it within 4 nm distance in the *x*-, *y*- and *z*-dimension. To prevent ColVI fragments from fraying at the N- and C-termini, the distances between the three N- and C-terminal Gly with the canonical hydrogen bonding carbonyl oxygen of the neighboring X-position amino acid were restrained to a 0.19 nm pair distance with harmonic distance restraints of 100 kJ/mol/nm.

***Molecular dynamics simulations*** The GROMACS 2026.2 simulation package was used to simulate all systems using fully GPU-resident mode on NVIDIA RTX Ada 4500 and 5000 and V100x GPUs [5]. The CHARMM36mGP force field, a reparameterization of CHARMM CMAP terms involving Gly, Pro, and Hyp to best reproduce collagen-mimetic peptide structure and dynamics was used to perform all simulations[57]. The CHARMM TIP3P water model was used [6]. Following the addition of neutralizing quantities of NaCl, 150 mM NaCl were added to all systems to approximate physiological conditions. All simulations were performed at 37° C.

The GROMACS leap-frog integrator was used with a time step of 4 fs, hydrogen mass repartitioning was applied with a factor of 3 [7], and all bonds involving hydrogen were constrained using LINCS [8]. Bonds in TIP3P water were constrained using SETTLE [9]. All systems were initially minimized using steepest descent and then equilibrated in the NPT ensemble using weak coupling thermo- and barostats [10], with $\tau$ = 1 and $\tau$ = 5 ps at 37° C and at 1 bar pressure coupling with $4.5 \times 10^{-5}$ bar$^{-1}$ compressibility every 10 steps, for 2 ns. Production simulations were performed in the NVT ensemble using the velocity rescaling thermostat with a $\tau$ = 1 ps [11]. Protein and solvent atoms were coupled to two separate thermostats with the same parameters. Non-bonded interactions were computed using a 1.2 nm cutoff with a force switching function applied from 1.0 nm. Electrostatics were computed using a 1.2 nm cutoff with Particle mesh Ewald [12]. The results are presented in Fig. 7 and Supplemental Figs. 17-21.

***ColVI–amyloid contact analysis*** To evaluate the propensity of ColVI residues to interact with amyloid surface, we counted contacts between all residues of all simulated ColVI fragments (indexed by *j*) with the closest amyloid residues (indexed by *i*) using a ≤3.5-Å distance cutoff between *any* pair of non-hydrogen atoms in each pair of residues (*a*,*b*), in frames sampled beyond 100 ns, $c_{i,j}$. ColVI contacts were summed based on amino acid identity, ($\alpha$), $c_{i,\alpha}$, and the observed contact likelihood for residue type $\alpha$, $p_{i,\alpha}$ was calculated by normalizing by the sum of $c_{i,\alpha}$. To evaluate the residue-specific amyloid interaction propensity normalized based on the amino acid frequency, the ColVI amino-acid frequency ($f_\alpha$)-weighted contact likelihood, $p^{f\alpha}_{i,\alpha}$, was computed.

$$c_{i,j} = \frac{1}{T - 100ns} \sum_{t>100\ \mathrm{ns}}^{T} H\left(3.5\ \text{Å} - \min_{a\in i, b\in j}\left(|\{\vec{r}_a(t)\} - \{\vec{r}_b(t)\}|\right)\right) f_\alpha = \frac{1}{N\mathrm{res}_{\mathrm{ColVI}}} \sum_{j}^{N\mathrm{res}_{\mathrm{ColVI}}} \delta\left(\alpha - \alpha_j\right)$$

$$c_{i,\alpha} = \sum_{j}^{N\mathrm{res}_{\mathrm{ColVI}}} \delta\left(\alpha - \alpha_j\right)\left(c_{i,j}\right) \qquad c_{i,\alpha}^{f_\alpha} = f_\alpha c_{i,\alpha}$$

$$p_{i,\alpha} = c_{i,\alpha} / \sum_{\alpha}^{M} \sum_{i}^{N\mathrm{res}_{\mathrm{AL150L}}} c_{i,\alpha} \qquad p_{i,\alpha}^{f_\alpha} = c_{i,\alpha}^{f_\alpha} / \sum_{\alpha}^{M} \sum_{i}^{N\mathrm{res}_{\mathrm{AL150L}}} c_{i,\alpha}^{f_\alpha}$$

In the above H is the Heaviside function. To evaluate the average time during which each ColVI residue maintains a persistent contact with amyloid, we computed the time during which each residue pair remained in contact with a 5 ns "grace period".

Frequency-weighted ColVI residue-specific contact likelihoods were used to determine an effective relative free energy of interaction, $\Delta E_\alpha$. The free energy for interactions of each amino acid type in ColVI with Y75 is $\Delta E'_{Y75,\alpha}$, and min($\Delta E'_{Y75,\alpha}$) is subtracted from it to define the relative free energy for interactions of amino acid types with Y75, $\Delta E_{Y75,\alpha}$.

$$\Delta E' = -k_{\mathrm{B}} T \ln(p)$$

$$\Delta E = \Delta E - \min(\Delta E')$$

Here, $k_{\mathrm{B}}$ is the Boltzmann constant and $T$ is the system temperature, 310.15 K. The value $\Delta E_{Y75,\alpha}$ was used to determine the orientational preference of the ColVI fragment in respect to amyloid. For this purpose, we used the following 225 residues of the triple helix.

**α1 lead**
RGPPGLRGDPGFEGERGKPGLPGEKGEAGDPGRPGDLGPVGYQGMKGEKGSRGEKGSRGPKGYK
GEKGKRGIDGVDGVKGEMGYPGLPGCKGSPGFDGIQGPPGPKGDPGAFGLKGEKGEPGADGEAGR
PGSSGPSGDEGQPGEPGPPGEKGEAGDEGNPGPDGAPGERGGPGERGPRGTPGTRGPRGDPGEA
GPQGDQGREGPVGVPGDPGEAGPIGPKGYRGDEGPPGSEGARGAPGPAGPPGDPGLMGERGE

**α2 middle**
PGPSGPKGYRGQKGAKGNMGEPGEPGQKGRQGDPGIEGPIGFPGPKGVPGFKGEKGEFGADGRKG
APGLAGKNGTDGQKGKLGRIGPPGCKGDPGNRGPDGYPGEAGSPGERGDQGGKGDPGRPGRRGP
PGEIGAKGSKGYQGNSGAPGSPGVKGAKGGPGPRGPKGEPGRRGDPGTKGSPGSDGPKGEKGDP
GPEGPRGLAGEVGNKGAKGDRGLPGPRGPQGALGEPGKQGSRGDPGDAGPRGDSGQPGPKGD

**α3 trailing** (QDSQ replaced with QDS, treating S as a G substitution)
SGQRGDRGPIGSIGPKGIPGEDGYRGYPGDEGGPGERGPPGVNGTQGFQGCPGQRGVKGSRGFPG
EKGEVGEIGLDGLDGEDGDKGLPGSSGEKGNPGRRGDKGPRGEKGERGDVGIRGDPGNPGQDSER
GPKGETGDLGPMGVPGRDGVPGGPGETGKNGGFGRRGPPGAKGNKGGPGQPGFEGEQGTRGAQ
GPAGPAGPPGLIGEQGISGPRGSGGAAGAPGERGRTGPLGRKGEPGEPGPKGGIGNRGPRGE

For each Y and X residue at each Cγ angular and height position in the 10/3 helix, relative free energy, ΔEY75,α, was plotted to visualize sites of favorable and unfavorable (lower and higher energy) interactions with the Y75 ladder. The value of ΔEY75,α was integrated over positions at each of 20 unique angular positions (Supplemental Fig. S20B) which define "faces" of the triple helix to elucidate whether there is a face of the triple helix which is expected to preferentially align with the Y75 ladder.

***Helical twist and angle calculations for ColVI triple helices*** To compute the helical twist of each triple-helical chain after MD simulations, one ought to dynamically determine a local cylindrical coordinate system centered on each triplet. In this system, the $C_\alpha$ atom of residue *i* will have the angle measured to the $C_\alpha$ of residue *i*+3, similar to that in the HELANAL α-helix analysis program [13, 14]. To determine this coordinate system, a set of atoms is chosen to compute an axis that best fits through all atoms proximal to the residue of interest for each conformation sampled in an MD trajectory. We used 18 residues surrounding a triplet of interest, as they form a relatively flush and uniform cloud of $C_\alpha$ positions in a 10/3 helix if carefully selected as follows. For a Glycine residue *i*, $C_\alpha$ positions of residues *i*-7 to *i*+10 of the leading chain, *i*-8 to *i*+9 of the middle chain, and *i*-9 to *i*+8 of the trailing chain are used to determine the eigenvector that best fits through these atoms, oriented based on the first and last alpha carbon. These atoms are then centered and rotated such that this principal axis lies along the *z*-axis, with the N-terminus below the C-terminus. In this coordinate system, we measure the twist angle for each triplet using only the Gly-position amino acids (in the GXY motif)in each chain. Because of the 18-residue window, we only measure the twist angles for residues 13-31 for 8 triplets in fragments 1-6, 6 triplets in fragment 7, and 4 triplets in fragment 8.

### SUPPLEMENTAL TABLES

| Characteristics | AL-252 | AL-150 | Reference range |
|---|---|---|---|
| ***Demographic features*** | | | |
| Gender | F | M | |
| Age at presentation, years | 55 | 56 | |
| Age at death, years | 55 | 63 | |
| Cause of death | Liver rupture | Heart failure | |
| ***Echocardiographic features*** | | | |
| Interventricular septal thickness (mm) | 12 | 8 | |
| Ejection fraction, % | 70 | 60 | |
| ***Laboratory parameters*** | | | |
| Bone marrow plasma cells, % | 5 | 5 | |
| LC restriction | λ | λ | |
| SIFE | IgAλ | λ | |
| FLC κ | 12.1 | 9.2 | 3.3–19.4 |
| FLC λ | 34.1 | 439 | 5.7–26.3 |
| FLC ratio | 0.36 | 0.02 | 0.26–1.65 |
| UIFE | IgAλ | λ | |
| BNP, pg/ml | 202 | n/a | 0-53.2 |
| Troponin I, ng/ml | n/a | n/a | <0.033 |
| Serum creatinine, mg/dL | 1.2 | 0.9 | 0.7–1.3 |
| eGFR, ml/min/1.73m$^2$ | 51 | 95 | |
| 24-h urine protein, mg | 8955 | 139 | |
| Alkaline phosphatase, U/L | 147 | 77 | 25–100 |
| ***Organ involvement*** | | | |
| Renal | + | - | |
| Cardiac | - | - | |
| Hepatic | + | - | |
| Neurologic | + | - | |
| Soft tissue | - | + | |
| ***Congo red staining*** | | | |
| Fat pad aspirate | - | + | |
| Bone marrow biopsy | + | - | |
| Kidney biopsy | + | n/a | |
| ***Post-mortem Congo red staining*** | | | |
| Interstitial amyloid deposits | | | |
| Heart | - | + | |
| Kidney | + | + | |
| Liver | + | - | |
| Gastrointestinal | - | + | |
| Lung | - | + | |
| Spleen | + | + | |
| Pancreas | n/a | - | |
| Adrenal glands | + | - | |
| Vascular amyloid deposits | none in all organs | in the medium & small vessels throughout the body | |

**Supplemental Table 1**. **Demographic, clinical, laboratory and histological characteristics of cases AL-252 and AL-150.** Patients' characteristics at initial evaluation and *post-mortem* histology results are listed. Abbreviations: BNP, B-type natriuretic peptide; eGFR, estimated glomerular filtration rate; FLC, free light chain; SIFE, serum immunofixation electrophoresis; UIFE, urine immunofixation electrophoresis.

| Rank | Uniprot Accession | Description | Coverage, % | Abundance, Intensity | # PSMs | # Unique Peptides | # a.a. | MW, kDa |
|---|---|---|---|---|---|---|---|---|
| 1 | AL-252L | Amyloidogenic LC sequence AL-252L OS=Homo sapiens | 96 | 7.73E+09 | 2189 | 62 | 216 | 23.1 |
| 2 | P02649 | Apolipoprotein E OS=Homo sapiens OX=9606 GN=APOE PE=1 SV=1 | 86 | 2.96E+09 | 1357 | 107 | 317 | 36.1 |
| 3 | P02743 | Serum amyloid P-component OS=Homo sapiens OX=9606 GN=APCS PE=1 SV=2 | 59 | 1.78E+09 | 741 | 45 | 223 | 25.4 |
| 4 | P01024 | Complement C3 OS=Homo sapiens OX=9606 GN=C3 PE=1 SV=2 | 29 | 8.6E+08 | 546 | 89 | 1663 | 187 |
| 5 | P04004 | Vitronectin OS=Homo sapiens OX=9606 GN=VTN PE=1 SV=1 | 32 | 7.91E+08 | 391 | 36 | 478 | 54.3 |
| 6 | P02748 | Complement component C9 OS=Homo sapiens OX=9606 GN=C9 PE=1 SV=2 | 26 | 7.28E+08 | 219 | 28 | 559 | 63.1 |
| 7 | P69905 | Hemoglobin subunit alpha OS=Homo sapiens OX=9606 GN=HBA1 PE=1 SV=2 | 77 | 4.41E+08 | 186 | 14 | 142 | 15.2 |
| 8 | P68871 | Hemoglobin subunit beta OS=Homo sapiens OX=9606 GN=HBB PE=1 SV=2 | 86 | 3.46E+08 | 277 | 14 | 147 | 16 |
| 9 | P58166 | Inhibin beta E chain OS=Homo sapiens OX=9606 GN=INHBE PE=1 SV=1 | 32 | 2.33E+08 | 214 | 32 | 350 | 38.5 |
| 10 | P12111 | Collagen alpha-3(VI) chain OS=Homo sapiens OX=9606 GN=COL6A3 PE=1 SV=5 | 23 | 1.90E+08 | 157 | 65 | 3177 | 344 |
| 11 | P0C0L4 | Complement C4-A OS=Homo sapiens OX=9606 GN=C4A PE=1 SV=2 | 22 | 1.24E+08 | 90 | 4 | 1744 | 193 |
| 12 | Q7Z4W1 | L-xylulose reductase OS=Homo sapiens OX=9606 GN=DCXR PE=1 SV=2 | 57 | 1.20E+08 | 101 | 21 | 244 | 25.9 |
| 13 | P12109 | Collagen alpha-1(VI) chain OS=Homo sapiens OX=9606 GN=COL6A1 PE=1 SV=3 | 22 | 8.40E+07 | 76 | 25 | 1028 | 109 |
| 14 | P07360 | Complement component C8 gamma chain OS=Homo sapiens OX=9606 GN=C8G PE=1 SV=3 | 45 | 6.29E+07 | 45 | 13 | 202 | 22.3 |
| 15 | P10909 | Clusterin OS=Homo sapiens OX=9606 GN=CLU PE=1 SV=1 | 26 | 6.25E+07 | 56 | 20 | 449 | 52.5 |
| 16 | P00325 | All-trans-retinol dehydrogenase [NAD(+)] ADH1B OS=Homo sapiens OX=9606 GN=ADH1B PE=1 SV=3 | 31 | 6.15E+07 | 66 | 8 | 375 | 39.8 |
| 17 | P11150 | Hepatic triacylglycerol lipase OS=Homo sapiens OX=9606 GN=LIPC PE=1 SV=3 | 31 | 6.09E+07 | 63 | 19 | 499 | 55.9 |
| 18 | P02768 | Albumin OS=Homo sapiens OX=9606 GN=ALB PE=1 SV=2 | 30 | 5.77E+07 | 56 | 20 | 609 | 69.3 |

| | | | | | | | | |
|---|---|---|---|---|---|---|---|---|
| 19 | P31327 | Carbamoyl-phosphate synthase [ammonia], mitochondrial OS=Homo sapiens OX=9606 GN=CPS1 PE=1 SV=2 | 25 | 5.75E+07 | 52 | 38 | 1500 | 165 |
| 20 | P68104 | Elongation factor 1-alpha 1 OS=Homo sapiens OX=9606 GN=EEF1A1 PE=1 SV=1 | 32 | 5.29E+07 | 58 | 18 | 462 | 50.1 |
| 21 | P01031 | Complement C5 OS=Homo sapiens OX=9606 GN=C5 PE=1 SV=4 | 25 | 5.15E+07 | 79 | 36 | 1676 | 188 |
| 22 | P12110 | Collagen alpha-2(VI) chain OS=Homo sapiens OX=9606 GN=COL6A2 PE=1 SV=4 | 20 | 4.95E+07 | 57 | 18 | 1019 | 109 |
| 23 | P02679 | Fibrinogen gamma chain OS=Homo sapiens OX=9606 GN=FGG PE=1 SV=3 | 32 | 4.59E+07 | 50 | 14 | 453 | 51.5 |
| 24 | P07358 | Complement component C8 beta chain OS=Homo sapiens OX=9606 GN=C8B PE=1 SV=4 | 12 | 4.09E+07 | 34 | 8 | 591 | 66.9 |
| 25 | P05164 | Myeloperoxidase OS=Homo sapiens OX=9606 GN=MPO PE=1 SV=1 | 25 | 4.01E+07 | 32 | 13 | 745 | 83.8 |
| 26 | Q13103 | Secreted phosphoprotein 24 OS=Homo sapiens OX=9606 GN=SPP2 PE=1 SV=1 | 30 | 3.88E+07 | 39 | 7 | 211 | 24.3 |
| 27 | P07357 | Complement component C8 alpha chain OS=Homo sapiens OX=9606 GN=C8A PE=1 SV=2 | 7 | 3.38E+07 | 32 | 6 | 584 | 65.1 |
| 28 | P13671 | Complement component C6 OS=Homo sapiens OX=9606 GN=C6 PE=1 SV=3 | 19 | 3.36E+07 | 44 | 16 | 934 | 105 |
| 29 | P10643 | Complement component C7 OS=Homo sapiens OX=9606 GN=C7 PE=1 SV=2 | 21 | 3.05E+07 | 33 | 16 | 843 | 93.5 |
| 30 | Q9BXR6 | Complement factor H-related protein 5 OS=Homo sapiens OX=9606 GN=CFHR5 PE=1 SV=1 | 12 | 3.02E+07 | 15 | 7 | 569 | 64.4 |
| 31 | Q06033 | Inter-alpha-trypsin inhibitor heavy chain H3 OS=Homo sapiens OX=9606 GN=ITIH3 PE=1 SV=2 | 17 | 3.0E+07 | 63 | 22 | 890 | 99.8 |
| 32 | P02766 | Transthyretin OS=Homo sapiens OX=9606 GN=TTR PE=1 SV=1 | 61 | 2.85E+07 | 34 | 13 | 147 | 15.9 |
| 33 | P08603 | Complement factor H OS=Homo sapiens OX=9606 GN=CFH PE=1 SV=4 | 5 | 2.75E+07 | 14 | 6 | 1231 | 139 |
| 34 | P02675 | Fibrinogen beta chain OS=Homo sapiens OX=9606 GN=FGB PE=1 SV=2 | 23 | 2.62E+07 | 29 | 12 | 491 | 55.9 |
| 35 | P17516 | Aldo-keto reductase family 1 member C4 OS=Homo sapiens OX=9606 GN=AKR1C4 PE=1 SV=3 | 33 | 2.51E+07 | 27 | 9 | 323 | 37 |
| 36 | P00367 | Glutamate dehydrogenase 1, mitochondrial OS=Homo | 28 | 2.47E+07 | 28 | 14 | 558 | 61.4 |

| | | | | | | | | |
|---|---|---|---|---|---|---|---|---|
| | | sapiens OX=9606 GN=GLUD1 PE=1 SV=2 | | | | | | |
| 37 | P60709 | Actin, cytoplasmic 1 OS=Homo sapiens OX=9606 GN=ACTB PE=1 SV=1 | 33 | 2.46E+07 | 29 | 16 | 375 | 41.7 |
| 38 | P07437 | Tubulin beta chain OS=Homo sapiens OX=9606 GN=TUBB PE=1 SV=2 | 42 | 2.32E+07 | 33 | 3 | 444 | 49.6 |
| 39 | Q92954 | Proteoglycan 4 OS=Homo sapiens OX=9606 GN=PRG4 PE=1 SV=3 | 7 | 2.31E+07 | 25 | 12 | 1404 | 151 |
| 40 | P06727 | Apolipoprotein A-IV OS=Homo sapiens OX=9606 GN=APOA4 PE=1 SV=4 | 43 | 2.27E+07 | 41 | 23 | 396 | 45.3 |
| 41 | P05783 | Keratin, type I cytoskeletal 18 OS=Homo sapiens OX=9606 GN=KRT18 PE=1 SV=2 | 27 | 2.19E+07 | 23 | 10 | 430 | 48 |
| 42 | P07099 | Epoxide hydrolase 1 OS=Homo sapiens OX=9606 GN=EPHX1 PE=1 SV=1 | 34 | 2.18E+07 | 41 | 13 | 455 | 52.9 |
| 43 | P30041 | Peroxiredoxin-6 OS=Homo sapiens OX=9606 GN=PRDX6 PE=1 SV=3 | 23 | 2.06E+07 | 22 | 7 | 224 | 25 |
| 44 | P02760 | Protein AMBP OS=Homo sapiens OX=9606 GN=AMBP PE=1 SV=1 | 14 | 2.06E+07 | 14 | 4 | 352 | 39 |
| 45 | P04114 | Apolipoprotein B-100 OS=Homo sapiens OX=9606 GN=APOB PE=1 SV=3 | 10 | 2.03E+07 | 49 | 39 | 4563 | 515 |
| 46 | P00966 | Argininosuccinate synthase OS=Homo sapiens OX=9606 GN=ASS1 PE=1 SV=2 | 37 | 2.03E+07 | 34 | 17 | 412 | 46.5 |
| 47 | P18428 | Lipopolysaccharide-binding protein OS=Homo sapiens OX=9606 GN=LBP PE=1 SV=3 | 19 | 1.98E+07 | 17 | 9 | 481 | 53.4 |
| 48 | Q9UKZ9 | Procollagen C-endopeptidase enhancer 2 OS=Homo sapiens OX=9606 GN=PCOLCE2 PE=1 SV=1 | 21 | 1.79E+07 | 22 | 8 | 415 | 45.7 |
| 49 | Q6UXH0 | Angiopoietin-like protein 8 OS=Homo sapiens OX=9606 GN=ANGPTL8 PE=1 SV=1 | 39 | 1.76E+07 | 25 | 6 | 198 | 22.1 |
| 50 | P01009 | Alpha-1-antitrypsin OS=Homo sapiens OX=9606 GN=SERPINA1 PE=1 SV=3 | 26 | 1.68E+07 | 23 | 11 | 418 | 46.7 |

**Supplemental Table 2. Proteins identified in hepatic AL-252 amyloid tissue extracts by LC-MS/MS.** Abbreviations: PSM, peptide-spectrum match; # a.a., number of amino acids in full-length protein; MW, molecular weight. Top 50 master proteins identified with at least two unique peptides are listed. The proteins are ranked according to abundance. Fibril-forming protein AL-252L (green) and collagen type-VI chains $\alpha$1, $\alpha$2 and $\alpha$3 (pink) are highlighted. In total, 304 proteins have been identified. The proteins and their ranking numbers relevant to this study include: Complement component C1q subunit C (#67); ColVI $\alpha$6-chain (#165), ColXVIII $\alpha$1-chain (#209); and cathepsin D (#88).

| Rank | Uniprot Accession | Description | Coverage, [%] | Abundance, Intensity | # PSMs | # Unique Peptides | # a.a. | MW, kDa |
|---|---|---|---|---|---|---|---|---|
| 1 | AL-150L | Amyloidogenic LC sequence AL-150L OS=Homo sapiens | 100 | 5.61E+09 | 2168 | 153 | 216 | 23.2 |
| 2 | P12883 | Myosin-7 OS=Homo sapiens OX=9606 GN=MYH7 PE=1 SV=5 | 85 | 1.78E+09 | 1680 | 205 | 1935 | 223 |
| 3 | P04004 | Vitronectin OS=Homo sapiens OX=9606 GN=VTN PE=1 SV=1 | 55 | 1.46E+09 | 896 | 118 | 478 | 54.3 |
| 4 | P06727 | Apolipoprotein A-IV OS=Homo sapiens OX=9606 GN=APOA4 PE=1 SV=4 | 80 | 8.23E+08 | 531 | 154 | 396 | 45.3 |
| 5 | P68032 | Actin, alpha cardiac muscle 1 OS=Homo sapiens OX=9606 GN=ACTC1 PE=1 SV=1 | 88 | 4.74E+08 | 423 | 4 | 377 | 42 |
| 6 | P68871 | Hemoglobin subunit beta OS=Homo sapiens OX=9606 GN=HBB PE=1 SV=2 | 97 | 4.53E+08 | 385 | 52 | 147 | 16 |
| 7 | P12111 | Collagen alpha-3(VI) chain OS=Homo sapiens OX=9606 GN=COL6A3 PE=1 SV=5 | 43 | 4.09E+08 | 522 | 203 | 3177 | 344 |
| 8 | P02743 | Serum amyloid P-component OS=Homo sapiens OX=9606 GN=APCS PE=1 SV=2 | 65 | 3.41E+08 | 201 | 39 | 223 | 25.4 |
| 9 | P02649 | Apolipoprotein E OS=Homo sapiens OX=9606 GN=APOE PE=1 SV=1 | 84 | 3.39E+08 | 200 | 70 | 317 | 36.1 |
| 10 | P35555 | Fibrillin-1 OS=Homo sapiens OX=9606 GN=FBN1 PE=1 SV=4 | 33 | 2.89E+08 | 400 | 98 | 2871 | 312 |
| 11 | P12109 | Collagen alpha-1(VI) chain OS=Homo sapiens OX=9606 GN=COL6A1 PE=1 SV=3 | 45 | 2.55E+08 | 252 | 98 | 1028 | 109 |
| 12 | P69905 | Hemoglobin subunit alpha OS=Homo sapiens OX=9606 GN=HBA1 PE=1 SV=2 | 99 | 2.25E+08 | 204 | 52 | 142 | 15.2 |
| 13 | P12110 | Collagen alpha-2(VI) chain OS=Homo sapiens OX=9606 GN=COL6A2 PE=1 SV=4 | 44 | 1.81E+08 | 238 | 88 | 1019 | 109 |
| 14 | P35609 | Alpha-actinin-2 OS=Homo sapiens OX=9606 GN=ACTN2 PE=1 SV=1 | 58 | 1.59E+08 | 233 | 67 | 894 | 104 |
| 15 | P02748 | Complement component C9 OS=Homo sapiens OX=9606 GN=C9 PE=1 SV=2 | 26 | 1.54E+08 | 105 | 32 | 559 | 63.1 |
| 16 | P02656 | Apolipoprotein C-III OS=Homo sapiens OX=9606 GN=APOC3 PE=1 SV=1 | 74 | 1.27E+08 | 86 | 23 | 99 | 10.8 |
| 17 | P02647 | Apolipoprotein A-I OS=Homo sapiens OX=9606 GN=APOA1 PE=1 SV=1 | 80 | 1.26E+08 | 163 | 63 | 267 | 30.8 |
| 18 | P02768 | Albumin OS=Homo sapiens OX=9606 GN=ALB PE=1 SV=2 | 76 | 1.05E+08 | 150 | 66 | 609 | 69.3 |
| 19 | Q8WZ42 | Titin OS=Homo sapiens OX=9606 GN=TTN PE=1 SV=4 | 9 | 9.50E+07 | 387 | 275 | 34350 | 3814 |

| | | | | | | | | |
|---|---|---|---|---|---|---|---|---|
| 20 | P09493 | Tropomyosin alpha-1 chain OS=Homo sapiens OX=9606 GN=TPM1 PE=1 SV=2 | 60 | 9.27E+07 | 72 | 13 | 284 | 32.7 |
| 21 | Q05707 | Collagen alpha-1(XIV) chain OS=Homo sapiens OX=9606 GN=COL14A1 PE=1 SV=3 | 17 | 8.46E+07 | 126 | 38 | 1796 | 193 |
| 22 | Q9UHI8 | A disintegrin and metalloproteinase with thrombospondin motifs 1 OS=Homo sapiens OX=9606 GN=ADAMTS1 PE=1 SV=4 | 21 | 6.13E+07 | 82 | 30 | 967 | 105 |
| 23 | P02675 | Fibrinogen beta chain OS=Homo sapiens OX=9606 GN=FGB PE=1 SV=2 | 51 | 5.69E+07 | 71 | 28 | 491 | 55.9 |
| 24 | P17661 | Desmin OS=Homo sapiens OX=9606 GN=DES PE=1 SV=3 | 47 | 5.45E+07 | 84 | 33 | 470 | 53.5 |
| 25 | Q9UKZ9 | Procollagen C-endopeptidase enhancer 2 OS=Homo sapiens OX=9606 GN=PCOLCE2 PE=1 SV=1 | 55 | 5.31E+07 | 68 | 32 | 415 | 45.7 |
| 26 | Q14896 | Myosin-binding protein C, cardiac-type OS=Homo sapiens OX=9606 GN=MYBPC3 PE=1 SV=4 | 49 | 5.26E+07 | 154 | 80 | 1274 | 141 |
| 27 | P08590 | Myosin light chain 3 OS=Homo sapiens OX=9606 GN=MYL3 PE=1 SV=3 | 67 | 4.73E+07 | 38 | 20 | 195 | 21.9 |
| 28 | P25705 | ATP synthase subunit alpha, mitochondrial OS=Homo sapiens OX=9606 GN=ATP5F1A PE=1 SV=1 | 45 | 4.64E+07 | 72 | 43 | 553 | 59.7 |
| 29 | P07093 | Glia-derived nexin OS=Homo sapiens OX=9606 GN=SERPINE2 PE=1 SV=1 | 59 | 4.42E+07 | 89 | 39 | 398 | 44 |
| 30 | P06576 | ATP synthase subunit beta, mitochondrial OS=Homo sapiens OX=9606 GN=ATP5F1B PE=1 SV=3 | 59 | 4.28E+07 | 91 | 45 | 529 | 56.5 |
| 31 | Q16778 | Histone H2B type 2-E OS=Homo sapiens OX=9606 GN=H2BC21 PE=1 SV=3 | 71 | 4.25E+07 | 50 | 3 | 126 | 13.9 |
| 32 | O75339 | Cartilage intermediate layer protein 1 OS=Homo sapiens OX=9606 GN=CILP PE=1 SV=4 | 25 | 4.25E+07 | 64 | 32 | 1184 | 133 |
| 33 | P78539 | Sushi repeat-containing protein SRPX OS=Homo sapiens OX=9606 GN=SRPX PE=1 SV=1 | 45 | 4.05E+07 | 70 | 39 | 464 | 51.5 |
| 34 | P10909 | Clusterin OS=Homo sapiens OX=9606 GN=CLU PE=1 SV=1 | 47 | 3.98E+07 | 64 | 37 | 449 | 52.5 |
| 35 | P21246 | Pleiotrophin OS=Homo sapiens OX=9606 GN=PTN PE=1 SV=1 | 58 | 3.62E+07 | 110 | 22 | 168 | 18.9 |
| 36 | Q8N474 | Secreted frizzled-related protein 1 OS=Homo sapiens OX=9606 GN=SFRP1 PE=1 SV=1 | 42 | 3.36E+07 | 61 | 27 | 314 | 35.4 |

| | | | | | | | | |
|---|---|---|---|---|---|---|---|---|
| 37 | P07585 | Decorin OS=Homo sapiens OX=9606 GN=DCN PE=1 SV=1 | 47 | 3.36E+07 | 67 | 27 | 359 | 39.7 |
| 38 | O15230 | Laminin subunit alpha-5 OS=Homo sapiens OX=9606 GN=LAMA5 PE=1 SV=8 | 6 | 3.29E+07 | 65 | 25 | 3695 | 400 |
| 39 | P02511 | Alpha-crystallin B chain OS=Homo sapiens OX=9606 GN=CRYAB PE=1 SV=2 | 81 | 3.26E+07 | 82 | 37 | 175 | 20.1 |
| 40 | P68371 | Tubulin beta-4B chain OS=Homo sapiens OX=9606 GN=TUBB4B PE=1 SV=1 | 52 | 3.08E+07 | 69 | 4 | 445 | 49.8 |
| 41 | P98160 | Basement membrane-specific heparan sulfate proteoglycan core protein OS=Homo sapiens OX=9606 GN=HSPG2 PE=1 SV=4 | 20 | 3.03E+07 | 117 | 76 | 4391 | 469 |
| 42 | P48735 | Isocitrate dehydrogenase [NADP], mitochondrial OS=Homo sapiens OX=9606 GN=IDH2 PE=1 SV=2 | 36 | 2.88E+07 | 49 | 25 | 452 | 50.9 |
| 43 | P02452 | Collagen alpha-1(I) chain OS=Homo sapiens OX=9606 GN=COL1A1 PE=1 SV=6 | 3 | 2.72E+07 | 17 | 6 | 1464 | 139 |
| 44 | P62805 | Histone H4 OS=Homo sapiens OX=9606 GN=H4C1 PE=1 SV=2 | 60 | 2.65E+07 | 39 | 16 | 103 | 11.4 |
| 45 | P14555 | Phospholipase A2, membrane associated OS=Homo sapiens OX=9606 GN=PLA2G2A PE=1 SV=2 | 54 | 2.41E+07 | 46 | 18 | 144 | 16.1 |
| 46 | P19429 | Troponin I, cardiac muscle OS=Homo sapiens OX=9606 GN=TNNI3 PE=1 SV=3 | 45 | 2.26E+07 | 44 | 25 | 210 | 24 |
| 47 | P13611 | Versican core protein OS=Homo sapiens OX=9606 GN=VCAN PE=1 SV=3 | 4 | 2.08E+07 | 21 | 12 | 3396 | 373 |
| 48 | P04406 | Glyceraldehyde-3-phosphate dehydrogenase OS=Homo sapiens OX=9606 GN=GAPDH PE=1 SV=3 | 65 | 2.03E+07 | 41 | 31 | 335 | 36 |
| 49 | P17540 | Creatine kinase S-type, mitochondrial OS=Homo sapiens OX=9606 GN=CKMT2 PE=1 SV=2 | 64 | 1.84E+07 | 60 | 34 | 419 | 47.5 |
| 50 | P08123 | Collagen alpha-2(I) chain OS=Homo sapiens OX=9606 GN=COL1A2 PE=1 SV=7 | 5 | 1.82E+07 | 28 | 9 | 1366 | 129 |

**Supplemental Table 3. Proteins identified in cardiac AL-150 amyloid tissue extracts by LC-MS/MS.** Top 50 master proteins identified with at least two unique peptides are listed. Abbreviations: PSM, peptide-spectrum match; # a.a., number of amino acids in full-length protein; MW, molecular weight. The proteins are ranked according to abundance. Fibril-forming protein AL-150L (green) and collagen chains $\alpha$1, $\alpha$2 and $\alpha$3 (pink) are highlighted. In total, 476 proteins have been identified. The proteins and their ranking numbers relevant to this study include: Complement component C1q subunits C (#146), B (#197) and A (#455); ColXVIII $\alpha$1-chain (#204), ColXV $\alpha$1-chain (#265), ColXII $\alpha$1-chain (#276), ColIV $\alpha$2-chain (#313), ColV $\alpha$1-chain (#332), ColIII $\alpha$1-chain (#356); and cathepsin D (#155).

| Rank | Uniprot Accession | Description | Coverage, % | Abundance, Intensity | # PSMs | # Unique Peptides | # a.a. | MW, kDa |
|---|---|---|---|---|---|---|---|---|
| 1 | P68871 | Hemoglobin subunit beta OS=Homo sapiens OX=9606 GN=HBB PE=1 SV=2 | 95 | 1.25E+10 | 1900 | 54 | 147 | 16 |
| 2 | AL-150L | Amyloidogenic LC sequence AL-150L OS=Homo sapiens | 97 | 1.04E+10 | 823 | 42 | 216 | 23.2 |
| 3 | P69905 | Hemoglobin subunit alpha OS=Homo sapiens OX=9606 GN=HBA1 PE=1 SV=2 | 85 | 4.16E+09 | 424 | 56 | 142 | 15.2 |
| 4 | P02748 | Complement component C9 OS=Homo sapiens OX=9606 GN=C9 PE=1 SV=2 | 57 | 3.31E+09 | 339 | 66 | 559 | 63.1 |
| 5 | P04004 | Vitronectin OS=Homo sapiens OX=9606 GN=VTN PE=1 SV=1 | 52 | 2.83E+09 | 435 | 71 | 478 | 54.3 |
| 6 | P05164 | Myeloperoxidase OS=Homo sapiens OX=9606 GN=MPO PE=1 SV=1 | 63 | 2.72E+09 | 485 | 83 | 745 | 83.8 |
| 7 | P01024 | Complement C3 OS=Homo sapiens OX=9606 GN=C3 PE=1 SV=2 | 46 | 2.58E+09 | 515 | 116 | 1663 | 187 |
| 8 | P02649 | Apolipoprotein E OS=Homo sapiens OX=9606 GN=APOE PE=1 SV=1 | 81 | 2.47E+09 | 197 | 60 | 317 | 36.1 |
| 9 | P12111 | Collagen alpha-3(VI) chain OS=Homo sapiens OX=9606 GN=COL6A3 PE=1 SV=5 | 39 | 2.45E+09 | 445 | 149 | 3177 | 344 |
| 10 | P02743 | Serum amyloid P-component OS=Homo sapiens OX=9606 GN=APCS PE=1 SV=2 | 68 | 2.25E+09 | 206 | 30 | 223 | 25.4 |
| 11 | P11678 | Eosinophil peroxidase OS=Homo sapiens OX=9606 GN=EPX PE=1 SV=2 | 53 | 1.78E+09 | 305 | 53 | 715 | 81 |
| 12 | P06727 | Apolipoprotein A-IV OS=Homo sapiens OX=9606 GN=APOA4 PE=1 SV=4 | 73 | 1.48E+09 | 183 | 66 | 396 | 45.3 |
| 13 | P12109 | Collagen alpha-1(VI) chain OS=Homo sapiens OX=9606 GN=COL6A1 PE=1 SV=3 | 54 | 1.40E+09 | 344 | 106 | 1028 | 109 |
| 14 | P59665 | Neutrophil defensin 1 OS=Homo sapiens OX=9606 GN=DEFA1 PE=1 SV=1 | 20 | 1.34E+09 | 48 | 5 | 94 | 10.2 |
| 15 | P60709 | Actin, cytoplasmic 1 OS=Homo sapiens OX=9606 GN=ACTB PE=1 SV=1 | 82 | 1.20E+09 | 265 | 35 | 375 | 41.7 |
| 16 | P01721 | Immunoglobulin lambda variable 6-57 OS=Homo sapiens OX=9606 GN=IGLV6-57 PE=1 SV=2 | 46 | 9.06E+08 | 107 | 5 | 117 | 12.6 |
| 17 | P12110 | Collagen alpha-2(VI) chain OS=Homo sapiens OX=9606 GN=COL6A2 PE=1 SV=4 | 38 | 6.37E+08 | 137 | 53 | 1019 | 109 |
| 18 | P01031 | Complement C5 OS=Homo sapiens OX=9606 GN=C5 PE=1 SV=4 | 53 | 6.16E+08 | 207 | 87 | 1676 | 188 |
| 19 | P35579 | Myosin-9 OS=Homo sapiens OX=9606 GN=MYH9 PE=1 SV=4 | 33 | 6.12E+08 | 195 | 50 | 1960 | 226 |
| 20 | P02768 | Albumin OS=Homo sapiens OX=9606 GN=ALB PE=1 SV=2 | 72 | 5.56E+08 | 134 | 51 | 609 | 69.3 |

| | | | | | | | | |
|---|---|---|---|---|---|---|---|---|
| 21 | O60814 | Histone H2B type 1-K OS=Homo sapiens OX=9606 GN=H2BC12 PE=1 SV=3 | 43 | 5.22E+08 | 71 | 2 | 126 | 13.9 |
| 22 | P07437 | Tubulin beta chain OS=Homo sapiens OX=9606 GN=TUBB PE=1 SV=2 | 73 | 4.68E+08 | 214 | 11 | 444 | 49.6 |
| 23 | P68104 | Elongation factor 1-alpha 1 OS=Homo sapiens OX=9606 GN=EEF1A1 PE=1 SV=1 | 51 | 4.28E+08 | 83 | 19 | 462 | 50.1 |
| 24 | P02452 | Collagen alpha-1(I) chain OS=Homo sapiens OX=9606 GN=COL1A1 PE=1 SV=6 | 11 | 4.09E+08 | 50 | 18 | 1464 | 139 |
| 25 | Q6FI13 | Histone H2A type 2-A OS=Homo sapiens OX=9606 GN=H2AC18 PE=1 SV=3 | 60 | 3.68E+08 | 30 | 2 | 130 | 14.1 |
| 26 | P0C0L5 | Complement C4-B OS=Homo sapiens OX=9606 GN=C4B PE=1 SV=2 | 28 | 3.63E+08 | 117 | 5 | 1744 | 193 |
| 27 | P13671 | Complement component C6 OS=Homo sapiens OX=9606 GN=C6 PE=1 SV=3 | 35 | 3.61E+08 | 103 | 35 | 934 | 105 |
| 28 | P07360 | Complement component C8 gamma chain OS=Homo sapiens OX=9606 GN=C8G PE=1 SV=3 | 77 | 3.55E+08 | 62 | 25 | 202 | 22.3 |
| 29 | P02788 | Lactotransferrin OS=Homo sapiens OX=9606 GN=LTF PE=1 SV=6 | 68 | 3.37E+08 | 114 | 45 | 710 | 78.1 |
| 30 | P08123 | Collagen alpha-2(I) chain OS=Homo sapiens OX=9606 GN=COL1A2 PE=1 SV=7 | 9 | 3.02E+08 | 26 | 11 | 1366 | 129 |
| 31 | P10643 | Complement component C7 OS=Homo sapiens OX=9606 GN=C7 PE=1 SV=2 | 51 | 3.02E+08 | 100 | 42 | 843 | 93.5 |
| 32 | P35555 | Fibrillin-1 OS=Homo sapiens OX=9606 GN=FBN1 PE=1 SV=4 | 26 | 2.95E+08 | 105 | 60 | 2871 | 312 |
| 33 | P07358 | Complement component C8 beta chain OS=Homo sapiens OX=9606 GN=C8B PE=1 SV=4 | 34 | 2.62E+08 | 70 | 24 | 591 | 66.9 |
| 34 | P01857 | Immunoglobulin heavy constant gamma 1 OS=Homo sapiens OX=9606 GN=IGHG1 PE=1 SV=2 | 56 | 2.52E+08 | 80 | 6 | 399 | 43.9 |
| 35 | P02679 | Fibrinogen gamma chain OS=Homo sapiens OX=9606 GN=FGG PE=1 SV=3 | 57 | 2.52E+08 | 81 | 29 | 453 | 51.5 |
| 36 | P02647 | Apolipoprotein A-I OS=Homo sapiens OX=9606 GN=APOA1 PE=1 SV=1 | 59 | 2.52E+08 | 66 | 28 | 267 | 30.8 |
| 37 | P02042 | Hemoglobin subunit delta OS=Homo sapiens OX=9606 GN=HBD PE=1 SV=2 | 89 | 2.28E+08 | 867 | 14 | 147 | 16 |
| 38 | P08311 | Cathepsin G OS=Homo sapiens OX=9606 GN=CTSG PE=1 SV=2 | 54 | 2.15E+08 | 57 | 17 | 255 | 28.8 |
| 39 | O14773 | Tripeptidyl-peptidase 1 OS=Homo sapiens OX=9606 GN=TPP1 PE=1 SV=2 | 33 | 1.99E+08 | 59 | 23 | 563 | 61.2 |
| 40 | P62805 | Histone H4 OS=Homo sapiens OX=9606 GN=H4C1 PE=1 SV=2 | 54 | 1.90E+08 | 44 | 12 | 103 | 11.4 |

| | | | | | | | | |
|---|---|---|---|---|---|---|---|---|
| 41 | P10909 | Clusterin OS=Homo sapiens OX=9606 GN=CLU PE=1 SV=1 | 38 | 1.88E+08 | 57 | 23 | 449 | 52.5 |
| 42 | P35625 | Metalloproteinase inhibitor 3 OS=Homo sapiens OX=9606 GN=TIMP3 PE=1 SV=2 | 59 | 1.87E+08 | 57 | 16 | 211 | 24.1 |
| 43 | P07357 | Complement component C8 alpha chain OS=Homo sapiens OX=9606 GN=C8A PE=1 SV=2 | 30 | 1.76E+08 | 63 | 13 | 584 | 65.1 |
| 44 | Q9BXR6 | Complement factor H-related protein 5 OS=Homo sapiens OX=9606 GN=CFHR5 PE=1 SV=1 | 53 | 1.73E+08 | 50 | 28 | 569 | 64.4 |
| 45 | O43927 | C-X-C motif chemokine 13 OS=Homo sapiens OX=9606 GN=CXCL13 PE=1 SV=1 | 50 | 1.70E+08 | 29 | 7 | 109 | 12.7 |
| 46 | P02675 | Fibrinogen beta chain OS=Homo sapiens OX=9606 GN=FGB PE=1 SV=2 | 45 | 1.64E+08 | 47 | 26 | 491 | 55.9 |
| 47 | P21333 | Filamin-A OS=Homo sapiens OX=9606 GN=FLNA PE=1 SV=4 | 23 | 1.60E+08 | 69 | 38 | 2647 | 281 |
| 48 | Q05707 | Collagen alpha-1(XIV) chain OS=Homo sapiens OX=9606 GN=COL14A1 PE=1 SV=3 | 9 | 1.56E+08 | 54 | 20 | 1796 | 193 |
| 49 | P07355 | Annexin A2 OS=Homo sapiens OX=9606 GN=ANXA2 PE=1 SV=2 | 56 | 1.52E+08 | 49 | 22 | 339 | 38.6 |
| 50 | P08758 | Annexin A5 OS=Homo sapiens OX=9606 GN=ANXA5 PE=1 SV=2 | 63 | 1.48E+08 | 41 | 21 | 320 | 35.9 |

**Supplemental Table 4. Proteins identified in splenic AL-150 amyloid tissue extracts by LC-MS/MS.** Top 50 master proteins identified with at least two unique peptides are listed. Abbreviations: PSM, peptide-spectrum match; # a.a., number of amino acids in full-length protein; MW, molecular weight. The proteins are ranked according to abundance. Fibril-forming protein AL-150L (green), collagen chains (pink), and cathepsin (orange) are highlighted. In total, 1091 proteins have been identified. The proteins and their ranking numbers relevant to this study include: Complement component C1q subunits C (#69), B (#130) and A (#299); ColIII $\alpha$1-chain (#83), ColIV $\alpha$1-chain (#212), Col-IV $\alpha$2-chain (#213), ColXXI $\alpha$1-chain (#224), ColXVIII $\alpha$1-chain (#289) and other collagen chains (#305-#878); cathepsins G (#38), D (#97) and B (#642).

| Rank | Uniprot Accession | Description | Coverage, % | Abundance, Intensity | # PSMs | # Unique Peptides | # a.a. | MW, kDa |
|---|---|---|---|---|---|---|---|---|
| 1 | P68871 | Hemoglobin subunit beta OS=Homo sapiens OX=9606 GN=HBB PE=1 SV=2 | 95 | 7.90E+09 | 1376 | 62 | 147 | 16 |
| 2 | P69905 | Hemoglobin subunit alpha OS=Homo sapiens OX=9606 GN=HBA1 PE=1 SV=2 | 85 | 3.40E+09 | 407 | 49 | 142 | 15.2 |
| 3 | P07911 | Uromodulin OS=Homo sapiens OX=9606 GN=UMOD PE=1 SV=1 | 44 | 3.14E+09 | 426 | 44 | 640 | 69.7 |
| 4 | P02452 | Collagen alpha-1(I) chain OS=Homo sapiens OX=9606 GN=COL1A1 PE=1 SV=6 | 15 | 2.30E+09 | 140 | 27 | 1464 | 139 |
| 5 | P08123 | Collagen alpha-2(I) chain OS=Homo sapiens OX=9606 GN=COL1A2 PE=1 SV=7 | 12 | 1.88E+09 | 78 | 23 | 1366 | 129 |
| 6 | P35555 | Fibrillin-1 OS=Homo sapiens OX=9606 GN=FBN1 PE=1 SV=4 | 46 | 1.54E+09 | 327 | 113 | 2871 | 312 |
| 7 | P98164 | Low-density lipoprotein receptor-related protein 2 OS=Homo sapiens OX=9606 GN=LRP2 PE=1 SV=3 | 38 | 1.53E+09 | 529 | 170 | 4655 | 522 |
| 8 | AL-150L | Amyloidogenic LC sequence AL-150L OS=Homo sapiens | 87 | 1.11E+09 | 189 | 13 | 216 | 23.2 |
| 9 | P62805 | Histone H4 OS=Homo sapiens OX=9606 GN=H4C1 PE=1 SV=2 | 58 | 1.03E+09 | 119 | 15 | 103 | 11.4 |
| 10 | P15144 | Aminopeptidase N OS=Homo sapiens OX=9606 GN=ANPEP PE=1 SV=4 | 49 | 9.77E+08 | 230 | 62 | 967 | 110 |
| 11 | P68104 | Elongation factor 1-alpha 1 OS=Homo sapiens OX=9606 GN=EEF1A1 PE=1 SV=1 | 56 | 8.31E+08 | 146 | 23 | 462 | 50.1 |
| 12 | P02768 | Albumin OS=Homo sapiens OX=9606 GN=ALB PE=1 SV=2 | 75 | 8.09E+08 | 161 | 53 | 609 | 69.3 |
| 13 | P02649 | Apolipoprotein E OS=Homo sapiens OX=9606 GN=APOE PE=1 SV=1 | 61 | 7.59E+08 | 82 | 30 | 317 | 36.1 |
| 14 | P63261 | Actin, cytoplasmic 2 OS=Homo sapiens OX=9606 GN=ACTG1 PE=1 SV=1 | 73 | 6.93E+08 | 172 | 24 | 375 | 41.8 |
| 15 | P49411 | Elongation factor Tu, mitochondrial OS=Homo sapiens OX=9606 GN=TUFM PE=1 SV=3 | 70 | 6.87E+08 | 127 | 41 | 455 | 49.8 |
| 16 | P12111 | Collagen alpha-3(VI) chain OS=Homo sapiens OX=9606 GN=COL6A3 PE=1 SV=5 | 30 | 6.64E+08 | 191 | 82 | 3177 | 344 |
| 17 | Q16822 | Phosphoenolpyruvate carboxykinase [GTP], mitochondrial OS=Homo sapiens OX=9606 GN=PCK2 PE=1 SV=4 | 70 | 6.48E+08 | 204 | 50 | 640 | 70.7 |
| 18 | P06727 | Apolipoprotein A-IV OS=Homo sapiens OX=9606 GN=APOA4 PE=1 SV=4 | 66 | 6.05E+08 | 103 | 39 | 396 | 45.3 |

| | | | | | | | | |
|---|---|---|---|---|---|---|---|---|
| 19 | P05141 | ADP/ATP translocase 2 OS=Homo sapiens OX=9606 GN=SLC25A5 PE=1 SV=7 | 74 | 5.75E+08 | 130 | 10 | 298 | 32.8 |
| 20 | P04004 | Vitronectin OS=Homo sapiens OX=9606 GN=VTN PE=1 SV=1 | 40 | 5.68E+08 | 123 | 31 | 478 | 54.3 |
| 21 | P21810 | Biglycan OS=Homo sapiens OX=9606 GN=BGN PE=1 SV=2 | 59 | 4.87E+08 | 147 | 28 | 368 | 41.6 |
| 22 | P48735 | Isocitrate dehydrogenase [NADP], mitochondrial OS=Homo sapiens OX=9606 GN=IDH2 PE=1 SV=2 | 58 | 4.79E+08 | 127 | 38 | 452 | 50.9 |
| 23 | P25705 | ATP synthase subunit alpha, mitochondrial OS=Homo sapiens OX=9606 GN=ATP5F1A PE=1 SV=1 | 52 | 4.13E+08 | 94 | 35 | 553 | 59.7 |
| 24 | P02461 | Collagen alpha-1(III) chain OS=Homo sapiens OX=9606 GN=COL3A1 PE=1 SV=4 | 4 | 4.08E+08 | 28 | 11 | 1466 | 139 |
| 25 | P27144 | Adenylate kinase 4, mitochondrial OS=Homo sapiens OX=9606 GN=AK4 PE=1 SV=1 | 67 | 4.00E+08 | 101 | 32 | 223 | 25.3 |
| 26 | P21796 | Non-selective voltage-gated ion channel VDAC1 OS=Homo sapiens OX=9606 GN=VDAC1 PE=1 SV=2 | 80 | 3.98E+08 | 93 | 18 | 283 | 30.8 |
| 27 | Q02252 | Methylmalonate-semialdehyde/malonate-semialdehyde dehydrogenase [acylating], mitochondrial OS=Homo sapiens OX=9606 GN=ALDH6A1 PE=1 SV=2 | 56 | 3.92E+08 | 96 | 41 | 535 | 57.8 |
| 28 | P27338 | Amine oxidase [flavin-containing] B OS=Homo sapiens OX=9606 GN=MAOB PE=1 SV=3 | 57 | 3.75E+08 | 72 | 22 | 520 | 58.7 |
| 29 | P12109 | Collagen alpha-1(VI) chain OS=Homo sapiens OX=9606 GN=COL6A1 PE=1 SV=3 | 47 | 3.74E+08 | 124 | 48 | 1028 | 109 |
| 30 | P19440 | Glutathione hydrolase 1 proenzyme OS=Homo sapiens OX=9606 GN=GGT1 PE=1 SV=2 | 43 | 3.44E+08 | 70 | 12 | 569 | 61.4 |
| 31 | O60814 | Histone H2B type 1-K OS=Homo sapiens OX=9606 GN=H2BC12 PE=1 SV=3 | 42 | 3.44E+08 | 50 | 2 | 126 | 13.9 |
| 32 | P02458 | Collagen alpha-1(II) chain OS=Homo sapiens OX=9606 GN=COL2A1 PE=1 SV=3 | 4 | 3.34E+08 | 9 | 7 | 1487 | 142 |
| 33 | Q96CM8 | Medium-chain acyl-CoA ligase ACSF2, mitochondrial OS=Homo sapiens OX=9606 GN=ACSF2 PE=1 SV=2 | 46 | 3.32E+08 | 83 | 34 | 615 | 68.1 |
| 34 | Q00610 | Clathrin heavy chain 1 OS=Homo sapiens OX=9606 GN=CLTC PE=1 SV=5 | 50 | 3.24E+08 | 162 | 56 | 1675 | 192 |
| 35 | P06576 | ATP synthase subunit beta, mitochondrial OS=Homo sapiens OX=9606 GN=ATP5F1B PE=1 SV=3 | 71 | 3.14E+08 | 99 | 37 | 529 | 56.5 |

| | | | | | | | | |
|---|---|---|---|---|---|---|---|---|
| 36 | O60494 | Cubilin OS=Homo sapiens OX=9606 GN=CUBN PE=1 SV=5 | 22 | 3.14E+08 | 139 | 64 | 3623 | 399 |
| 37 | P45880 | Voltage-dependent anion-selective channel protein 2 OS=Homo sapiens OX=9606 GN=VDAC2 PE=1 SV=2 | 57 | 3.06E+08 | 41 | 15 | 294 | 31.5 |
| 38 | O60656 | UDP-glucuronosyltransferase 1A9 OS=Homo sapiens OX=9606 GN=UGT1A9 PE=1 SV=1 | 49 | 2.95E+08 | 109 | 22 | 530 | 59.9 |
| 39 | Q7Z4W1 | L-xylulose reductase OS=Homo sapiens OX=9606 GN=DCXR PE=1 SV=2 | 66 | 2.80E+08 | 43 | 18 | 244 | 25.9 |
| 40 | P50440 | Glycine amidinotransferase, mitochondrial OS=Homo sapiens OX=9606 GN=GATM PE=1 SV=1 | 65 | 2.63E+08 | 98 | 41 | 423 | 48.4 |
| 41 | P08473 | Neprilysin OS=Homo sapiens OX=9606 GN=MME PE=1 SV=2 | 56 | 2.58E+08 | 118 | 37 | 750 | 85.5 |
| 42 | P05023 | Sodium/potassium-transporting ATPase subunit alpha-1 OS=Homo sapiens OX=9606 GN=ATP1A1 PE=1 SV=1 | 33 | 2.48E+08 | 58 | 34 | 1023 | 113 |
| 43 | P16444 | Dipeptidase 1 OS=Homo sapiens OX=9606 GN=DPEP1 PE=1 SV=3 | 47 | 2.35E+08 | 65 | 18 | 411 | 45.6 |
| 44 | P02511 | Alpha-crystallin B chain OS=Homo sapiens OX=9606 GN=CRYAB PE=1 SV=2 | 64 | 2.16E+08 | 65 | 14 | 175 | 20.1 |
| 45 | P04406 | Glyceraldehyde-3-phosphate dehydrogenase OS=Homo sapiens OX=9606 GN=GAPDH PE=1 SV=3 | 73 | 2.07E+08 | 74 | 29 | 335 | 36 |
| 46 | P16662 | UDP-glucuronosyltransferase 2B7 OS=Homo sapiens OX=9606 GN=UGT2B7 PE=1 SV=2 | 34 | 2.06E+08 | 46 | 16 | 529 | 60.7 |
| 47 | Q08AH3 | Acyl-coenzyme A synthetase ACSM2A, mitochondrial OS=Homo sapiens OX=9606 GN=ACSM2A PE=1 SV=2 | 58 | 1.99E+08 | 73 | 7 | 577 | 64.2 |
| 48 | P12110 | Collagen alpha-2(VI) chain OS=Homo sapiens OX=9606 GN=COL6A2 PE=1 SV=4 | 28 | 1.95E+08 | 70 | 27 | 1019 | 109 |
| 49 | P35232 | Prohibitin 1 OS=Homo sapiens OX=9606 GN=PHB1 PE=1 SV=1 | 79 | 1.82E+08 | 44 | 24 | 272 | 29.8 |
| 50 | Q99623 | Prohibitin-2 OS=Homo sapiens OX=9606 GN=PHB2 PE=1 SV=2 | 70 | 1.82E+08 | 46 | 19 | 299 | 33.3 |

**Supplemental Table 5. Proteins identified in renal AL-150 amyloid tissue extracts by LC-MS/MS.** Top 50 master proteins identified with at least two unique peptides are listed. Abbreviations: PSM, peptide-spectrum match; # a. a., number of amino acids in full-length protein; MW, molecular weight. The proteins are ranked according to abundance. Fibril-forming protein AL-150L (green) and collagen chains (pink) are highlighted. In total, 1296 proteins have been identified. The proteins and their rankings relevant to this study include: C1q subunits C (#554) and B (#688); ColXXI α1-chain (#64), ColV α1-chain (#128), ColIV α1-chain (#141), ColXVIII α1-chain (#171), ColXIV α1-chain (#192) and other collagen chains (#234 - #1021); and cathepsin D (#224).

| Parameters | Values | | | |
|---|---|---|---|---|
| | AL-252L liver | AL-150L heart | AL-150L spleen | AL-150L kidney |
| | **Data Collection** | | | |
| Magnification (X) | 130,000 | 130,000 | 130,000 | 130,000 |
| Voltage (kV) | 200 | 200 | 200 | 200 |
| Electron exposure ($e^-$/Å$^2$) | 49.02 | 49.50 | 49.14 | 48.70 |
| Dose rate ($e^-$/px/s) | 9.89 | 9.96 | 9.91 | 9.84 |
| Defocus range (µM) | −0.8 to −1.9 | −0.8 to −2.5 | −0.8 to −2.5 | −0.8 to −2.5 |
| Pixel size (Å) | 0.89 | 0.89 | 0.89 | 0.89 |
| Camera | Falcon 4i | Falcon 4i | Falcon 4i | Falcon 4i |
| Energy Filter slit width (eV) | 10 | 10 | 10 | 10 |
| Micrographs (#) | 9,890 | 13,522 | 8,994 | 7,176 |
| | **Data processing** | | | |
| Curated micrographs (#) | 6,685 | 8,823 | 7,251 | 5,833 |
| Box size (px) | 400 | 400 | 400 | 400 |
| Segment separation (Å) | 45 | 45 | 45 | 45 |
| Initial uncleaned segment stack (#) | 465,812 | 413,361 | 386,656 | 400,131 |
| Helical rise (Å) | 4.80 | 4.76 | 4.76 | 4.77 |
| Helical twist (°) | -1.00 | -1.90 | -1.85 | -1.85 |
| Particle duplication symmetry | C1 | C1 | C1 | C1 |
| Final segments (#) | 196,850 | 211,260 | 225,004 | 99,553 |
| Map resolution at 0.143 FSC cutoff (Å) | 3.44 | 2.66 | 2.79 | 3.40 |
| | **Model building and validation** | | | |
| Non-hydrogen atoms (#) | 2665 | 3480 | 3510 | 3440 |
| Subunits | 5 | 5 | 5 | 5 |
| CC mask | 0.65 | 0.88 | 0.81 | 0.68 |
| Bond length RMSD (Å) | 0.002 | 0.002 | 0.002 | 0.002 |
| Bond angles RMSD (°) | 0.530 | 0.436 | 0.469 | 0.468 |
| Clash score | 10.91 | 5.36 | 9.00 | 6.76 |
| Ramachandran Outliers (%) | 0.00 | 0.00 | 0.00 | 0.00 |
| Ramachandran Allowed (%) | 11.76 | 2.30 | 5.68 | 1.16 |
| Ramachandran Favoured (%) | 88.24 | 97.70 | 94.32 | 98.84 |
| Rotamer Outliers (%) | 1.64 | 0.00 | 0.00 | 0.00 |
| Cβ outliers (%) | 0.00 | 0.00 | 0.00 | 0.00 |
| MolProbity score | 2.31 | 1.35 | 1.87 | 1.37 |

**Supplemental Table 6. Cryo-EM data collection, processing, and model validation statistics**. Data collection, image processing, reconstruction, atomic model parameters, and validation statistics for the four structures of amyloid fibrils determined in the current study for two patients, AL-252L and AL-150.

| $V_L$ a.a. | AL-252L | AL-150L | Reference pKa |
|---|---|---|---|
| **R24** | 15.44 | 15.14 | 12.50 |
| **K82** | 10.33 | 10.12 | 10.5 |
| **E84** | 2.42 | 4.59 | 4.50 |
| **D85** | 5.29 | 5.53 | 3.80 |
| **E86** | 6.50 | 5.77 | 4.50 |
| **D88** | 11.51 | 8.68 | 3.80 |

**Supplemental Table 7**. **Predicted pKa values of the ionizable side chains in the acidic-rich segment E84-D88 and the basic side chains of R24 and K82 located nearby**. Continuous amino acid numbering is used. The values are predicted based on the structures of hepatic AL-252L and cardiac AL-150L amyloids using PROPKA-3 server[50], available at https://github.com/jensengroup/propka-3.0 Normal pKa values for free amino acids are provided for reference.

### SUPPLEMENTAL FIGURES

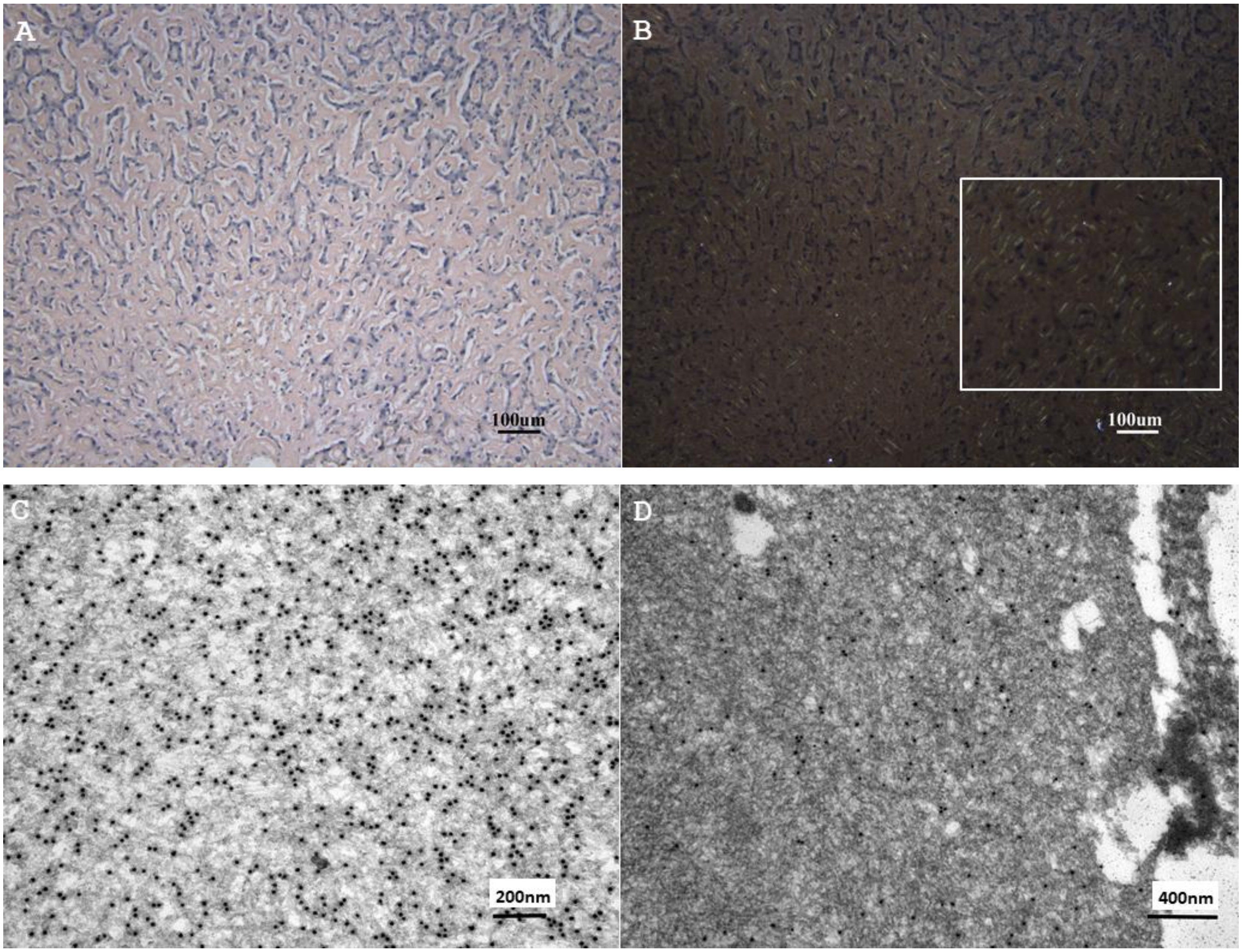


**Supplemental Figure 1**. **Case AL-252 histological analyses of the post-mortem hepatic tissue**.

**A,** Light microscopy images of Congo red-stained tissues indicate amyloid deposits (original magnification ×100). **B,** Areas in panel **A** viewed by polarized microscopy show amyloid deposits with very weak green birefringence (original magnification ×100). Zoomed-in view (boxed) shows area with weak birefringence.

**C, D,** Electron micrographs of post-mortem hepatic tissue show haystack-like deposits of amyloid fibrils in the extracellular space. Immunogold labeling shows numerous electron-dense deposits with antibody against Ig λ-LC (original magnification ×60,000, **C**), and no immunoreactivity with antibody against Ig κ-LC (original magnification ×40,000, **D**).

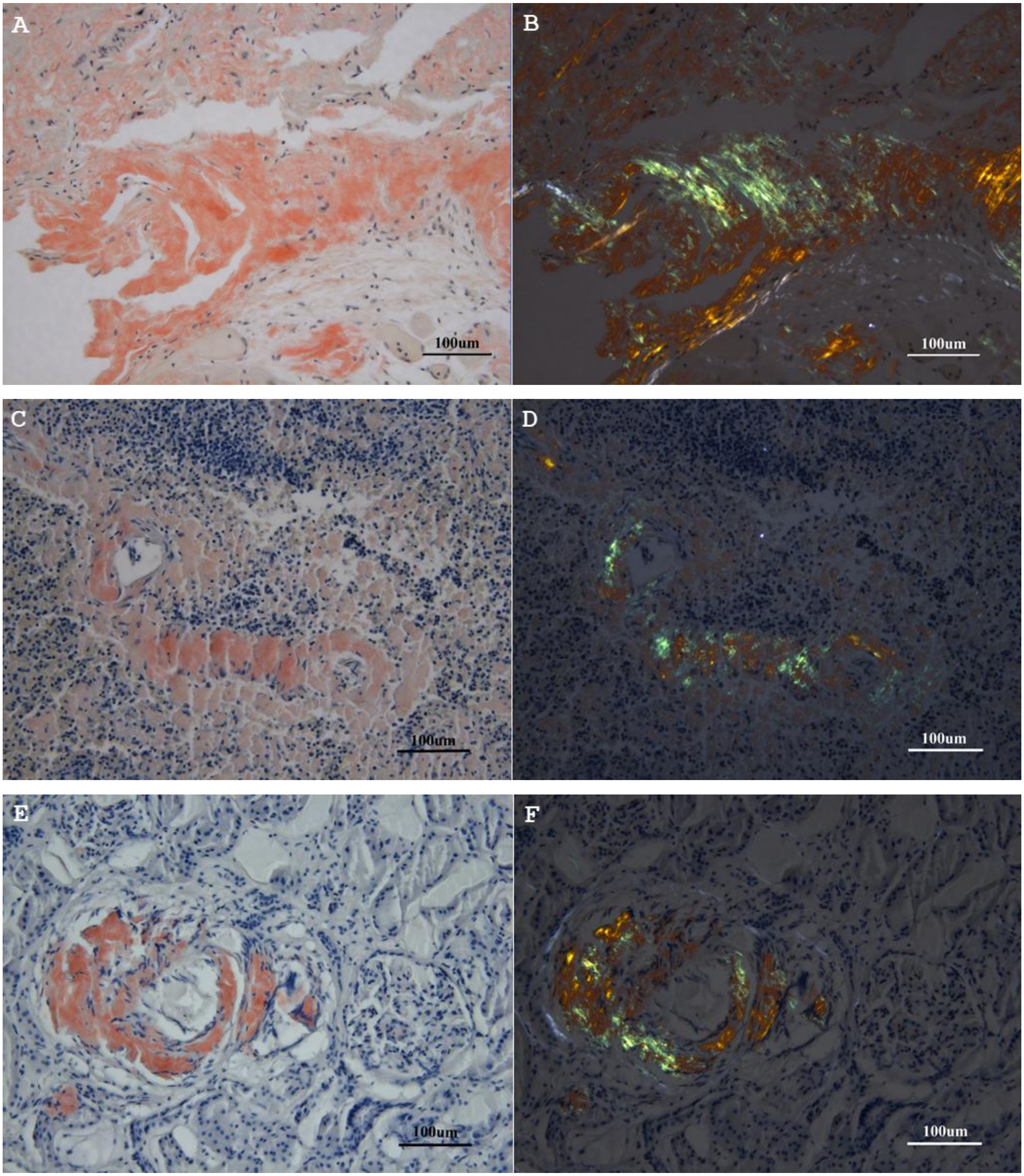


**Supplemental Figure 2**. **Case AL-150 histological analyses of the post-mortem tissues**.

Cardiac (**A, B**), splenic (**C, D**) and renal (**E, F**) tissues are shown. **A, C, E,** Light microscopy images of Congo-red-stained tissues indicate amyloid deposits. **B, D, F,** Same areas viewed by polarized microscopy show amyloid deposits with green birefringence (original magnification ×200 in all panels).

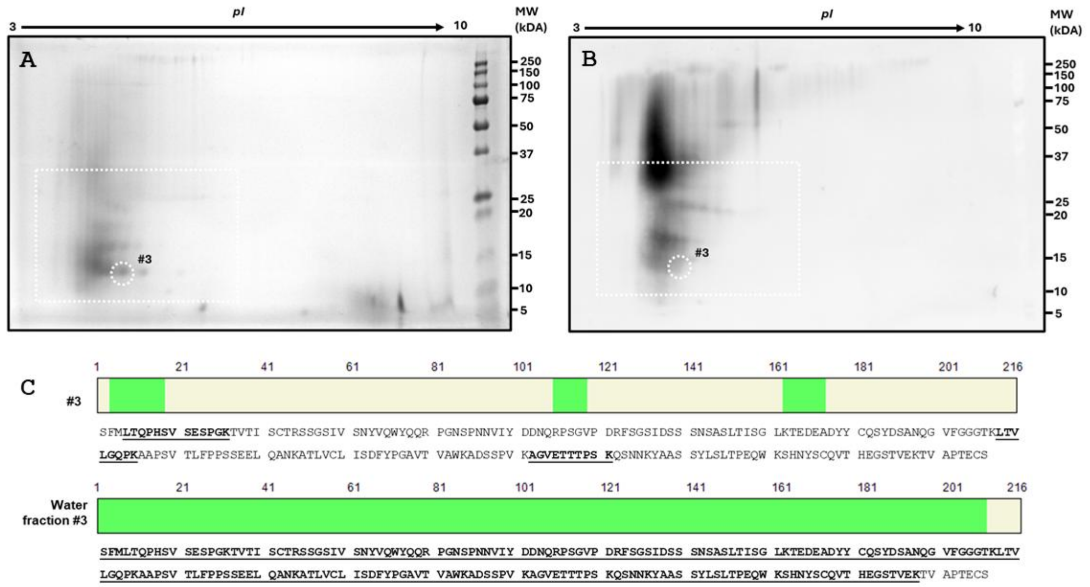


**Supplemental Figure 3. Analysis of AL-252 proteins in fibril extracts from hepatic tissue using 2D SDS-PAGE, 2D Western blotting, and LC-MS/MS**. **A, B**. Coomassie-stained gel (**A**) and the corresponding 2D Western blot (**B**) probed with a primary anti-human λ-LC antibody. A representative spot (circled and marked #3) in the most prominent train of spots was excised and analyzed by LC-MS/MS, along with the entire water fraction. **C**. The position (in green) and sequence (underlined bold) of the peptides identified by LC-MS/MS in spot #3 and in the entire water fraction 3.

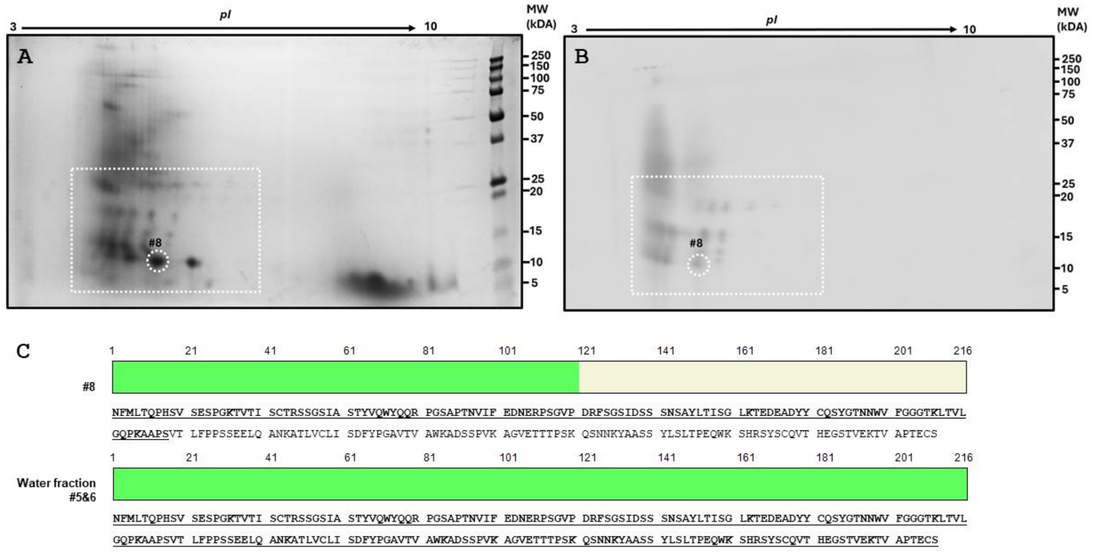


**Supplemental Figure 4**. **Analysis of AL-150 proteins in fibril extracts from cardiac tissue using 2D SDS-PAGE, 2D Western blotting, and LC-MS/MS**. Data are shown for combined water fractions 5 and 6. **A, B**. Coomassie-stained SDS-PAGE (**A**) and the corresponding 2D Western blot (**B**) probed with a primary anti-human λ-LC antibody. The most prominent spot #8 (circled) was excised and analyzed by LC-MS/MS, along with the entire water fractions. **C**. The position (in green) and sequence (bold underlined) of the peptides identified in spot #8 and in the entire water fractions 5 and 6.

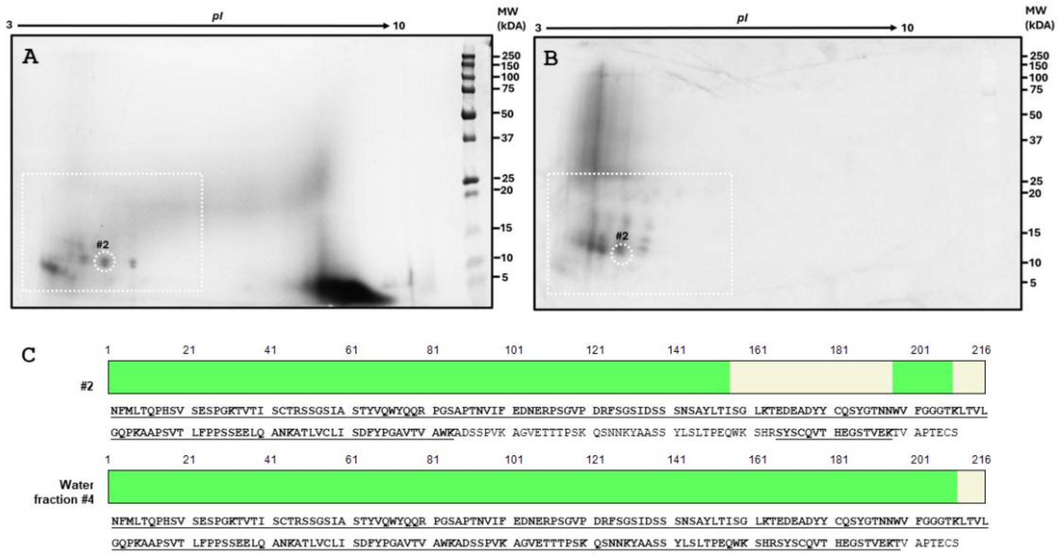


**Supplemental Figure 5**. **Analysis of AL-150 proteins in fibril extracts from splenic tissue using 2D SDS-PAGE, 2D Western blotting, and LC-MS/MS**. **A, B**. Coomassie-stained SDS-PAGE (**A**) and the corresponding 2D Western blot (**B**) probed with a primary anti-human λ-LC antibody. A representative spot (circled and marked #2) in the most prominent train of spots was excised and analyzed by LC-MS/MS, along with the entire water fraction. **C**. The position (in green) and sequence (bold underlined) of the peptides identified in spot #2 and in the entire water fraction.

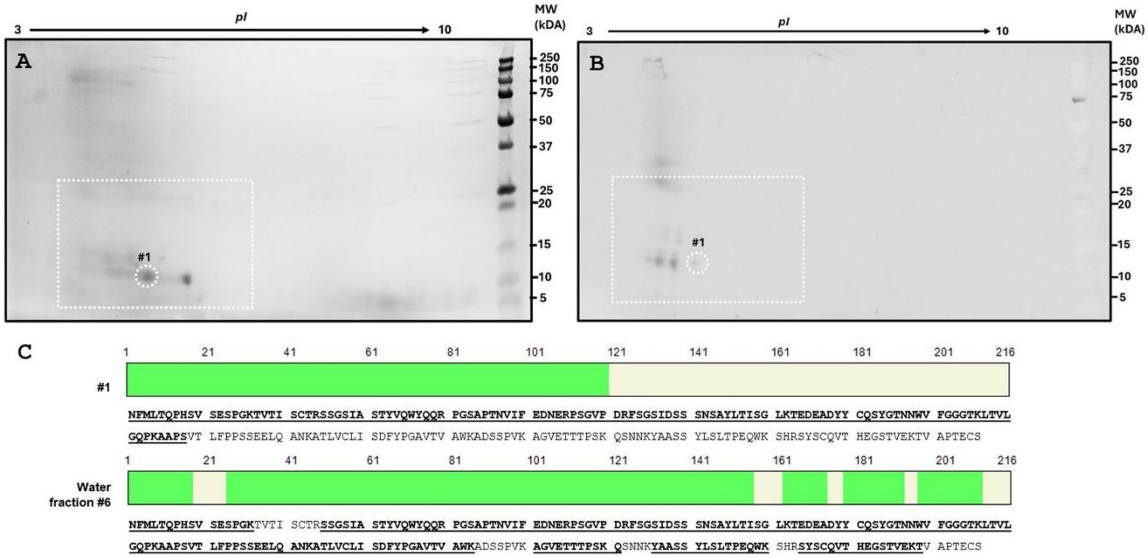


**Supplemental Figure 6**. **Analysis of AL-150 proteins in renal tissue fibril extract using 2D SDS-PAGE, 2D Western blotting, and LC-MS/MS**. **A, B**. Coomassie-stained SDS-PAGE (**A**) and the corresponding 2D Western blot (**B**) probed with a primary anti-human λ-LC antibody. A representative spot (circled and marked #1) in the most prominent train of spots was excised and analyzed by LC-MS/MS, along with the entire water fraction. **C**. The position (in green) and sequence (bold underlined) of the peptides identified in spot #1 and in the entire water fraction.

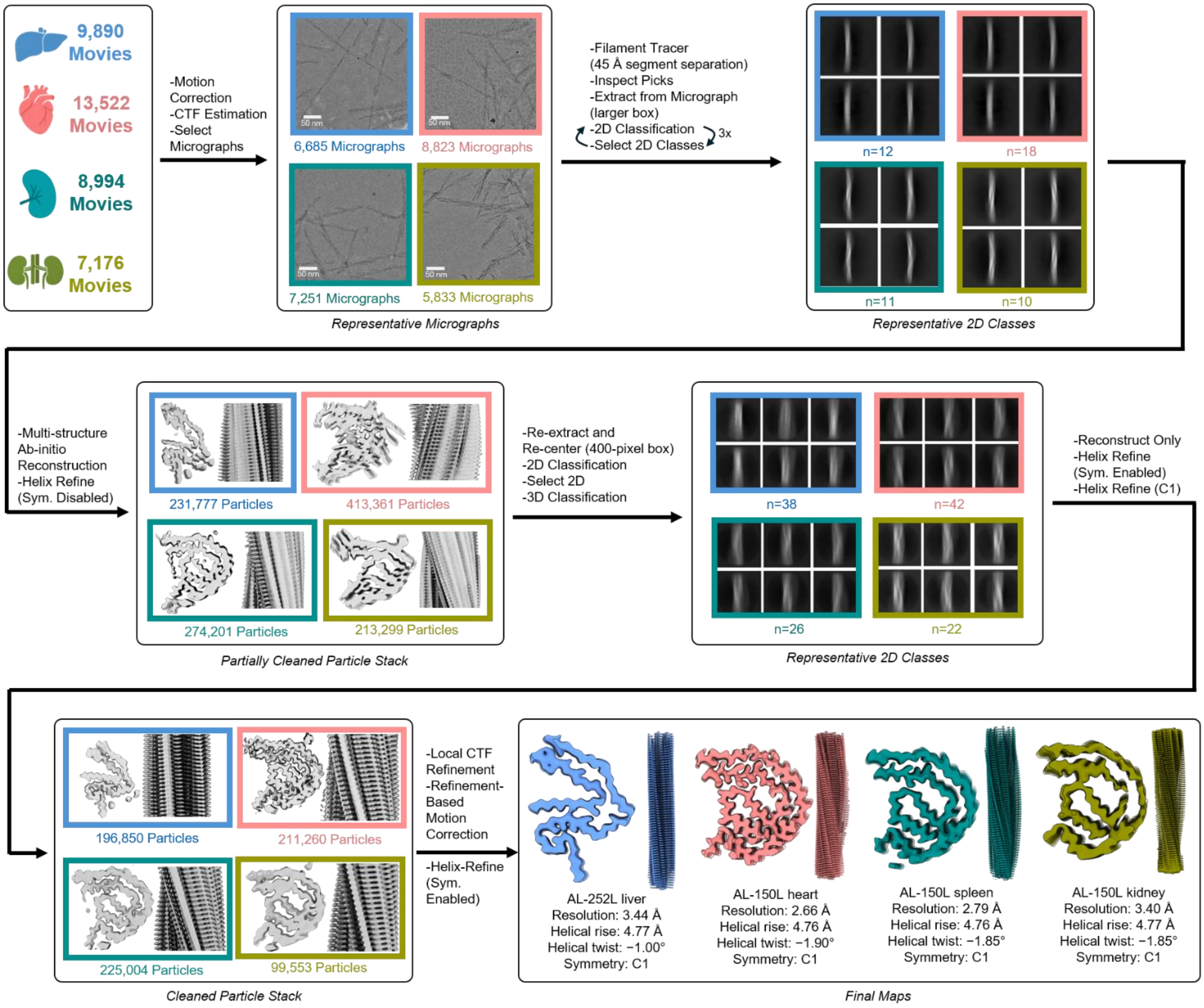


**Supplemental Figure 7**. **Cryo-EM data processing workflow**. Processing pipeline to generate the cryo-EM maps for the four amyloid structures determined in the current study.

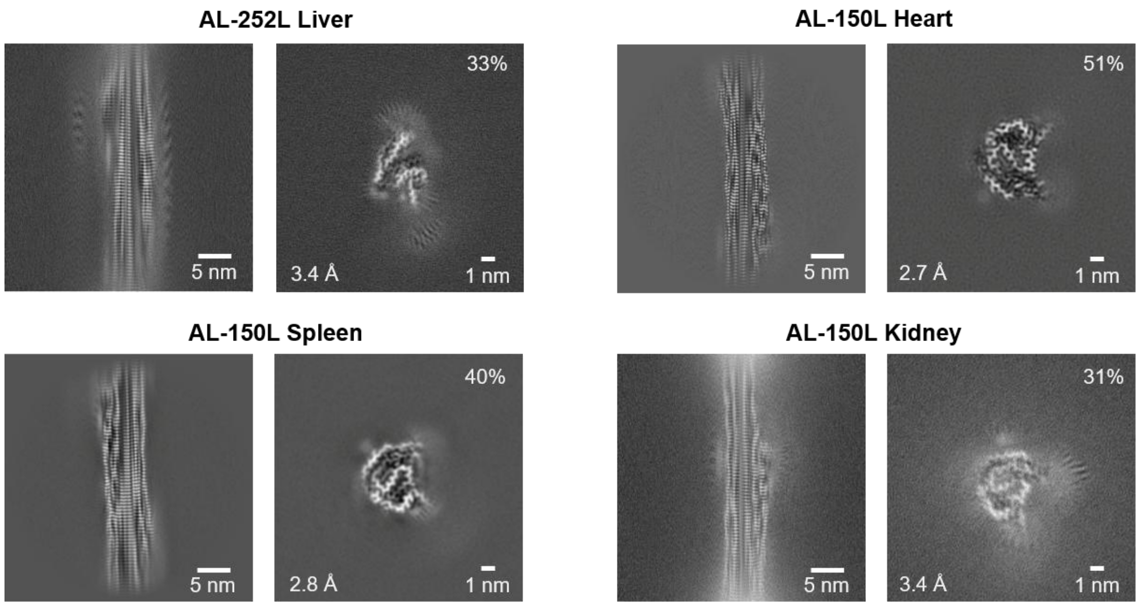


**Supplemental Figure 8**. **Representative 2D class averages and 3D reconstructions of tissue-derived amyloid fibrils**. Longitudinal 2D class averages and corresponding cross-sectional 3D reconstructions are shown for hepatic AL-252L, cardiac AL-150L, splenic AL-150L, and renal AL-150L amyloids. Percentages indicate the particle fraction assigned to each reconstructed class. The reported map resolutions are shown at the lower left of the cross-sectional panels.

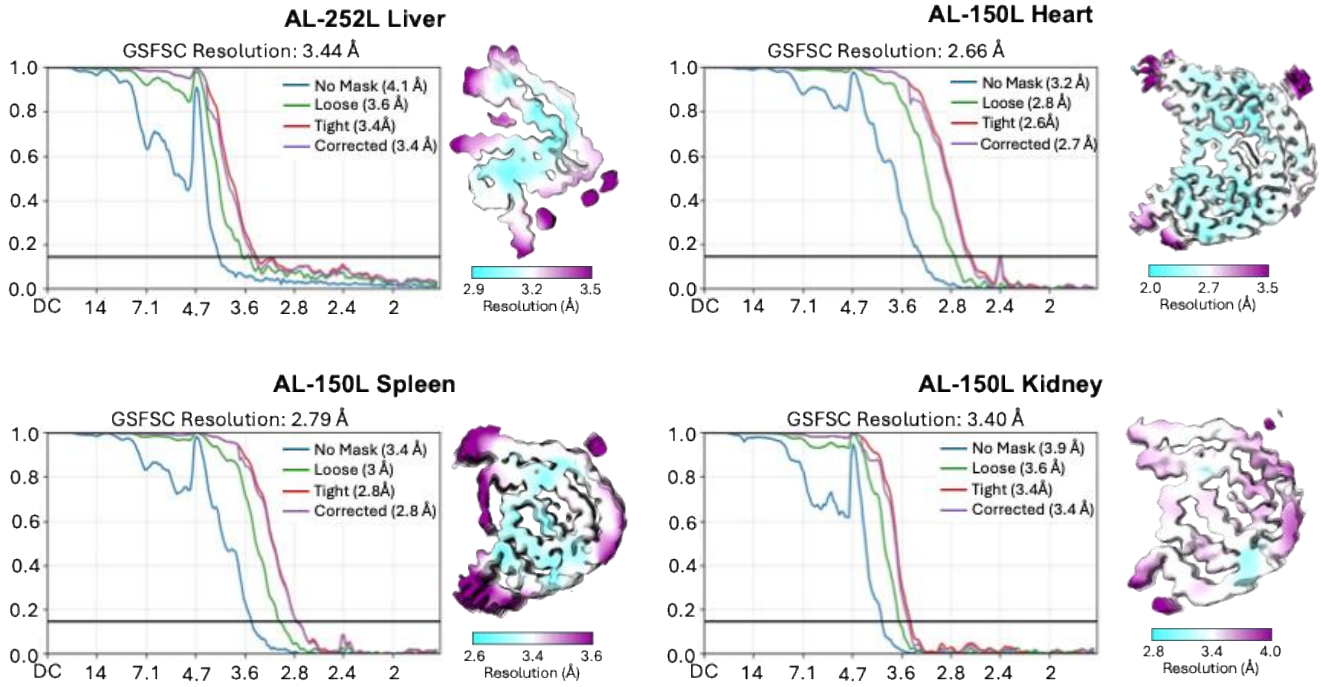


**Supplemental Figure 9**. **Global and local resolution of AL amyloid fibril reconstructions**. Gold-standard Fourier shell correlation (FSC) curves and cryo-EM maps colored according to local resolution are shown for hepatic AL-252L, cardiac AL-150L, splenic AL-150L, and renal AL-150L amyloids. The horizontal line marks the FSC = 0.143 criterion. Global resolution is indicated on each plot.

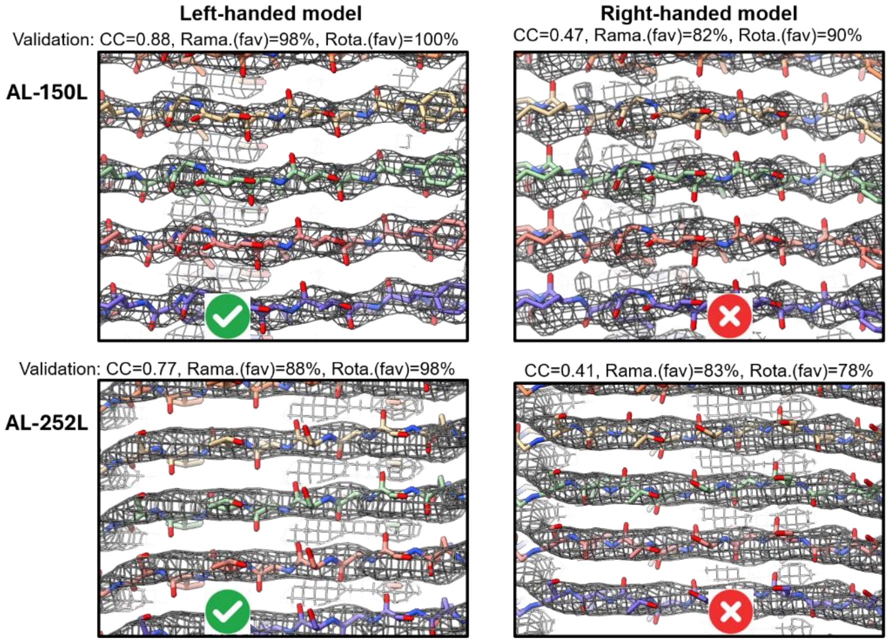


**Supplemental Figure 10**. **Helical handedness determination in amyloid fibrils**. Superimposition of the atomic models of AL-150L (top) and AL-252L (bottom) amyloids on the highly sharpened left- and right-handed cryo-EM maps (shown as mesh). The left-handed models show better alignment of the backbone carbonyl orientations with the corrugated map density, higher map–model correlation (CC), and more favorable Ramachandran (Rama) and rotamer (Rota) statistics. Validation values are indicated.

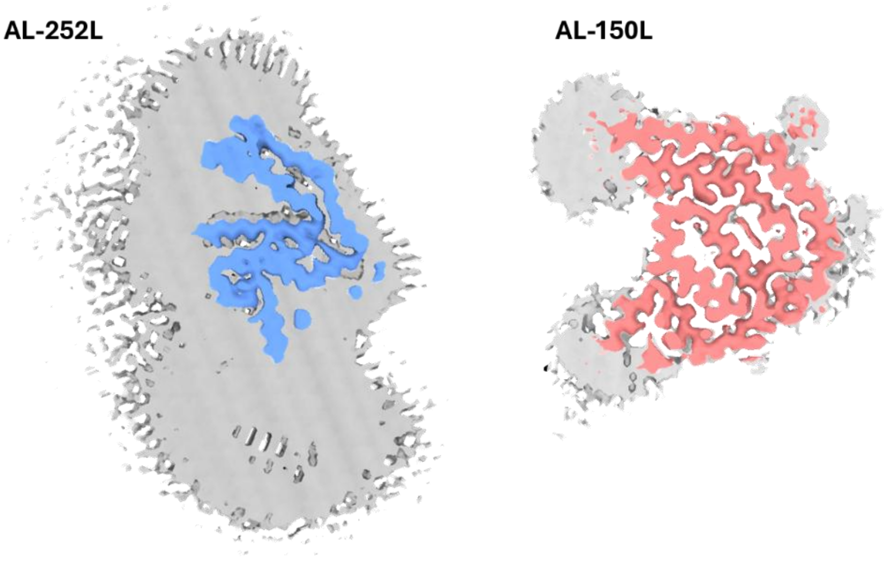


**Supplemental Figure 11**. **False discovery rate (FDR)-thresholded confidence maps of AL-252L and AL-150L amyloid fibrils.** Sharpened EM map for AL-252L amyloid (blue) is shown within the confidence map (gray) generated by FDR thresholding of the unsharpened input map. The confidence map was visualized at a threshold of 0.9999999, equivalent to FDR = 0.00001% for the displayed voxels. Sharpened EM map for AL-150L amyloid (salmon) is shown within the confidence map (gray) visualized at a threshold of 0.99999999, equivalent to FDR = 0.000001% for the displayed voxels.

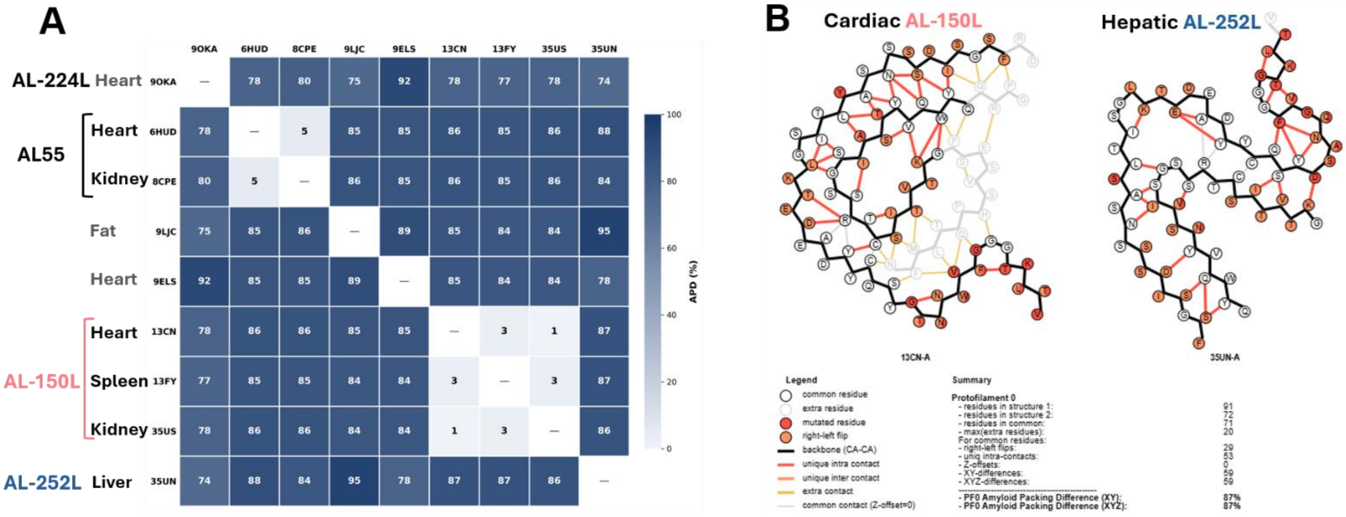


**Supplemental Figure 12 Pairwise comparisons of side chain packing in fibril structures of λ6-LCs.** **A.** Heat map shows amyloid packing differences (APDs) for the available structures of λ6-LC amyloid calculated using the APD metrics[40]. PDB IDs and available NCBI numbers are shown, other details are listed in Table 1. APDs are color-coded from 0% to 100% as shown. Structures of fibrils extracted from different organs of the same patient, which include AL55 cardiac and renal amyloids (PDB ID: 6HUD and 8CPE) and AL-150L cardiac, splenic and renal amyloids (PDB ID:13CN, 13FY and 35US), show APDs of 1-5%, indicating nearly identical amyloid core packing. Amyloid structures from different patients show APDs of 74-95%, indicating very different amyloid core packing despite shared features in the λ6-LC fibril folds, which are all encoded by the same germline. **B.** Bead diagrams compare side chain contacts in cardiac AL150L and hepatic AL252L amyloid structures. Thick lines show well-ordered shared (black) and structure-specific (gray) backbone. Orange circles – flipped side chains. Common contacts are in thin gray lines; unique intra-chain contacts are in red lines. Other details are listed in the panel.

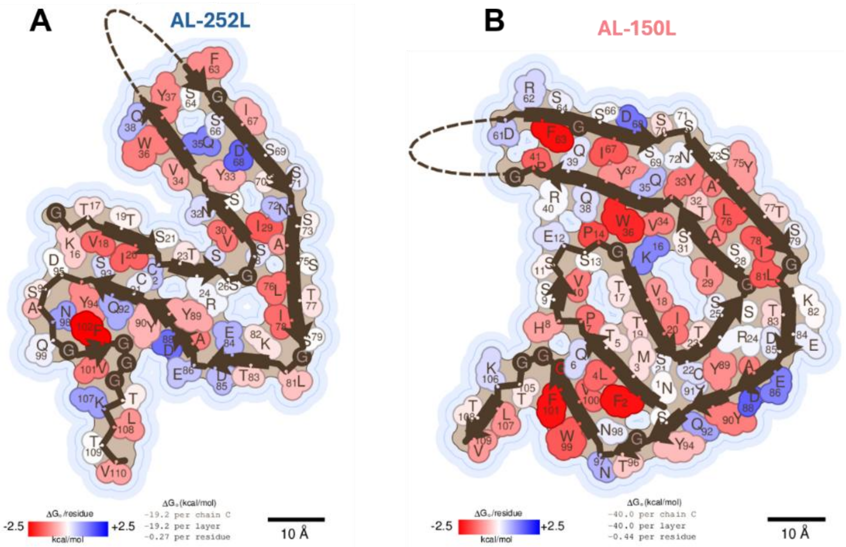

**Supplemental Figure 13. Side chain packing contributions to amyloid core stability in AL-252L and AL-150L fibrils.** Side chain packing in amyloid cores of hepatic AL-252L **(A)** and cardiac AL-150L(**B**) fibrils. Energy map shows favorable (ΔG<0, red) and unfavorable (ΔG>0, blue) contributions to the amyloid core stability. The diagrams were generated using Amyloid Illustrator server, available at https://zenodo.org/records/15218932.

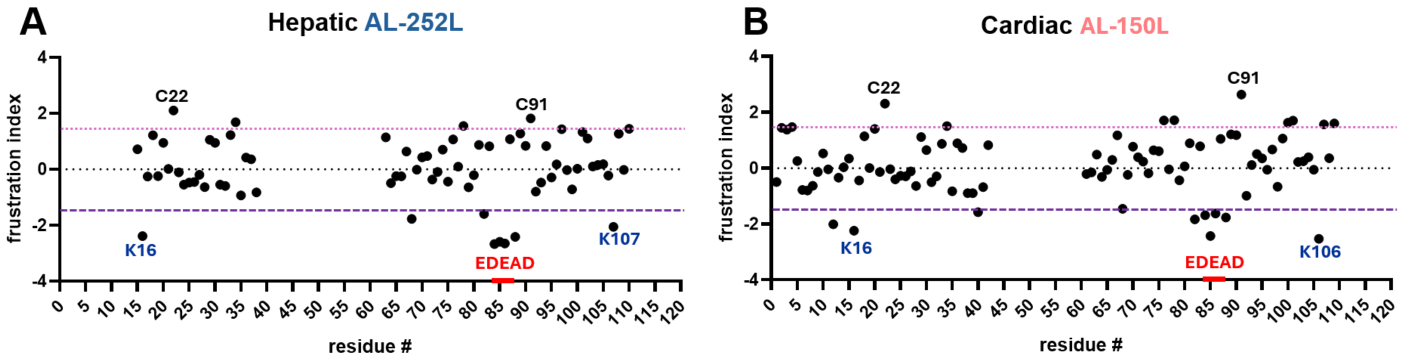


**Supplemental Figure 14. Frustration indices of individual sidec hains in amyloid cores.**

Frustration indices as a function of side chain position were computed for amyloid structures of hepatic AL-252L **(A)** and cardiac AL-150L **(B)** using the Frustratometer server[48] available at http://frustratometer.qb.fcen.uba.ar Values below -1.5 (purple dashed line, lowest) indicate high structural frustration. Values above +1.5 (pink dotted line, highest) indicate minimal frustration. Representative side chains with minimal frustration (disulfide-forming cysteines C22 and C91) and with high frustration (acidic residues from the E84-D88 segment, EDEAD, and uncompensated charges on K16 and K106/107) are marked. These ionizable side chains show similar trends in other amyloid structures of λ6-LCs.

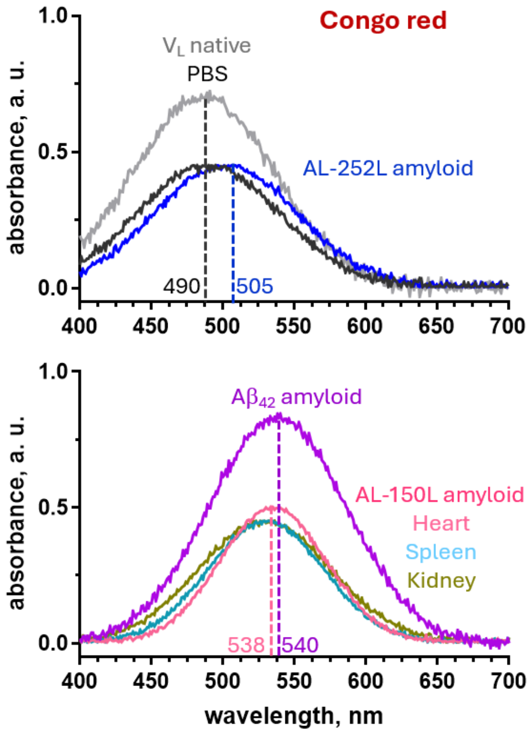


**Supplemental Figure 15. Absorption spectra of the amyloid diagnostic dye, Congo red, indicate differential dye binding to AL-252L and AL-150L amyloids.** Congo red binding to amyloid induces a red shift in the absorption peak, from 490 nm of the dye in buffer at pH~7 to >500 nm for the dye bound to amyloid. Top panel: Absorption spectra of Congo red with AL-252L amyloid show a small red shift from 490 nm to 505 nm, suggesting weak binding (blue). Bottom panel: A much larger red shift to 538 nm is observed in the presence of cardiac, splenic or renal AL-150L amyloids, indicating much stronger binding (pink). Spectra of the dye in buffer alone (PBS, black) or in the presence of natively folded $V_L$ (grey) provide negative controls; spectrum in the presence of $A\beta_{42}$ amyloid (violet) provides a positive control.

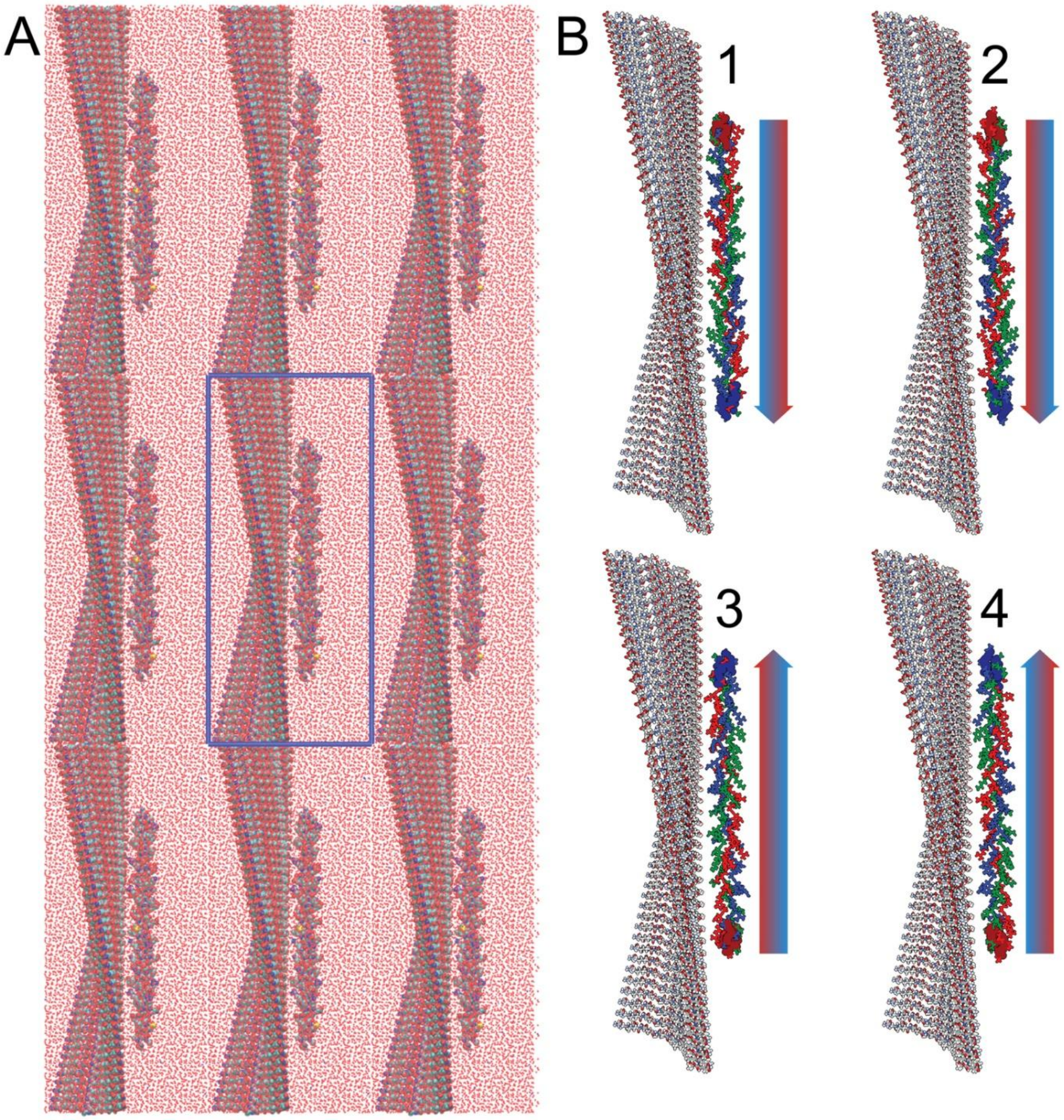


**Supplemental Figure 16**. **Initial configuration of the system for molecular dynamics simulations of AL150L amyloid in complex with a triple-helical ColVI fragment**.

**A.** The 9 nm x 7 nm x 20.3 nm box contains ~124,600 atoms including AL-150L residue segment S64-E84 in amyloid conformation, a representative fragment of ColVI triple helix, and solvent. Blue lines demarcate the periodic boundary condition. Atoms are color-coded N (blue), O (red), C (teal), and H (white). **B**. Initial relative orientations 1, 2, 3, and 4 of a triple-helical fragment in respect to amyloid surface, each differing by 180° rotations about the z-axis and y-axis. Leading, middle, and trailing strands of ColVI fragment are colored in blue, green, and red, respectively. Backbone atoms of three N-terminal and three C-terminal amino acid atoms are displayed as spheres in dark blue and dark red, respectively. Gradient arrows illustrate the triple-helical orientation in respect to amyloid.

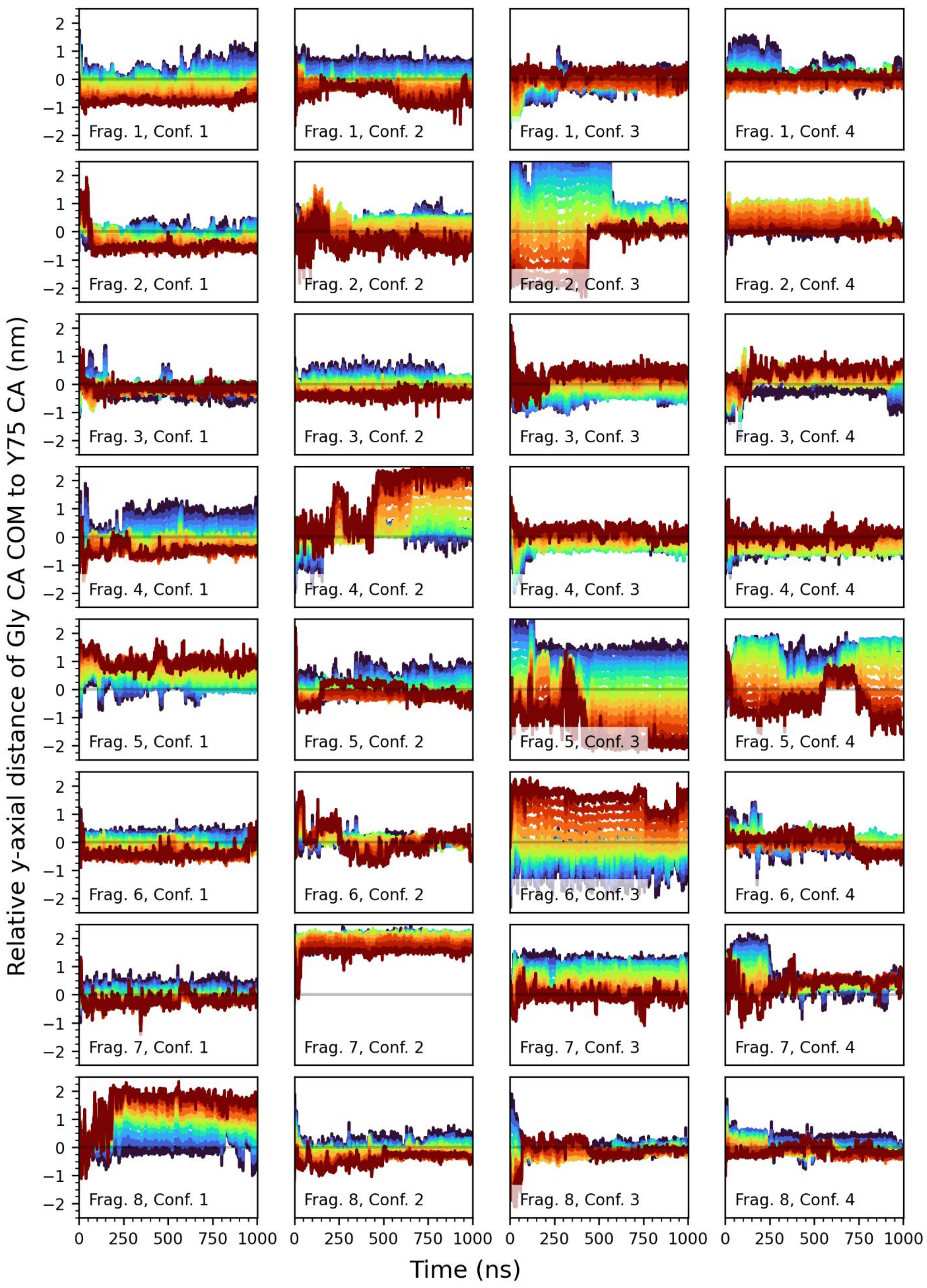


**Supplemental Figure 17**. **Exploration of the amyloid surface by ColVI triple-helical fragments during 1 μs MD simulations**. MD trajectories of each fragment (Frag. 1-8 described in supplemental Methods) in each of the four initial configurations (Conf. 1-4, supplemental Fig. 16B) show time-dependent y-axial distance for the center of mass of every third Gly $C_\alpha$ in the ColVI triple helix to the closest Y75 $C_\alpha$ in amyloid. GXY triplets in each fragment are rainbow-colored from the N-terminal (dark blue) to the C-terminal (dark red).

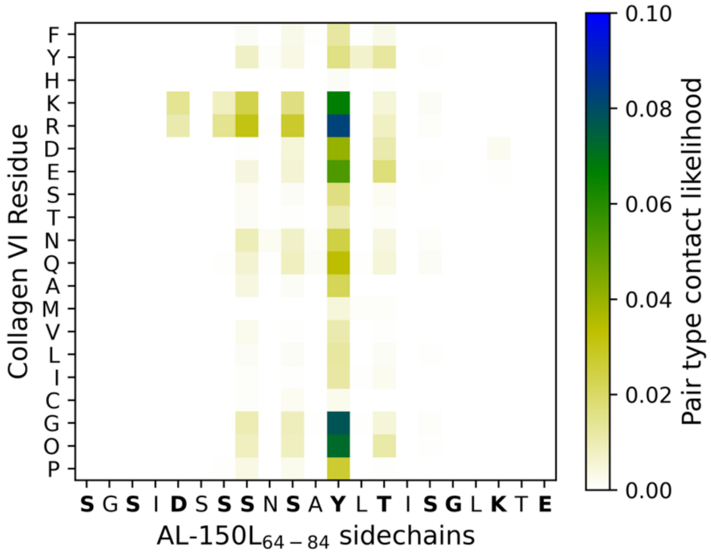


**Supplemental Figure 18**. **Probability of contacts between ColVI and AL-150L amyloid side chains determined in molecular dynamics simulations**. Contacts within a 3.5 Å cutoff, $p_{i,a}$, between non-hydrogen atoms of ColVI and amyloid are plotted by residue type. ColVI triple helix contains a high fraction of charged residues (K, R, D, E), Gly, and Hyp (O). MD simulations suggest these residues are principally responsible for interactions with amyloid. Out of six solvent-facing serines in S64-E84 segment, only S64 interacts with the triple helices in MD simulations.

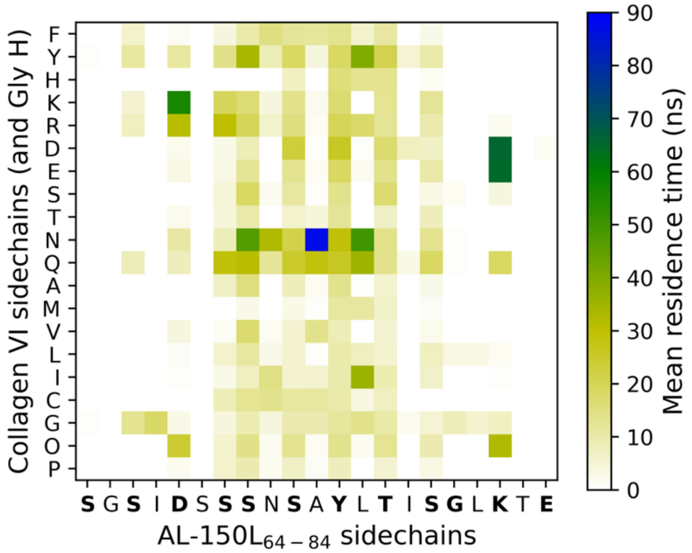


**Supplemental Figure 19. Average contact lifetimes between ColVI triple-helical fragments and AL-150L amyloid during molecular dynamics simulations**. Contacts between heavy atoms of ColVI and amyloid are plotted by residue type using a 3.5 Å cutoff and a 5 ns grace period. Charge-charge interactions and hydrogen bonding of ColVI Tyr to the amyloid backbone between Y75 and T77 are the longest-lived interactions but are not the principal determinants of ColVI-amyloid interactions. Out of six solvent-facing serines in S64-E84 segment, only S64 interacts with the triple helices in MD simulations.

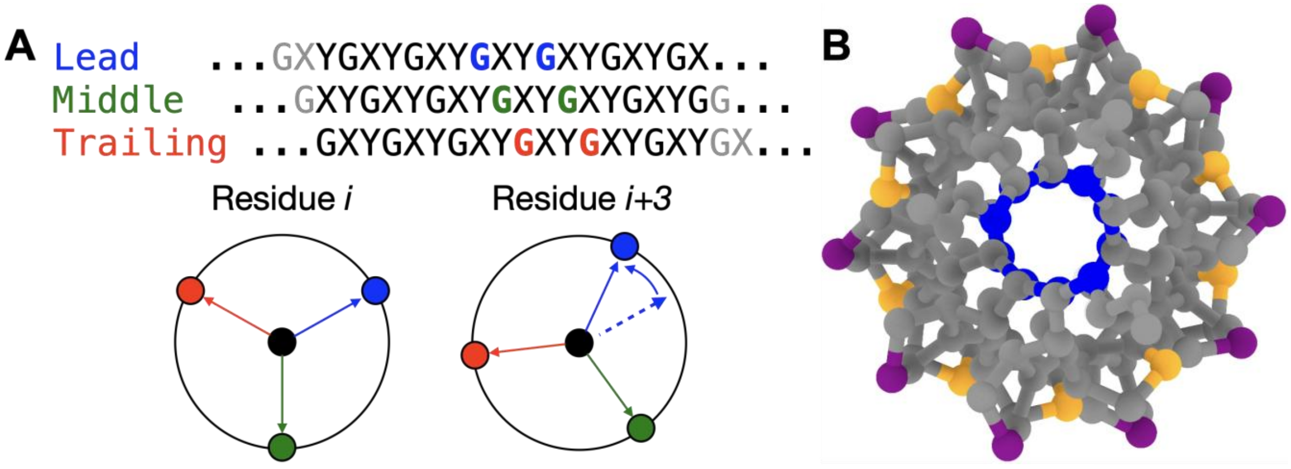


**Supplemental Figure 20. Schematic illustration of atoms defining the cylindrical coordinate system used to measure helical twist for leading, middle, and trailing chains of a triple helix**.

**A.** The local cylindrical coordinate system is defined using 18 residues that are relatively flush across the three chains, offset by 1 residue each. Gray residues are not used for defining the coordinate system. **B.** *X* (orange) and *Y* (purple) Cγ defining 20 unique angular positions separated by 18° in an idealized 10/3 helix model of a triple helix comprised of Pro-Hyp-Gly repeats. Gly Cα is in blue.

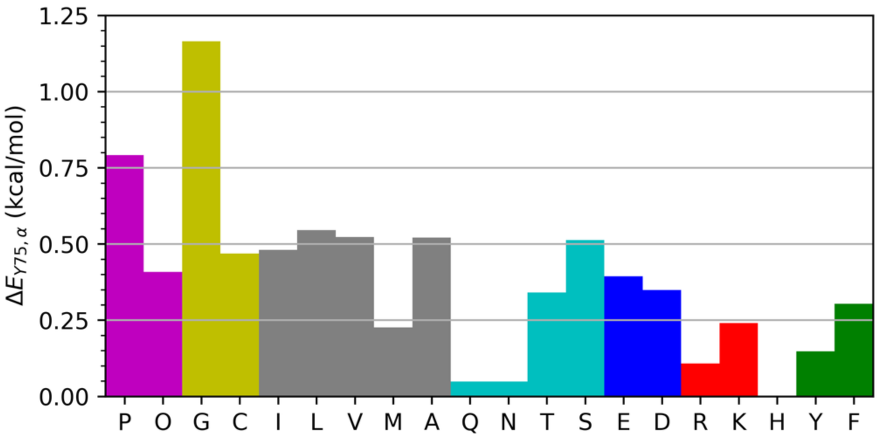


**Supplemental Figure 21. Relative free energy of interaction of each type of residue at *X*- or *Y*-positions in the ColVI triple helix with Y75 of amyloid, $\Delta E_{Y75,\alpha}$.**

The energies are shown compared to Y of ColVI. Positive values mean less favorable interactions than those between Y of ColVI and Y75 of amyloid.

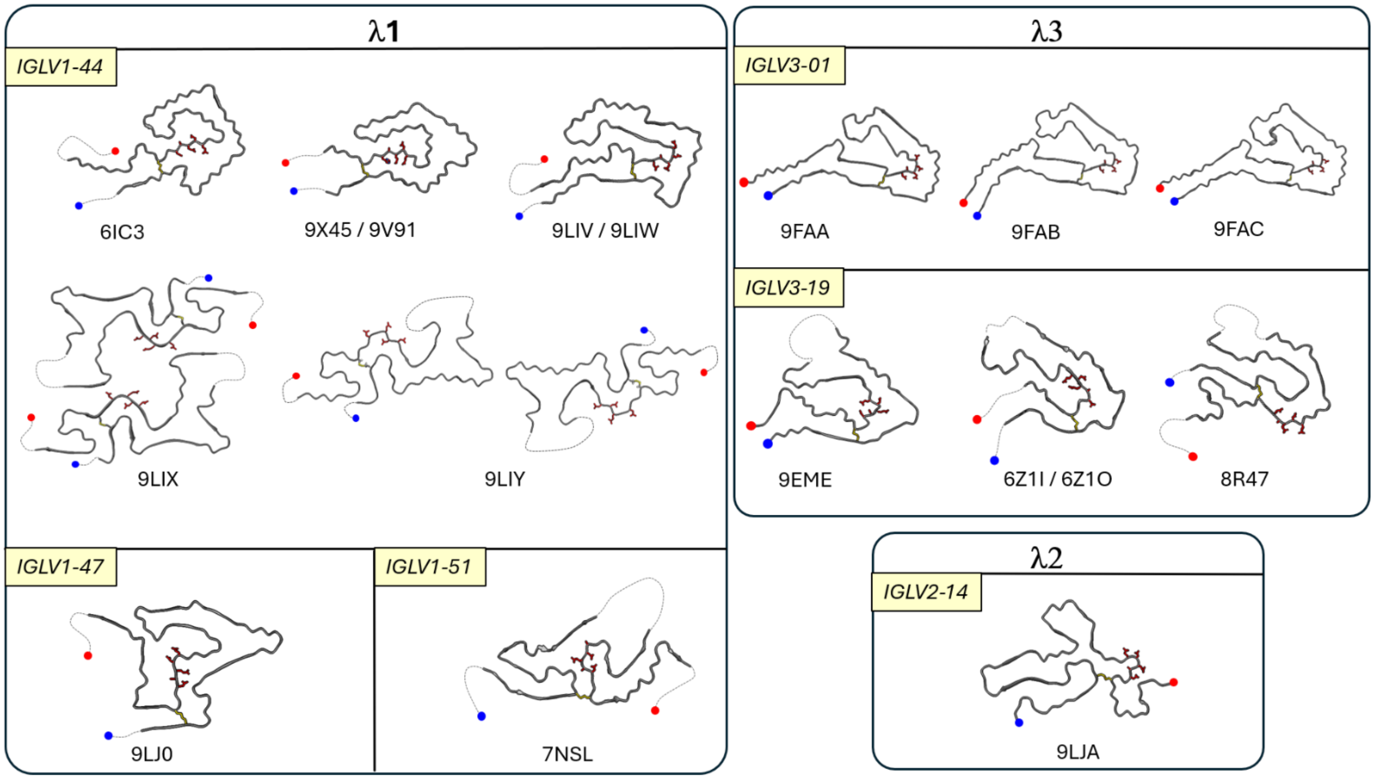


**Supplemental Figure 22. Representative cryo-EM structures of human tissue-extracted AL fibrils from λ1, λ2 and λ3 protein families**. PDB ID for each structure is indicated. $V_L$-encoding germlines are in italics. Backbone conformations are shown for one molecule in a fibril layer. Poorly ordered segments are in dotted lines. Blue and red circles mark N- and C-termini of the fibril-forming protein. Stick models show residues in the acidic-rich segment. While this segment is fully exposed on the fibril surface in PDB ID 8R47 and is partially exposed in PDB ID 9LIY and 9LJA, it is sequestered in all other structures from λ1-LC and λ3-LC families. In contrast, in all known AL amyloid structures of λ6-LCs, the acidic-rich EDEAD segment is exposed (Fig. 8).
